# Genome-wide analysis of social behaviour in context: a meta-regression approach across social domains, reporters and developmental stages

**DOI:** 10.1101/2025.08.16.667148

**Authors:** Lucía de Hoyos, Fenja Schlag, Sanjeevan Jahagirdar, Elizabeth C. Corfield, Andrea G. Allegrini, Danielle A. G. Admiraal, Eveline L. de Zeeuw, Ilja M. Nolte, Natalia Llonga, Alexander Neumann, Katherine Lange, Simone van den Bedem, Ebba Du Rietz, Ehsan Motazedi, Else Eising, Nicole Y. T. Ng, Teemu Palviainen, Carol A. Wang, Elisabeth Thiering, Sofia Scatolin, Priyanka Choudhary, Natalia Vilor-Tejedor, Zhijie Liao, Silvia Alemany, Casper-Emil T. Pedersen, Nora Fernandez-Jimenez, Megan L. Campbell, Polina Girchenko, Eli Barthome, Hanna Seelemeyer, Afsheen Kumar, Alexandre Jeanne, Tarunveer S. Ahluwalia, Melissa H. Black, Jan K. Buitelaar, Simon E. Fisher, Anni Heiskala, María Hernández-Lorca, Vincent W. Jaddoe, Marjo-Riitta Järvelin, Sheri-Michelle Koopowitz, Marilyn T. Lake, Paul Lichtenstein, Sabrina Llop, Mannan Luo, Anni Malmberg, Sergi Marí, Albertine J. Oldehinkel, Zdenka Pausova, Craig E. Pennell, Robert Plomin, Ted Reichborn-Kjennerud, Sheena Reilly, Alina Rodriguez, Thomas Rolland, Angelica Ronald, Katri Räikkönen, Jean Shin, María Soler Artigas, Ashley E. Tate, Henning Tiemeier, Ellen Verhoef, Eero Vuoksimaa, Melissa Wake, Varun Warrier, Eivind Ystrom, Sven Bölte, Christine Ecker, Thomas Bourgeron, Jari Lahti, Dan J. Stein, Jose Ramon Bilbao, Klaus Bønnelykke, Mariona Bustamante, Tomas Paus, Christel M. Middeldorp, Jordi Sunyer, Sylvain Sebert, Marie Standl, Andrew J. O. Whitehouse, Jaakko Kaprio, J. Antoni Ramos-Quiroga, Marta Ribases, Miguel Casas, Rosa Bosch, Angela T. Morgan, Tanja G. M. Vrijkotte, Henrik Larsson, Charlotte A. M. Cecil, Catharina A. Hartman, Dorret I. Boomsma, Kaili Rimfeld, Alexandra Havdahl, Beate St Pourcain

## Abstract

Social behaviour is a heritable, context-dependent trait that changes across social settings and development, influencing wellbeing and mental health. We present the first genome-wide meta-regression study of social behaviour from infancy to early adulthood, leveraging 491,246 repeat measures of low prosocial behaviour and peer/social difficulties in European-ancestry cohorts (N_eff_=121,777, N_ind_=73,321). We modelled heterogeneity in genetic effects across social domains, informants, and ages (2–29 years), capturing social context through genomic influences. Six loci were identified, including variation within *CADM2* (*p*=2.51x10^-9^). The SNP-based heritability was modest (2–7%), and the genetic architecture of social behaviour multidimensional. Polygenic scores demonstrated predictability and accuracy in independent European-ancestry cohorts and, partially, in African-ancestry cohorts (N_ind_=16,305). Genetic correlations with later-life and mental health outcomes showed context-dependent patterns. Modelling predicted onsets of associations with social behaviour revealed distinct profiles, as observed for autism, ADHD, depression and schizophrenia, highlighting novel opportunities to genetically proxy developmental trajectories.

## INTRODUCTION

Human social behaviour encompasses a broad spectrum of inter-individual interactions that change across development and vary across social situations^1,2^. Prosocial behaviour involves voluntary actions that benefit others, such as helping, sharing, and comforting. Early forms of prosocial behaviour can be observed from the age of two years^3^, and mastering increasingly complex levels of social competence during development shapes social roles in society^2^. Social behaviour also includes peer and social difficulties, such as challenges to conform to developmental expectations, poor social skills, difficulties developing friendships and peer rejection^4,5^. In particular, childhood peer difficulties have been associated with lower academic performance, self-esteem, social engagement, problem-solving skills, and resilience^6,7^. Thus, early social behaviour plays an important part in later-life friendships, wellbeing and (mental) health^6–8^.

Twin studies have shown that both prosocial behaviour and peer and social problems during childhood and adolescence are heritable, with twin heritability (h^2^_twin_) estimates of 26-73%^9,10^ and 24-71%^11–13^, respectively. Corresponding single-nucleotide polymorphism (SNP)-based heritability (h^2^_SNP_) estimates for peer and social problems ranged from 2-27%^11,14^. Yet, genetic influences vary by age and informant (e.g. parent vs teacher reports)^9–11,15^, and changes in genetic effects may reflect variation in developmental processes driving social behaviour in different social situations.

Here, we present a genome-wide meta-regression analysis, a versatile framework integrating trait characteristics as moderators into genetic association analyses^16^, to model heterogeneity in genetic effects underlying social behaviour from infancy to young adulthood. We examine SNP effects and the genetic architecture of social behaviour, accounting for variation in social domain, informant and age. We evaluate the predictive ability and accuracy of polygenic scores (PGS) of social behaviour and report genetic correlates, including population-based traits and neurodevelopmental and neuropsychiatric conditions (NNCs). Using genetically-inferred developmental differences, we estimate the predicted onset of genetic associations with social behaviour, proxying developmental trajectories for later life outcomes.

## RESULTS

### Modelling heterogeneity through genome-wide meta-regression analysis

Within the EArly Genetics and Lifecourse Epidemiology (EAGLE) Consortium^17^ (https://www.eagle-consortium.org), we conducted a genome-wide association study (GWAS) meta-analysis combining 195 GWAS summary statistics from 73,321 individuals of European ancestry comprising 491,246 repeat observations (N_eff_=121,777) across 25 independent cohorts (Figure 1a-c, Supplementary Tables 1-2, Supplementary Note 1). Cohort-level meta information included (1) social domain, assessing two domains of social difficulties (LPB: reverse-coded low prosocial behaviour; PSP: peer and social problems), (2) informants (parent-, teacher-, and self-reports, Supplementary Table 2; hereafter referred to as reporter) and (3) age spanning 2-29 years (Figure 1c). Social behaviour was predominantly measured with standardised instruments (Supplementary Table 3), such as the Strengths and Difficulties Questionnaire (SDQ)^4^ or the Child Behaviour Checklist (CBCL)^18^. To map psychometric instruments onto a social behavioural domain, each questionnaire item was aligned to a category of the WHO’s International Classification of Functioning, Disability and Health (ICF, Children & Youth Version) (Figure 1b, Supplementary Table 4, Supplementary Figure 1, Supplementary Note 2, Methods), a biopsychosocial framework and classification system for describing and organising information on human functioning and disability^19^. For family-based cohorts, relatedness was accounted for by each cohort-level GWAS.

**Figure 1.**
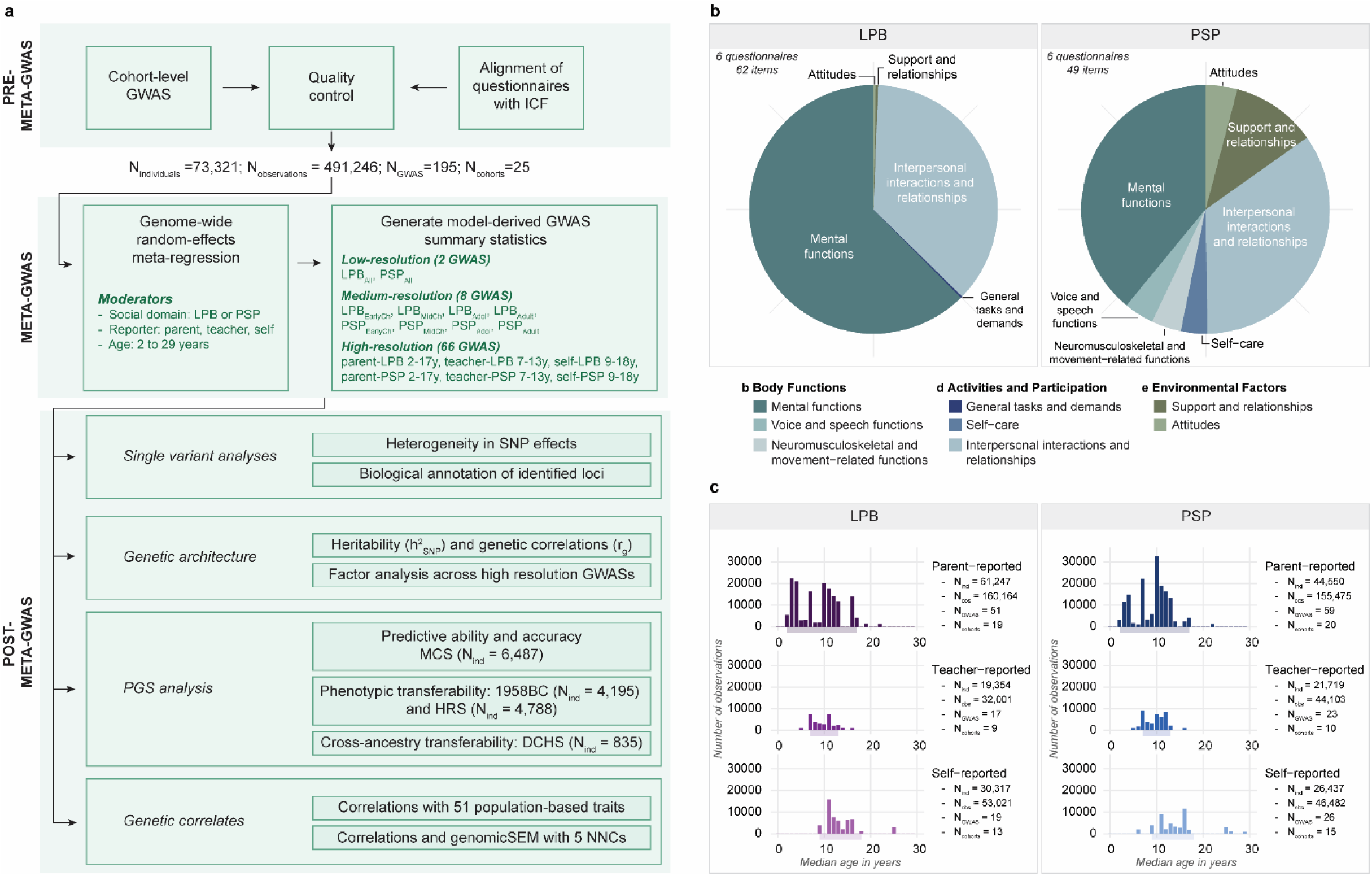
Social behaviour meta-GWAS design. (**a**) Analysis design including a meta-regression across two social domains, three reporters and ages. (**b**) Distribution of ICF codes per social domain. (**c**) Distribution of the number of individuals across age, per social domain and reporter, in addition to the exact number of observations (N_obs_), the number of individuals (N_ind_), GWAS statistics (N_GWAS_) and cohorts (N_cohorts_). Abbreviations: 1958BC (1958 birth cohort), DCHS (Drakenstein Child Health Study), HRS (Health and Retirement Study), ICF (WHO’s International Classification of Functioning, Disability and Health, Children & Youth Version), LPB (Low prosocial behaviour), MCS (Millennium Cohort Study), NNCs (Neurodevelopmental and neuropsychiatric conditions), PSP (Peer and social problems)

To model genetic associations allowing for effect moderation, we implemented a genome-wide random-effects meta-regression approach (meta-GWAS, Figure 1a) using the metafor R package^20^. For each SNP, fixed effects (θ parameters) captured systematic changes in the genetic effect due to differences in social domain (reference: LPB), reporter (reference: parent) and (centered grand mean) age, across the 195 GWAS statistics, while accounting for repeat observations, with random effects nested within each cohort (Methods). Thus, the fixed-effect intercept represented the parent-reported LPB SNP effect at the grand mean age.

From each SNP-based model, genome-wide, we derived three groups of GWAS summary statistics with increasingly detailed levels of moderator information (Supplementary Table 5, Methods): (i) low-resolution GWAS differentiating between social domains (2 GWAS: LPB_All_ and PSP_All_), (ii) medium-resolution GWAS differentiating between social domains across broad developmental stages, i.e. early childhood (2-5 years), middle childhood (6-11 years), adolescence (12-17 years) and young adulthood (18-29 years) (8 GWAS: LPB_EarlyCh_, LPB_MidCh_, LPB_Adol_, LPB_Adult_, PSP_EarlyCh_, PSP_MidCh_, PSP_Adol_, and PSP_Adult_) and (iii) high-resolution GWAS differentiating between social domains across different reporters and individual ages (66 GWAS). Deriving GWAS at high resolution, based on a single multivariate model per SNP, allowed capturing systematic and dynamic changes in genetic effect sizes across the genome, as visualised in dynamic Manhattan plots (Supplementary Videos 1-6). Thus, a meta-regression approach facilitates the harmonisation of summary statistics across diverse phenotypes into interpretable patterns while controlling effectively for multiple testing.

Across the 76 derived GWAS, we identified six independent genome-wide associated (GWA) loci (p<5x10^-8^; with one lead variant per locus (lead SNPs: rs148774891, rs181817841, rs7490422, rs77125329, rs538645, and rs150685860), as confirmed in conditional and joint analyses using GCTA v1.94^21^ software (Methods (COJO), Table 1, Supplementary Table 6, Supplementary Figure 2-Supplementary Figure 4, Supplementary Note 3, Supplementary Videos 1-6). All six loci exhibited considerable heterogeneity in genetic effects with moderating (θ) effects attributable to differences in social domain, reporter, or age (Supplementary Table 7, Supplementary Note 3).

**Table 1.** Genome-wide associated loci for social behaviour across social domain, reporter, and age.

| Lead SNP | Position (hg19) | EA | EAf | $\beta$(SE) | Strongest association (p-value) | GWAS where hit is reported | Nearest gene | max pLI (gene) | max CADD (gene) |
| --- | --- | --- | --- | --- | --- | --- | --- | --- | --- |
| rs148774891 | chr3:86,059,243 | A | 1.29 % | 0.202 (0.034) | Self-reported LPB 9 years (p=2.51x10<sup>-9</sup>) | Self-reported LPB 9-14 years | *CADM2*<br>*CADM2-AS1* | 0.98<br>(*CADM2*) | 3.14<br>(*CADM2*) |
| rs181817841 | chr8:78,481,590 | T | 1.25 % | 0.154 (0.026) | Parent-reported LPB 6 years (p=3.33x10<sup>-9</sup>) | Parent-reported LPB 3-8 years | *RP11-38H17.1* | - | 5.12<br>(*RP11-38H17.1*) |
| rs7490422 | chr13:23,284,225 | G | 35.04 % | -0.035 (0.006) | Parent-reported PSP 6 years (p=5.37x10<sup>-9</sup>) | Parent-reported PSP 4-8 years | *DDX39AP1* | - | 1.68<br>(*DDX39AP1*) |
| rs77125329 | chr10:78,297,483 | T | 1.24 % | -0.150 (0.026) | Parent-reported LPB 4 years (p=5.70x10<sup>-9</sup>) | Parent-reported LPB 2-5 years and LPB<sub>EarlyCh</sub> | *LRMDA* | 0.04<br>(*LRMDA*) | 3.81<br>(*LRMDA*) |
| rs538645 | chr11:118,711,069 | T | 76.43 % | -0.044 (0.008) | PSP<sub>EarlyCh</sub> (p=1.41x10<sup>-8</sup>) | PSP<sub>EarlyCh</sub> | *SETP16*<br>*Y_RNA* | 1.00<br>(*ARCN1*) | 14.02<br>(*SETP16* & *Y_RNA*) |
| rs150685860 | chr20:30,561,451 | A | 2.09 % | -0.164 (0.030) | Parent-reported PSP 2 years (p=2.72x10<sup>-8</sup>) | Parent-reported PSP 2-3 years | *XKR7* | 0.92<br>(*XKR7*) | 6.35<br>(*XKR7*) |

### Biological annotation of single-variant signals

As part of post-meta-GWAS analyses (Figure 1a; Supplementary Tables 8-11, Supplementary Figures 5-15, Supplementary Note 4, Methods), the six GWA loci were mapped to 18 genes (8 protein-coding) based on positional and expression quantitative loci (eQTL) mapping using FUMA^22^ (Supplementary Table 8) and to another 36 genes (8 protein-coding genes) through chromatin interaction mapping (Supplementary Table 9). Here, we focus on three GWA loci (rs148774891, rs150685860, and rs538645) with likely biological impact in detail (Figure 2).

**Figure 2.**
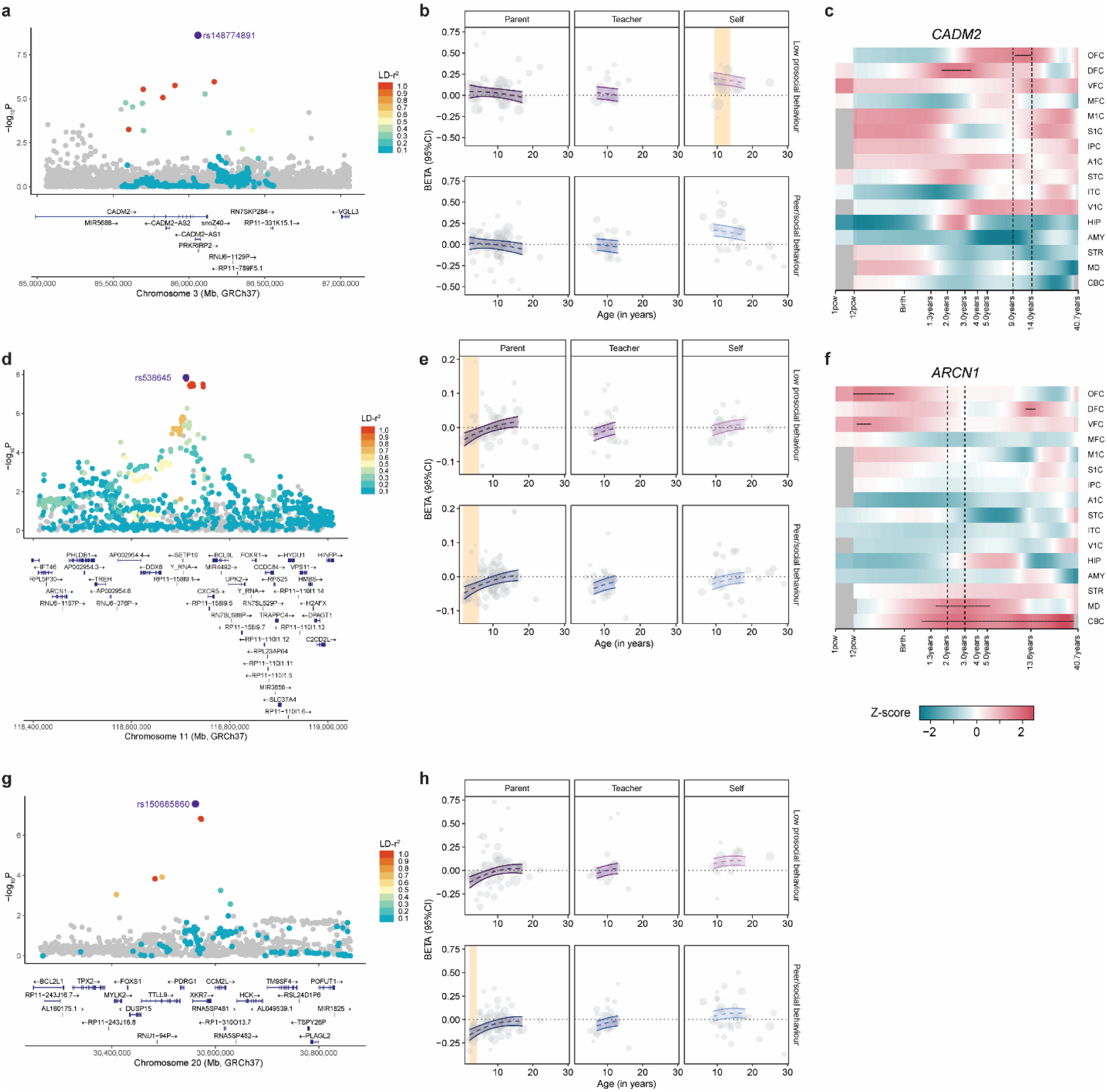
Identified genetic loci with potential biological implications. (**a**) Locus plot and (**b**) model-derived genetic effect for rs148774891. (**c**) Brain expression patterns of *CADM2* (mapped to rs148774891). (**d**) Locus plot and (**e**) model-derived genetic effects for rs150685860. (**f**) Brain expression patterns of *ARCN1* (mapped to rs150685860). (**g**) Locus plot and (**h**) model-derived genetic effect of rs538645. (**b,e,h**) The yellow shade marks social domain-, informant- and age-related context of each locus. (**c,f**) Age-related brain expression patterns were studied in BrainSpan Atlas microarray data (probes for XKR7 were not available). Overexpression is marked with a horizontal black line and the age-related window of the associated trait with vertical dashed lines. Abbreviations: A1C (Primary auditory (A1) cortex), AMY (Amygdala), CBC (Cerebellar cortex), DFC (Dorsolateral prefrontal cortex), HIP (Hippocampus), IPC (Posterior inferior parietal cortex), ITC (Inferior temporal cortex), M1C (Primary motor (M1) cortex), MD (Mediodorsal nucleus of the thalamus), MFC (Medial prefrontal cortex), OFC (Orbital prefrontal cortex), S1C (Primary somatosensory (S1) cortex), STC (Superior temporal cortex), STR (Striatum), V1C (Primary visual (V1) cortex), VFC (Ventrolateral prefrontal cortex)

The strongest signal, rs148774891, is a low-frequency variant (EAF=1.29%, A allele, Figure 2a) located on chromosome 3p12.1 within an intron of the Cell Adhesion Molecule 2 (*CADM2*) gene and *CADM2* Antisense RNA 1 gene (*CADM2-AS1*). Each effect allele increased self-reported LPB at 9 years by 0.20 standard deviation (SD) units (SE=0.034, p=2.5x10^-9^), but the signal remained detectable until age 14 (Table 1). *CADM2,* widely expressed in the brain, has a high score for probability loss-of-function intolerance (pLI=0.98) and plays a vital role in synapse organisation^23^ (Supplementary Figures 6 and 12). Matching the developmental window of the associated trait, *CADM2* over-expression peaks in the orbitofrontal cortex at 9-14 years (Figure 2c, Supplementary Figure 13), a region important for decision-making in social and reward-based contexts^24^.

The lead variant rs538645 (EAF=76.43%, T allele) on chromosome 11q23.3 was associated with a 0.044 SD decrease in early childhood PSP (ages 2-5 years; SE=0.0077, p=1.4x10^-8^, Figure 2e). The locus was mapped to two genes: Within the SET Pseudogene 16 (*SETP16*, ENSG00000201535: *Y RNA,* Supplementary Table 8), the variant has a Combined Annotation–Dependent Depletion (CADD) score of 14.02, indicating potential deleteriousness. Via eQTL mapping, the locus was linked to Archain 1 (*ARCN1,* p_eQTL_=2.67x10^-7^, rs7104519:LD-r^2^=1, Supplementary Table 8), a high-pLI score gene (pLI=1) expressed in the brain (Supplementary Figure 10). Matching the developmental stage of the associated trait, *ARCN1* mRNA showed above-average expression in the cerebellar cortex and the mediodorsal nucleus of the thalamus across ages of 1 to 5 years (Figure 2f, Supplementary Figure 13), where the latter is a control hub influencing social behaviour via thalamocortical circuits^25^.

Lastly, the low-frequency variant rs150685860 (EAF=2.09%, A allele) resides within an intron of the XK-related 7 gene (*XKR7*) on chromosome 20q11.21 (Figure 2d), a high pLI score gene (pLI=0.92) involved in apoptotic processes. *XKR7* is widely expressed in the Cerebellum, but cortical and subcortical expression was low (Supplementary Figure 11). Each effect allele was associated with a 0.16 SD decrease in parent-reported PSP at 2 years (SE=0.029, p=2.7x10^-8^), indicating a rare protective effect limited to ages 2 to 3 years (Table 1).

### The genetic architecture of social behaviour is multidimensional

Social behaviour was modestly heritable (h^2^_SNP_ =2-7%, Figure 3a, Supplementary Table 12), but GWAS were overall sufficiently powered, as estimated with linkage disequilibrium score (LDSC) regressions^26^. At low resolution, LPB_All_ and PSP_All_ had h^2^_SNP_ estimates of 3.55% (SE=0.47%, Z-score=7.55) and 3.08% (SE=0.56%, Z-score =5.50). The range of h^2^_SNP_ estimates increased to 2.13% (SE=2.36%; LPB_Adult_, Z-score=0.93) and 4.06% (SE=0.62%; LPB_Adol_, Z-score=6.55) when allowing for differences in social domain and broad developmental stages (medium-resolution GWAS), and even further to 2.25% (SE=0.65%; parent-reported PSP 6 years, Z-score=3.46) and 6.60% (SE=1.36%; teacher-reported LPB 13 years, Z-score=4.85) when accounting for variation in the social domain, reporter, and specific ages (high-resolution GWAS). Genetic correlations increased in range, too, when studied at higher resolution using LDSC^27^ (Supplementary Figure 16-17). Low-resolution GWAS (LPB_All_ and PSP_All_) were moderately interrelated (r_g_=0.58, SE=0.096, Supplementary Figure 16a). Allowing for differences in the social domain and broad developmental stages (medium-resolution GWAS), patterns of high genetic similarity emerged (Supplementary Figure 16b), and accounting for variation in social domain, reporter, and age (high-resolution GWAS, Supplementary Figure 17), revealed increasingly complex genetic interrelationships necessitating a multivariate model description.

**Figure 3.**
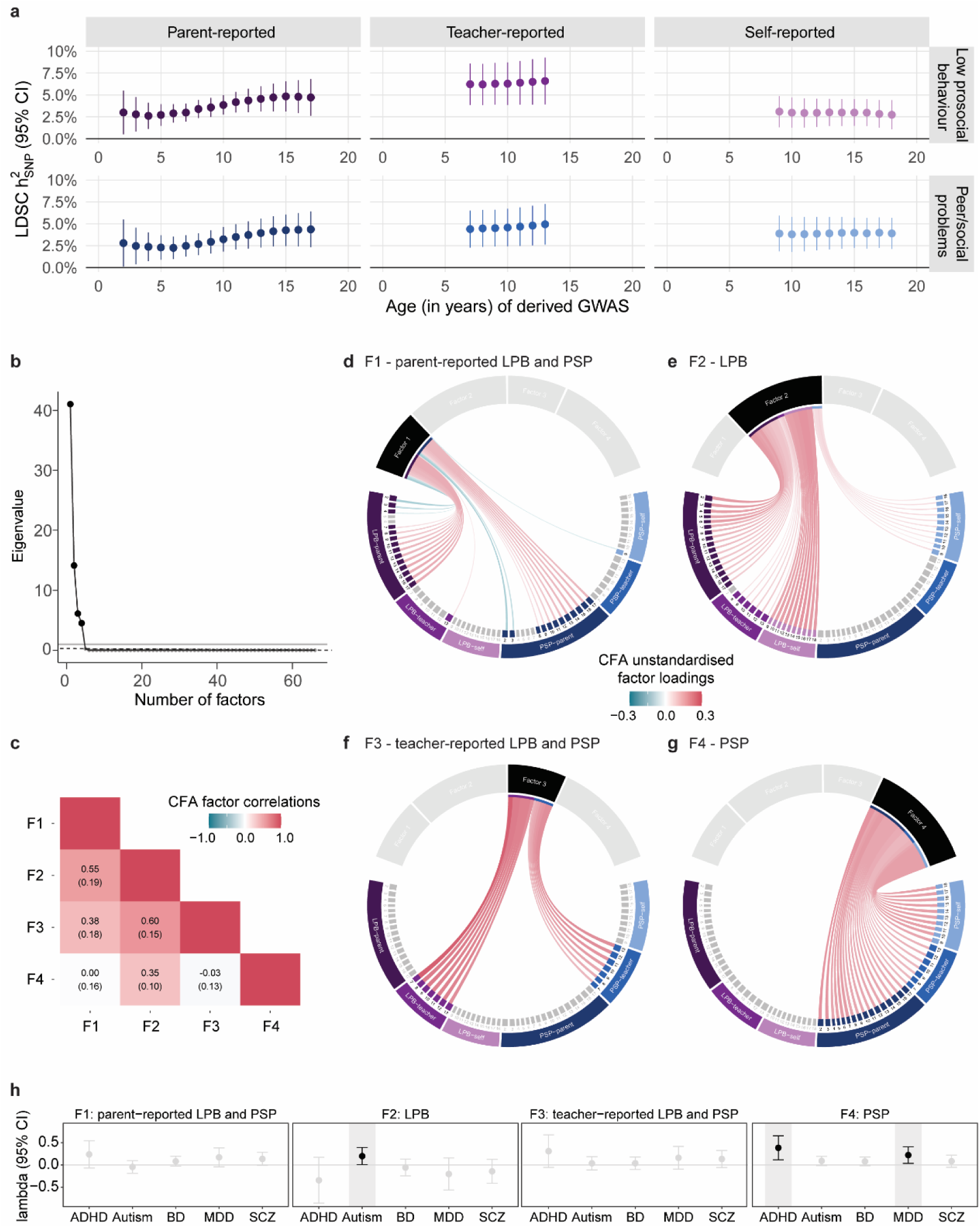
Genetic architecture of social behaviour. (**a**) LDSC h^2^_SNP_ across social domain, reporter, and age shown with corresponding 95% confidence bands across 66 high-resolution social behaviour GWAS. (**b**) Scree plot of genetic eigenvalues. (**c**) Factor correlations of a correlated 4-factor genetic CFA model of social behaviour and (**d-g**) corresponding unstandardised factor loadings, where the colour represents the magnitude of each factor loading. Detailed model information, including all factor loadings, is available in Supplementary Table 15. (**h**) Mapping NNCs onto the 4-factor social behaviour model, shown with their unstandardised factor loadings (p≤0.05, black). (**d-h**) Note that unstandardised factor loadings reflect relationships at the level of the captured h^2^_SNP._ Abbreviations: ADHD (Attention-Deficit/Hyperactivity Disorder), ASD (Autism Spectrum Disorder), BD (bipolar disorder), CFA (Confirmatory factor analysis), LPB (Low prosocial behaviour), MDD (major depressive disorder), NNCs (Neurodevelopmental and neuropsychiatric conditions), PSP (Peer and social problems), SCZ (schizophrenia)

We dissected the genetic covariance across high-resolution GWAS (N_GWAS_=66, Supplementary Note 5) using exploratory factor analysis (EFA) and confirmatory factor analysis (CFA) techniques, adopting a data-driven approach (Methods, Figure 3b-g)^28^. A 4-factor CFA model with correlated factors (Figure 3c-g, Supplementary Tables 13-15, Supplementary Figure 18, Supplementary Note 5) fitted the data best (CFI=0.98, SRMR=0.05, Supplementary Table 14), as identified with genomicSEM^29^ software. The four factors explained genetic variation across parent-reported LPB and PSP (F1, largest factor loading parent-reported LPB at 14 years: λ_F1_=0.14(SE=0.033), Figure 3d), parent-/teacher-/self-reported LPB (F2, largest factor loading self-reported LPB at 15 years: λ_F2_=0.18(SE=0.017), Figure 3e), teacher-reported LPB and PSP (F3, largest factor loading teacher-reported LPB at 7 years: λ_F3_=0.27(SE=0.034), Figure 3f), and parent-/teacher-/self-reported PSP (F4, largest factor loading parent-reported PSP at 7 years: λ_F4_=0.17(SE=0.019), Figure 3g). Thus, the identified factor structure aligned either with differences in the social domain (LPB: F2; PSP: F4) or informants (F1: parent reports, F3: teacher reports). Factor correlations (Figure 3c) ranged from - 0.03(SE=0.13) to 0.60(SE=0.15)), strongest between F2 and F3 (r_g_=0.60(SE=0.15)) and F1 and F2 (r_g_=0.55(SE=0.19)). Notably, F1 captured also an inverse association between parent-reported PSP in early childhood and LPB in middle/late childhood, also found with bivariate correlations (e.g. PSP at 2 years and LPB at 13 years, r_g_=-0.71(SE=0.22); Supplementary Figure 17), consistent with qualitative developmental changes in social behaviour.

### Polygenic scores predict contextual differences in social behaviour

We validated PGS for social behaviour in four European- and African-ancestry cohorts (N_ind_=16,305, N_obs_=115,254, Supplementary Notes 6-7).

Studying children of European ancestry from the Millennium Cohort Study (MCS)^30^, a cohort with extensive LPB and PSP phenotype information repeatedly assessed during childhood and adolescence by different informants using the SDQ^4^ (N_ind_=6,487, N_obs_=96,259, Supplementary Table 16), we evaluated the predictive ability and predictive accuracy of social behaviour PGS.

First, we confirmed the predictive ability of PGS, matching the contextual information of each discovery social behaviour GWAS to the corresponding social behaviour target in the MCS (18 LPB and PSP measures, ages 3 to 17 years, parent-/teacher-/self-reported, Supplementary Table 16). Matching discovery-target pairs yielded consistently positive effects (incremental-R^2^ = 0.18-1.13%, Supplementary Table 17, Supplementary Figure 19), with all associations, except for parent-reported PSP at 3 years, passing the multiple testing threshold (p<0.0037; Methods).

Second, we evaluated the predictive accuracy of PGS as the percentage of associations with larger effects (in incremental-R^2^) for phenotypically matching discovery-target pairs than non-matching pairs (Supplementary Tables 18-19, Supplementary Note 7), for all associations passing the multiple testing threshold. For domain-matched PGS signals, the predictive accuracy was high, especially for the PSP social domain (PSP=81.6%, LPB=45.6%, Supplementary Table 18). For reporter-matched PGS, the predictive accuracy was highest for parent-report (65.53%) and teacher-report (66.07%) outcomes, followed by self-reports (38.8%). For both domain- and reporter-matched PGS, the predictive accuracy increased even further (parent-report outcomes: 88.6%; teacher-report outcomes: 71.4%; self-report outcomes: 50%), although the predictive accuracy for self-reported LPB using self-report PGS was poor, and the outcome was better captured by parent-report PGS (Supplementary Table 19, Supplementary Figure 22). Self-reported pro-social behaviour measures are prone to socially desirable responding^31^, and this bias may affect genomic analyses, especially when using single-cohort self-report measures.

Third, comparing the largest PGS signal (in incremental-R^2^) with the fully matching PGS, we observed, for associations passing the multiple testing threshold, 89% domain-matched, 78% reporter-matched and 78% age-matched (within a 5-year age window) signals capturing contextual information of the target phenotype, with similar magnitude (Figure 4, Supplementary Table 18, Supplementary Note 7). Thus, while matching discovery and target characteristics increased predictability (incremental-R^2^≤1.32%, Figure 4, Supplementary Table 17, Supplementary Figures 20-21), meta-regression-derived PGS also predicted contextual target information, if unmatched, except for self-report LPB outcomes.

**Figure 4.**
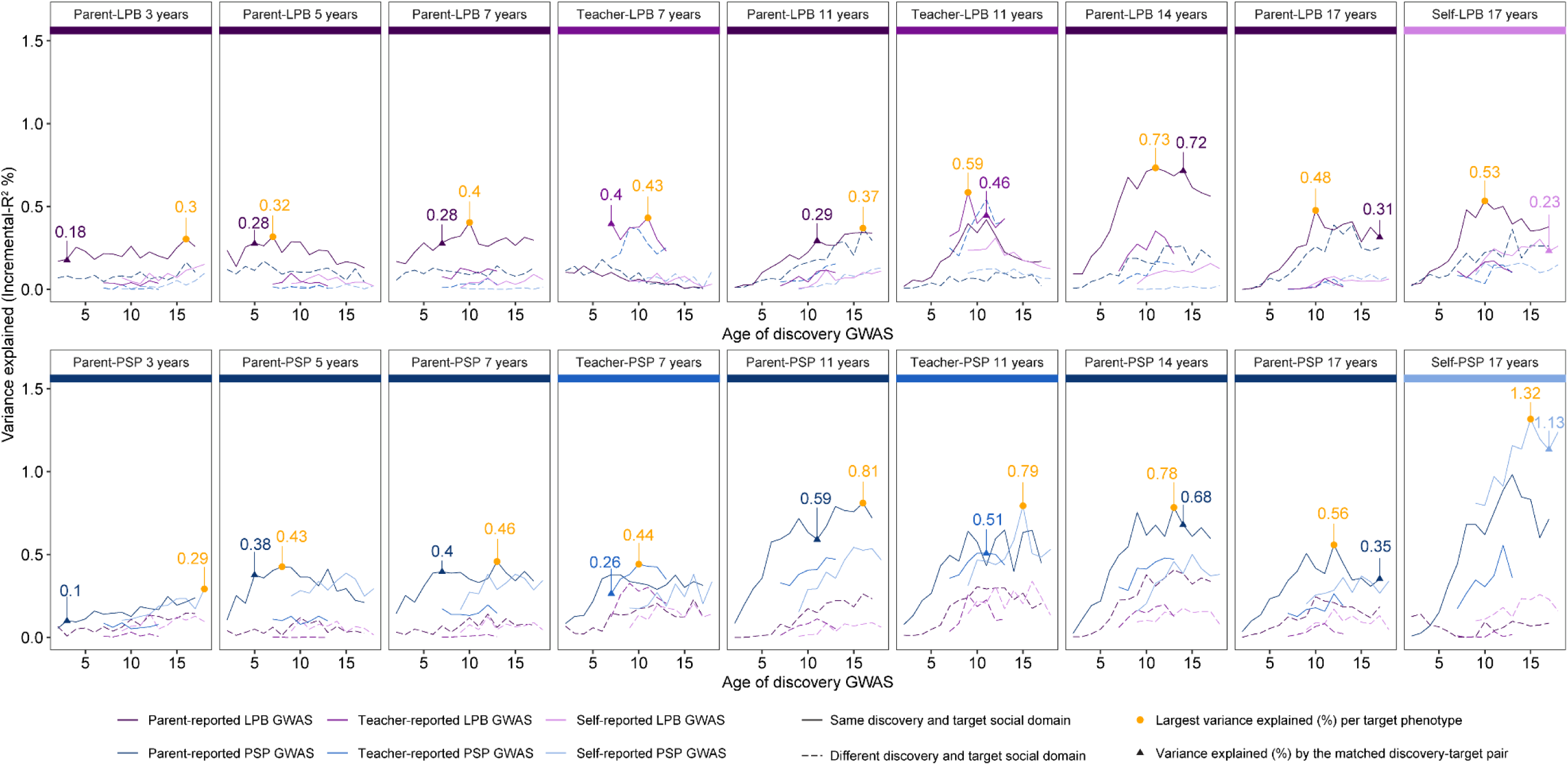
Polygenic predictability of social behaviour. Social measures in the Millennium Cohort Study (MCS) (target) were predicted with PGS from 66 high-resolution social behaviour GWAS, allowing for changes in social domain, reporter and age (discovery). For each target phenotype (panel), the explained variance (incremental-R^2^ percentage, y-axis) is shown against the predicted age (in years) of the discovery GWAS. Blue lines indicate PSP discovery GWAS based on parent (darker), teacher (medium), and self (lightest) reports. Purple lines indicate LPB discovery GWAS based on parent (darker), teacher (medium), and self (lightest) reports. Lines are solid if the target and discovery social domains match (e.g. parent-reported LPB GWAS predicting parent-LPB at 3 years in the MCS) and dashed otherwise. For each target phenotype, the variance explained by a matching PGS, aligned by social domain, reporter and age, is highlighted with a triangle. For each target phenotype of the MCS, the largest variance explained by any discovery PGS is shown in orange. Abbreviations: GWAS (Genome-wide association study), LPB (Low prosocial behaviour), PGS (Polygenic score), PSP (Peer and social problems)

Subsequently, we corroborated the phenotypic transferability of social behaviour PGS to broadly defined social measures assessed with non-standardised instruments during early and late adulthood, studying two further European-ancestry samples (incremental R^2^≤0.22%, total N_ind_=8,983, Supplementary Tables 20-21, Supplementary Note 7).

Lastly, we predicted SDQ and CBCL measures in an African-ancestry cohort (N_ind_=835, Supplementary Table 16 and 22, Supplementary Note 7). Matching social domains and development stages, social behaviour GWAS PGS was associated with parent-reported LPB at ages 7 and 8 (incremental-R^2^≤0.95%), highlighting the cross-ancestry transferability of LPB PGS.

### Genetic correlates of social behaviour reveal contextual association patterns

With PGS systematically capturing trait heterogeneity, we explored genetic correlation patterns with social behaviour in depth, capitalising on contextual information, including genetically-inferred ages that allow predicting the onset of associations with social behaviour.

First, we examined genetic correlations across a spectrum of population-based traits (N_traits_=51), including externalising, internalising, social, wellbeing, personality, cognition and educational attainment (EA), language, in addition to anthropometric and birth measures, using LDSC software^26^ (Figures 5-6, Supplementary Table 23, Supplementary Figures 23-24, Methods, multiple testing threshold of p<8.89x10^-4^).

**Figure 5.**
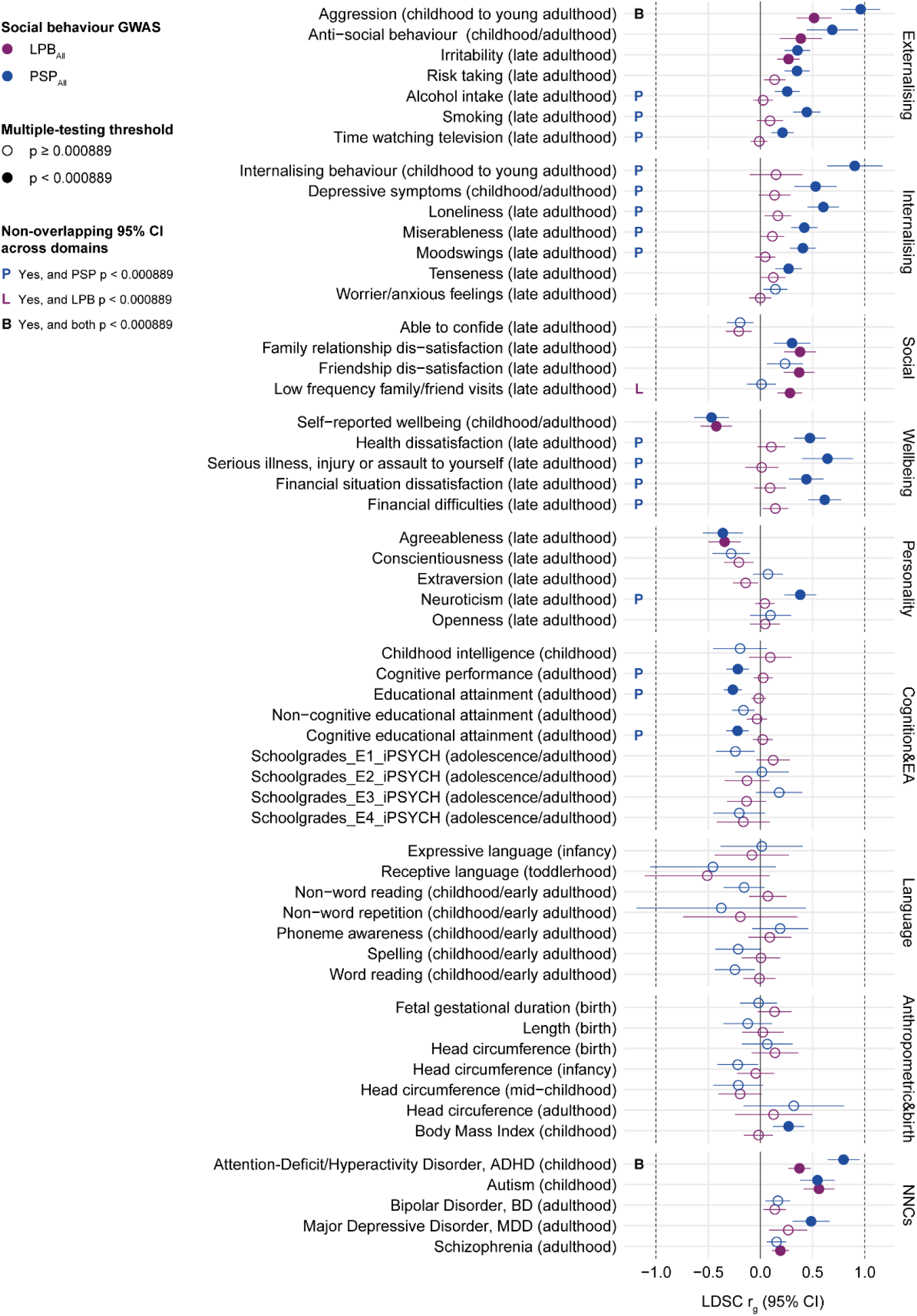
Genetic correlations of low-resolution social behaviour with population-based traits and NNCs. Linkage disequilibrium score (LDSC) genetic correlations with LPB_All_ and PSP_All_ GWAS are shown. Filled circles indicate correlations passing the multiple testing threshold (p<8.89x10^-4^). Differences in PSP and LPB genetic correlations, indicating specific effects based on non-overlapping 95% confidence bands, are shown for PSP correlations (blue P), LPB correlations (purple L) or both (black B), once they passed the multiple testing threshold. Abbreviations: LPB (Low prosocial behaviour), NNCs (Neurodevelopmental and neuropsychiatric conditions), PSP (Peer and social problems)

**Figure 6.**
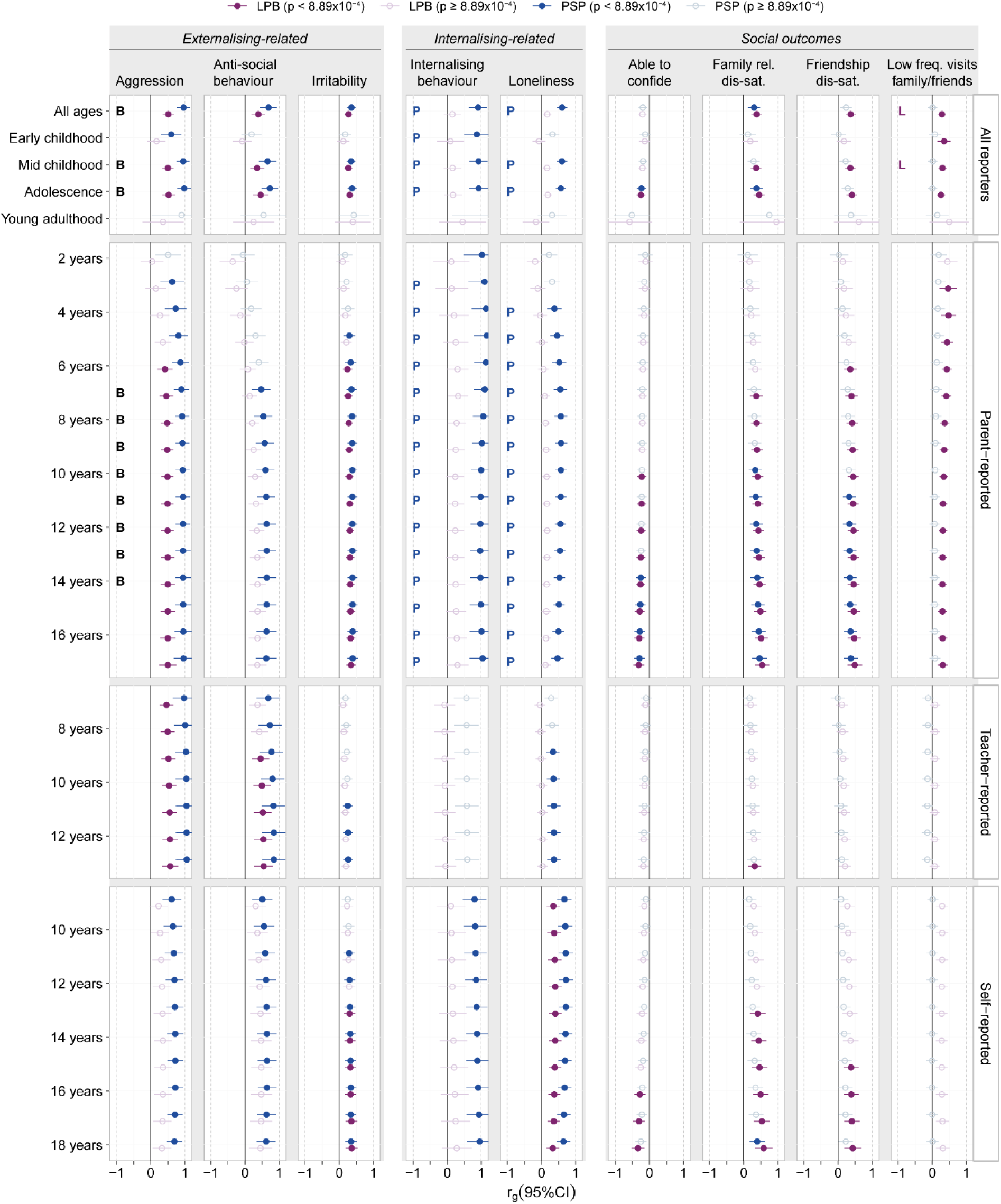
Contextual genetic correlations of social behaviour with (near) adult externalising, internalising and social outcomes. Linkage disequilibrium score (LDSC) genetic correlations with low-resolution, medium-resolution and high-resolution LPB and PSP GWAS are shown. Filled circles indicate correlations passing the multiple testing threshold (p<8.89x10^-4^). Differences in PSP and LPB genetic correlations at different moderator levels, indicating specific effects based on non-overlapping 95% confidence bands, are shown for PSP correlations (blue P), LPB correlations (purple L) or both (black B), once they passed the multiple testing threshold. Abbreviations: LPB (Low prosocial behaviour), PSP (Peer and social problems)

The largest and strongest correlates of social behaviour were behavioural outcomes (Figure 5). Genetic correlations with (near) adult externalising traits were positive and distinctly larger for PSP compared to LPB, with non-overlapping 95% confidence bands (e.g. aggression: PSP_All_-r_g_=0.96(SE=0.10); LPB_All_-r_g_=0.51(SE=0.09)), indicating domain-specific effects. For non-substance-use-related externalising behaviour, the predicted onset of PSP associations preceded LPB associations, especially for parent-report PSP (Figure 6, Supplementary Figure 24). For (near) adult internalising traits (e.g. internalising behaviour: PSP_All_-r_g_=0.9(SE=0.14); loneliness: PSP_All_-r_g_=0.6(SE=0.08)), many lack-of-wellbeing traits, substance-use-related traits and neuroticism, we observed moderate-to-strong PSP-specific correlations, predicted to emerge in early childhood, e.g. based on parent-report GWAS (Figures 5-6, Supplementary Table 25, Supplementary Figures 23-24). Genetic correlations with EA and cognition were modest, inverse and PSP-specific, too (e.g. EA: PSP_All_-r_g_=-0.26). In contrast, for social outcomes as well as agreeableness, the predicted onset of LPB associations preceded PSP associations, especially for parent-report GWAS (Figure 6, Supplementary Table 25, Supplementary Figures 23-24). Genetic correlations consistently involved LPB, with weak-to-moderate effects that were either similar for both LPB and PSP (e.g. family relationship dissatisfaction: LPB_All_-r_g_=0.38(SE=0.08); PSP_All_-r_g_=0.30(SE=0.09)), or LPB-specific (e.g. low-frequency-of-family/friends visits: LPB_All_-r_g_=0.28(SE=0.06)).

We next investigated the genetic correlation profile for five clinically defined NNCs (Figure 3h, Figure 5, Figure 7, Supplementary Table 25): ADHD, autism, bipolar disorder (BD), major depressive disorder (MDD) and schizophrenia. For ADHD, both PSP- and LPB-specific associations were observed with larger correlations for PSP (PSP_All_-r_g_=0.80(SE=0.078); LPB_All_-r_g_=0.37(SE=0.055)), particularly with parent- and teacher-report GWAS (Figure 5, Figure 7a). MDD was moderately and positively associated with PSP (PSP_All_-r_g_=0.49(SE=0.091)) throughout, and only related to LPB for parent-report GWAS at 13 to 17 years (Figure 7a). For both ADHD and MDD, the predicted onset of PSP correlations in early and mid-childhood was earlier than for LPB correlations, especially for parent-report GWAS (Figure 7a), and both were linked to the social behaviour PSP factor (F4, Figure 3h). Autism was moderately correlated with both social domains (Figure 5; LPB_All_-r_g_=0.56(SE=0.08); PSP_All_-r_g_=0.55(SE=0.08)) across all reporters, with early-onset associations, especially as predicted by parent-report GWAS (Figure 7a). For self-reported LPB, genetic correlations with autism were consistent with one (e.g. at 15 years; LPB-r_g_=0.80(SE=0.15); Figure 7a), indicating a strong genetic overlap between population-based and clinical social dimensions^32^, consistent with genetic links between autism and the social behaviour LPB factor (F2, Figure 3h). Schizophrenia showed a modest genetic overlap with LPB (LPB_All_-r_g_=0.19(SE=0.042)), most apparent for parent- and teacher-report GWAS (Figure 7a), in line with our previous work^33^. Several parent-reported PSP GWAS (ages 7-17 years) shared genetic links with schizophrenia, too, but the predicted onset of LPB correlations preceded PSP correlations (if present) in mid-childhood (Figure 7a). Schizophrenia was not related to a social behaviour factor (Figure 3h). In line with previous studies^33^, there was little evidence for an association between social behaviour and BD.

**Figure 7.**
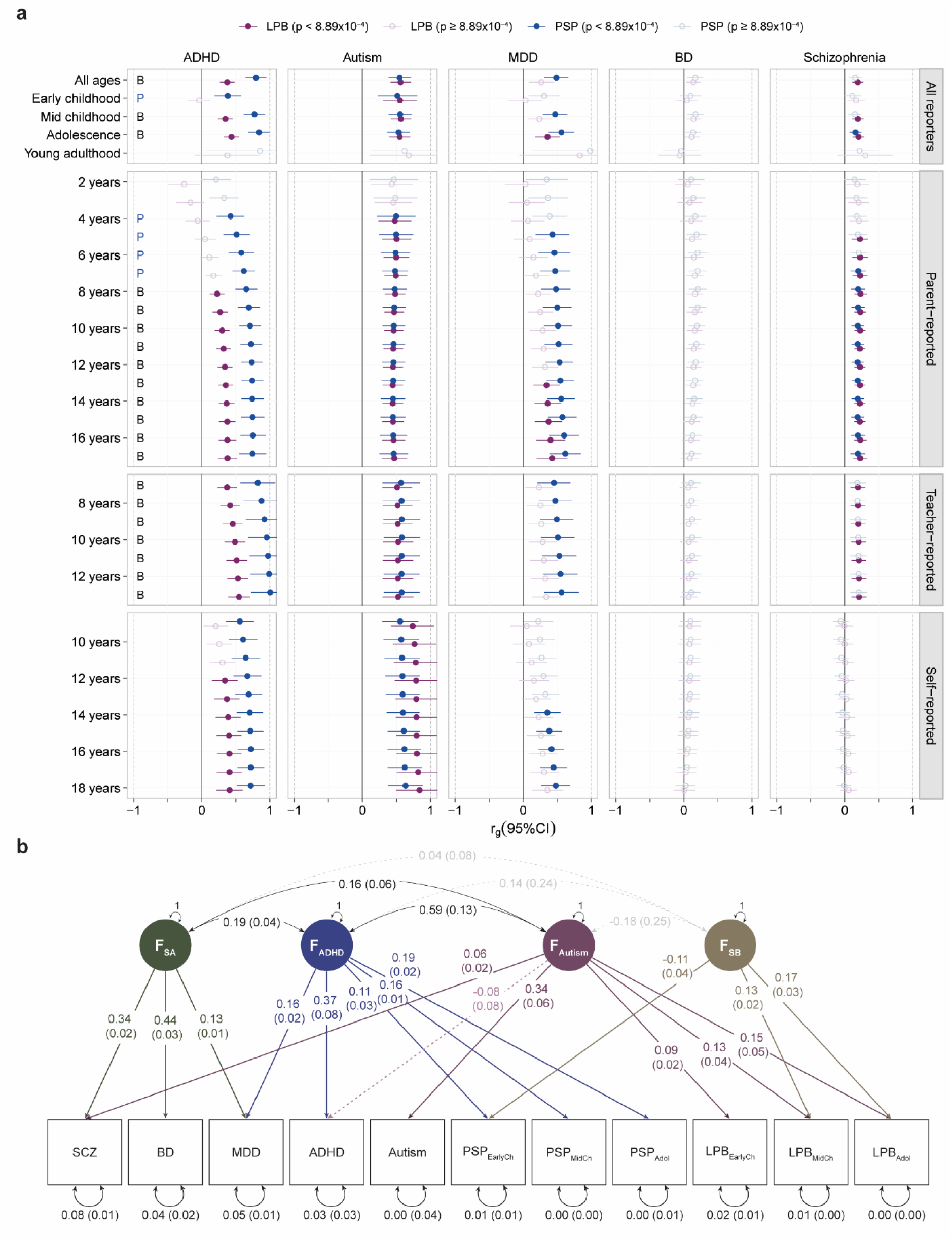
Genetic architecture across social behaviour and NNCs. (**a**) Contextual genetic correlations of social behaviour with NNCs. Linkage disequilibrium score (LDSC) genetic correlations with low-resolution, medium-resolution and high-resolution LPB and PSP GWAS are shown. Filled circles indicate correlations passing the multiple testing threshold (p<8.89x10^-4^). Differences in PSP and LPB genetic correlations at different moderator levels, indicating specific effects based on non-overlapping 95% confidence bands, are shown for PSP correlations (blue P), LPB correlations (purple L), or both (black B), once they passed the multiple testing threshold. (**b**) Genetic CFA across medium-resolution social behaviour and NNCs. The CFA model provided an optimal model fit (CFI=0.99, SRMR=0.069). Observed measures are represented by squares and latent variables by circles. Coloured, single-headed arrows define unstandardised factor loadings (shown with their corresponding SEs). Factor loadings with p ≤ 0.05 are solid, and dashed otherwise. Double-headed arrows define factor correlations (shown with their corresponding SEs). Factor correlations are represented with a black solid line if p ≤ 0.05 and with a dashed grey line otherwise. Note that unstandardised factor loadings reflect relationships at the level of the captured h^2^_SNP._ Analyses were restricted to social behaviour GWAS statistics with sufficient power (h^2^_SNP_ Z-score >3). Abbreviations: ADHD (Attention-Deficit/Hyperactivity Disorder), BD (bipolar disorder), F_ADHD_ (ADHD genetic factor), F_Autism_ (autism genetic factor), F_SA_ (schizoaffective genetic factor), F_SB_ (social behaviour genetic factor), LPB (Low prosocial behaviour), LPB_EarlyCh_ (Early childhood LPB), LPB_MidCh_ (Mid childhood LPB), LPB_Adol_ (Adolescence LPB), MDD (major depressive disorder), NNCs (Neurodevelopmental and neuropsychiatric conditions), PSP (Peer and social problems), PSP_EarlyCh_ (Early childhood PSP), PSP_MidCh_ (Mid childhood PSP), PSP_Adol_ (Adolescence PSP), SCZ (schizophrenia)

To triangulate findings, we also fitted a structural model across NNCs and medium-resolution social behaviour GWAS (Supplementary Tables 31-33), leveraging the larger h^2^_SNP_ of NNCs for factor identification. Using genomic SEM, we identified a correlated 4-factor CFA model with a good model fit (CFI=0.99, SRMR=0.069; Figure 7b, Supplementary Figure 27, Supplementary Note 8, Methods). Two intercorrelated factors (r=0.59(SE=0.13)) linked NNCs and social behaviour: F_ADHD_ (most strongly related to ADHD, λ=0.37(SE=0.08)) explaining shared genetic variation underlying ADHD, MDD and PSP across development (Figure 7b), and F_Autism_ (most strongly related to autism, λ=0.34(SE=0.06)), capturing genetic variation shared between autism, LPB across development and, weakly, schizophrenia. Both F_ADHD_ and F_Autism_, were related to EA, once mapped onto the factor structure (Supplementary Figure 28, Supplementary Table 34), consistent with socio-economic contributions.

### Comparison of meta-regression with conventional meta-analytic approaches

We compared our meta-regression-based findings with those from a conventional meta-analysis approach, such as multivariate N-GWAMA^34^ that assumes no contextual heterogeneity (Supplementary Note 9). Notably, the magnitude and strength of h^2^_SNP_ estimates decreased when adopting a conventional approach (Supplementary Table 35). Still, LDSC intercepts, which capture the mean inflation due to confounding bias^26^, were below one for all meta-regression GWAS, suggesting an (overly) conservative approach when accounting for sample overlap (Supplementary Table 12). Genetic correlations across low-resolution social domains derived with either method were strong (Supplementary Figure 29, Supplementary Table 35), and genetic correlation patterns with population-based traits and NNCs were highly consistent (Supplementary Figure 30, Supplementary Table 36), indicating that the presented results are robust to methodological choices.

## DISCUSSION

Adopting a genome meta-regression approach (N_eff_=121,777, N_ind_=73,321, 491,246 repeat observations) and following up findings in European- and African-ancestry cohorts (N_ind_=16,305), this study demonstrates that social behaviour is heterogeneous with systematic changes in genetic effects across social domains, informants, and age. Here, we (i) identified GWA signals with potential biological impact, (ii) characterised the multidimensional genetic architecture of social behaviour, (iii) generated and validated PGS predicting the context of social behavioural difficulties, (iv) uncovered genetic correlates of social behaviour with contextual association patterns and (v) enhanced the insight into the genetic architecture of population-based traits NNCs by predicting the age of association onset.

Our study confirms that social behaviour is a highly context-dependent trait, with social context being captured by both differences in single-variant genetic effects and genomic influences. We report six GWA loci associated with social behaviour, all of which had either domain-varying, age-varying and/or reporter-varying genetic effects. Three loci (rs148774891, rs150685860, and rs538645) were mapped to genes with a high mutation intolerance (pLI>0.9), including *CADM2* (rs148774891), *XKR7* (rs150685860) and *ARNC1* (rs538645), or high load of deleterious variation (CADD>12.37) such as *SETP16* (rs538645). rs148774891 captured a large increase in self-reported childhood LPB (0.2 SD) and was mapped to *CADM2,* a gene that has been previously linked to autism^35,36^, but also risk-taking behaviour^37^, smoking and alcohol consumption^38^, externalising behaviour^39^, EA^40^, and general cognitive ability^41^, with development-specific expression patterns in the orbitofrontal cortex. *ARNC1,* mapped to rs538645, although not yet associated with cognition and behaviour, is over-expressed in a development-dependent manner in subcortical structures that play a role in regulating social behaviour^25^, and the locus is in weak LD with known GWAS signals for depression within *CXCR5* (r^2^≤0.35)^42^. *XKR7* has not been implicated in a known GWAS as yet, but the rare protective effect at rs150685860, associated with a 0.16 SD decrease in early childhood parent-reported PSP, suggests possible resistance mechanisms with translational potential, analogous to effects reported in Alzheimer’s disease^43^.

The heritability of social behaviour was modest (h^2^_SNP_ =2-7%) but comparable to adult social traits^44–46^, and most social behaviour GWAS were sufficiently powered (mean h^2^_SNP_ Z-score=4.41, range: 0.90-7.55), except for medium-resolution adult GWAS and very early childhood GWAS (h^2^_SNP_ Z-score≤3). The identification of four correlated genetic factors using a genetic CFA model underlines the heterogeneous nature of social behaviour^1^, disentangling here differences in informant characteristics from differences in social domains, of which only the latter were related to NNCs. The lack of genetic links between both the parent-report (F1) and teacher-report (F3) factors with the PSP factor (F4) is consistent with informant-specific characteristics, widely known in twin research^15^, including differences in knowledge about the young person, but also in social propensities elicited by different situations.

The novelty of this study lies not only in the systematic modelling of heterogeneity in genetic effects underlying social behaviour, but also, after extensive validation, in the systematic prediction of contextual information when studying genetic correlates. Correlational analyses can provide insight into the studied trait through associated phenotypes. We outline here novel forms of contextual correlations that support the search for underlying developmental processes, including the predicted onset of associations.

Genetic correlation patterns showed strong PSP-specific links to a wide range of (near) adult externalising, internalising and lack-of-wellbeing measures, but also ADHD and MDD. Correlational patterns predicted the earlier emergence of links with PSP rather than LPB (if present), particularly for parent-report GWAS. Consistently, genetic similarity across ADHD, MDD and PSP was identified with complementary genetic CFA models.

LPB was genetically related to a narrower range of adult population-based traits and primarily associated with challenges in forming and maintaining quality relationships, likely due to a reduced tendency to engage in helpful and sharing behaviours^47^, but also autism and schizophrenia. Genetic overlap across LPB and autism was detected using complementary genetic CFA models, and links were weakly shared with schizophrenia. In particular, an earlier predicted onset of parent-reported LPB rather than PSP associations was observed for both later-life social outcomes and schizophrenia. In comparison, for autism, moderate-to-strong genetic correlations with both PSP and LPB, predicted to emerge from 4 years onwards across all reporters, matched the typical age of diagnosis^48^ with social difficulties involving both a lack of sharing with others and problems in interacting with others^49^. Thus, supported by similarities across multiple social correlates and the largely comparable power in social behaviour GWAS, the identified correlational patterns and structures are consistent with differences in developmental processes, where either PSP, LPB or both mark early stages in pathways leading to different later-life and mental health outcomes, without precluding bi-directional effects between PSP and LPB during development.

This study demonstrates that meta-regression approaches are capable of modelling heterogeneity in genetic effects genome-wide, harmonising cohort and phenotype information, especially through phenotypic mappings to the ICF^19^, thus moving the field closer towards dynamic GWAS analyses. We provide a scalable analytical framework for capturing the genomic basis of context-dependent traits, including developmental changes, that allowed us to gain a deeper insight into the individual differences underlying social behaviour. Meta-regression-derived PGS demonstrated predictive ability and context-specific accuracy, especially for European-ancestry cohorts. When discovery and target characteristics were matched, PGS captured up to half of the h^2^_SNP_ for social behaviour GWAS (incremental-R^2^≤1.32%), consistent with research showing that calibrating PGS analyses to different contexts enhances PGS accuracy^50^. Capitalising on the context-specific accuracy of meta-regression-derived PGS, meta-information could be inferred, even with little information about target characteristics. However, the prediction accuracy will depend on factors such as the studied population^50^, ancestry^50^ and also bias (e.g. socially desirable responding^31^) and was notably poor for self-reported LPB outcomes that were better predicted using parent-report LPB PGS.

Despite a considerable sample size capitalising on repeat assessments, we detected only a few GWA loci. The adjustment for repeated measures in this study might have been overly conservative, consistent with the lower power of meta-regression approaches to detect main effects in the absence of moderators^16^. Nonetheless, we demonstrated that meta-regression techniques are a powerful meta-analytic tool, enhancing both granularity and power of meta-GWAS beyond conventional meta-GWAS approaches. Expectedly, the predicted social behaviour GWAS will only partially proxy social-behavioural heterogeneity and additional moderators are likely to explain further variance, although an unspecific increase in the number of meta-regression moderators will lower statistical power. While we find LPB cross-trait transferability to children of African ancestry, future studies will benefit from non-European-ancestry discovery populations.

In conclusion, we have shown that the genetic architecture of social behaviour is multidimensional and context-dependent. Adopting a meta-regression approach, we demonstrated that heterogeneity in genetic effects can be modelled and is genetically predictable. Leveraging contextual information in GWAS and PGS offers new insight into the developmental processes linking social behaviour to later-life outcomes and mental health.

## METHODS

### Study cohorts

Within this study, we included data from 25 independent cohorts (Supplementary Table 1) across 11 different countries (73,321 individuals of European ancestry), embedded within the behaviour and cognition working group of the EAGLE consortium^17^ (https://www.eagle-consortium.org/working-groups/behaviour-and-cognition/). For each participating cohort, ethical approval was granted by their local research ethics committee, and written informed consent was given by the participants and/or their parents/legal guardians (Supplementary Note 1). Each cohort provided GWAS summary statistics and phenotype descriptive data on PSP and/or on prosocial behaviour (which was reverse-coded into LPB) assessed with parent-, teacher-, and/or self-reports across ages 2 to 29 years. Detailed information on the cohorts and phenotypes is available in Supplementary Tables 2-3 and Supplementary Note 1.

### Phenotype definition

Prosocial behaviour measures were reverse-coded (LPB) to align with the PSP measures, where higher scores reflect more social behaviour difficulties. Where available, we studied repeatedly assessed (parent-, teacher-, self-reported) measures of PSP and LPB up to the age of 29 years (N_observations_=491,246, Supplementary Table 2).

Across the cohorts, dimensions of social behaviour (LPB and PSP) were predominantly assessed with the SDQ^4^ and the CBCL^18^ (Supplementary Table 3). Both questionnaires are widely used, validated, and standardised instruments assessing behavioural problems for research and clinical practice in both clinical and general population-based settings^51^. All other instruments were mapped to either the LPB or PSP domain using the ICF^19^ (Supplementary Table 3) by aligning all questionnaire items to the ICF^19^ (Supplementary Note 2, Supplementary Table 4).

### Cohort-specific association analyses

Within each cohort, participants were genotyped using high-density SNP arrays (Supplementary Table 37). Genotype data were subjected to stringent quality control and subsequently imputed against the Haplotype Reference Consortium version 1.1 (HRCr1.1) panel^52^ or the 1000 Genomes Project Phase 3^53^ as reference panel, following standard procedures for GWAS (Supplementary Note 10, Supplementary Table 37).

Per cohort, social behaviour scores for each questionnaire were jointly Z-standardised across all available reporters and ages such that, within each cohort, absolute differences across sex, reporters and ages could still be captured while allowing for comparable effect sizes across cohorts. Subsequently, univariate GWAS analyses were performed using linear regression methods, assuming an additive linear model by regressing Z-standardised scores on posterior genotype probability while adjusting for covariate effects (sex, age, age^2^, and ancestry-informative principal components). Note that the number of principal components differed across cohorts. In cases where adjusting for age^2^ led to collinearity-related problems, this covariate was dropped from the linear model. Association analyses were carried out with different software as described in the Supplementary Notes (Supplementary Note 10, Supplementary Table 37).

### Quality control of GWAS summary statistics

Quality control of the GWAS summary statistics from all participating cohorts was carried out using the EasyQC^54^ R package (*R::EasyQC*, v9.2). SNPs were excluded if imputation quality scores were lower than 0.6, minor allele frequency was lower than 0.5%, and/or minor allele count was ≤10. Furthermore, alleles were aligned against the HRCr1.1 reference panel^52^, and variants were excluded if not contained in the reference panel, alleles were missing or mismatched, or the reported allele frequency deviated more than 0.2 from the reported HRCr1.1^52^ frequency. Additionally, insertions/deletions, duplicate SNPs, and multi-allelic SNPs were excluded. The final number of SNPs retained for each cohort is available in Supplementary Table 37.

GWAS summary statistics for which h^2^_SNP_ values (as computed with LDSC^26^) had either an upper 95% confidence interval below zero or a lower 95% confidence interval above one were removed from the analysis to exclude unreliable measures. GWAS summary statistics were also removed if GWAS summary statistics for either male or female subsamples (created for quality control purposes) had either an upper 95% confidence interval below zero or a lower 95% confidence interval above one.

### Genome-wide meta-regression analysis

To meta-analyse the 195 univariate GWAS summary statistics, we carried out a random-effects meta-regression with the metafor^20^ R package (*R::metafor*, v3.0-2). Compared to fixed-effect meta-analysis, random-effect meta-analysis allows for heterogeneity of the true genetic effect sizes across different cohorts that may arise due to, for instance, differences in sample characteristics^16^.

For each SNP, the variation (i.e. heterogeneity) in SNP genetic effects across cohorts was modelled using social domain, reporter, age and age^2^ as moderators (Eq 1):

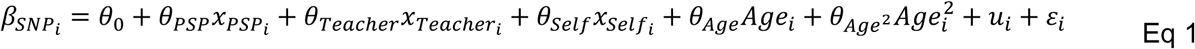

where 𝛽_𝑆𝑁𝑃𝑖_ is the genetic association effect for the SNP for cohort 𝑖; 𝜃_0_ is the mean genetic SNP effect of parent-reported LPB at the grand N-weighted mean age (age=10.14 years, i.e. for each SNP the grand mean age was centred at 0); 𝜃_𝑃𝑆𝑃_ is the difference in SNP effect when measuring PSP; 𝜃_𝑇𝑒𝑎𝑐ℎ𝑒𝑟_ is the difference in the SNP effect when assessed with teacher reports; 𝜃_𝑆𝑒𝑙𝑓_ is the difference in the SNP effect when using self-reports; 𝜃_𝐴𝑔𝑒_ is the difference in the mean SNP effect for an increase in one year of age; 𝜃_𝐴𝑔𝑒2_ is the difference in the mean SNP effect for an increase in one year in the square of age; 𝑢_𝑖_, distributed as 𝑢_𝑖_∼𝑁(0, 𝜏^2^), represents the random intercept for the 𝑖^𝑖𝑡ℎ^ cohort, thus, describing study-specific variation from the mean SNP effect or between-cohort sampling error (where 𝜏^2^is the between-cohort heterogeneity); 𝜀_𝑖_, distributed as 𝜀_𝑖_∼𝑁(0, 𝜎_i_^2^), represents the within study sampling error with sampling variance 𝜎_i_^2^ in the 𝑖^𝑖𝑡ℎ^cohort (i.e. the within-cohort heterogeneity). To estimate the between-cohort variance (𝜏_i_^2^), we used a restricted maximum likelihood (REML) estimator^55^ and a nested random-effects structure where each variable is nested within its corresponding cohort.

The sampling variance (𝜎_i_^2^) within each cohort i was approximated by accounting for repeated observations for the same individuals across different social domains, reporters and ages (Eq 2). This model is analogous to models accounting for correlated phylogenetic histories^56^, where, within each cohort i, S is a diagonal matrix with the standard errors (SE) of the 𝛽_𝑆𝑁𝑃_ effect sizes across 𝑘 GWAS summary statistics, and Σ_pheno_ is the Person’s phenotypic correlation matrix of social scores with 𝑘 𝑥 𝑘 dimensions.

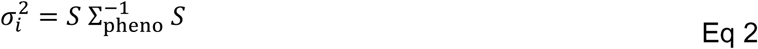

Note that the phenotypic correlation matrix (Σ_pheno_) was adjusted for sample overlap following the formula by Bulik-Sullivan-Finucane and colleagues^27^ (Eq 3), where ϱ is the phenotypic correlation between two measures, N_s_ is the number of overlapping samples, 𝑁_1_ is the sample size of the first measure, and 𝑁_2_ is the sample size of the second measure. Assuming that sampling errors are independent between cohorts, phenotypic correlations of measures from different cohorts were set to zero in the phenotypic correlation matrix (Σ_pheno_). Therefore, the full V matrix per SNP was constructed across all 𝜎_i_^2^, setting all covariance terms across cohorts to zero.

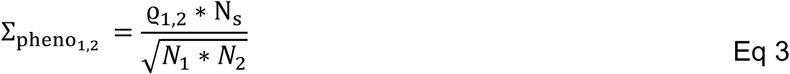

To allow for sufficient information to estimate the levels of the fixed effects, analyses were restricted to SNPs that were available for both domains (PSP and LPB), for the three reporters (parent, teacher and self), with at least three different age timepoints, and available in at least two cohorts, which resulted in 7,852,971 SNPs.

To quantify the amount of heterogeneity explained by the moderators and the model fit per SNP, in addition to the model in Eq 1, we fitted a (baseline) model without the fixed effects, i.e. only the intercept, the random effect term (𝑢_𝑖_) and the sampling error (𝜀_𝑖_). The model fit was assessed with a likelihood-ratio test across the baseline and the full covariate model.

### Deriving genome-wide genetic effects

For all SNPs for which the random-effects meta-regression model converged (N_SNPs_=7,830,986), predictions using the *predict.rma* function (*R::metafor,* v3.0-2)^20^ were conducted. We generated three groups of GWAS (Supplementary Table 5):

1. <u>Low-resolution GWAS: across social domain (2 GWAS)</u>. A grand estimate was derived for each social domain (LPB_All_, PSP_All_). To do so, we set the social domain moderator (θ_PSP_) to 0 for LPB and to 1 for PSP. Age (θ_Age_) and age^2^ (θ_Age_^2^) moderators were set to the N-weighted mean across all 195 summary statistics (10.14 years old). Reporter moderators (θ_Teacher_, θ_Self_) were set to the proportion of summary statistics (available for that SNP) assessed with teacher or self-reports. For example, if a SNP has 50 parent-reported, 20 teacher-reported and 10 self-reported measures, then the teacher-reported moderator (θ_Teacher_) will be set to 0.25 (20/80 total measures) and the self-reported moderator (θ_Self_) to 0.125 (10/80).
2. <u>Medium-resolution GWAS: across social domain and developmental stage (8 GWAS)</u>. Additionally, GWAS were derived per social domain (LPB and PSP) and developmental window: early childhood (ages 2-5; LPB_EarlyCh_, PSP_EarlyCh_), mid-childhood (ages 6-11; LPB_MidCh_, PSP_MidCh_), adolescence (ages 12-17; LPB_Adol_, PSP_Adol_), and young adulthood (ages 18-29; LPB_Adult_, PSP_Adult_). Age (θ_Age_) and age^2^ (θ_Age_^2^) moderators were set to the N-weighted mean age per developmental window across the 195 GWAS summary statistics. Specifically, this was 3.48 years for early childhood, 9.28 years for mid-childhood, 13.80 years for adolescence and 23.4 years for young adulthood. Reporter moderators were created as described in the “across social domain” predictions.
3. <u>High-resolution GWAS: across social domain, reporter, and age (66 GWAS)</u>. Derived GWAS were created based on a grid of moderator values that were contingent on the reported data across the 195 GWAS summary statistics (Figure 1, Supplementary Figure 31). Therefore, we used both social domains (LPB and PSP) and the three reporters (parent, teacher and self). To allow for a reliable estimation, for each combination of social domain and reporter, we restricted the age ranges that had data on at least two cohorts or that had data on one cohort but with a sample size above 2,000. Parent-reported LPB and PSP GWAS were derived from 2 to 17 years (16 ages x 2 domains = 32 GWAS). Teacher-reported LPB and PSP GWAS were derived from 7 to 13 years (7 ages x 2 domains = 14 GWAS). For self-reported LPB and PSP GWAS, the range was set from 9 to 18 years (10 ages x 2 domains = 20 GWAS).

### Effective sample size

To estimate the effective sample size of predicted GWAS summary statistics, we computed for each SNP *j* its effective sample size 𝑛_𝑗_ following Mallard and colleagues^57^ (Eq 4), where 𝑍 is the Z-statistic, 𝛽 is the predicted effect size and 𝑀𝐴𝐹 is the minor allele frequency for that SNP.

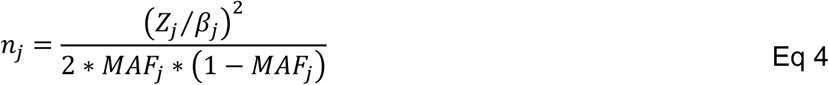

### Biological annotation of associated loci and mapped genes

#### Identification of genome-wide associated loci

SNP associations of the social behaviour GWAS were identified and annotated using Functional Mapping and Annotation of genetic associations software^22^ (FUMA, v1.6.0). A genome-wide threshold of 5x10^-8^ was applied to identify lead SNPs. Note that no further correction for multiple testing was required, given that GWAS summary statistics were derived from the same meta-regression model for each SNP. Independent GWAS signals were identified if they had a pairwise LD r²≤0.6 using the 1000 Genomes Phase 3 European reference panel^53^. These SNPs were used in subsequent steps for gene mapping and gene prioritisation. Lead SNPs were identified based on a pairwise LD-r²≤0.1 within an LD block distance of 250kb. Furthermore, the Major Histocompatibility Region was excluded from the analysis due to its extended LD structure.

#### Functional annotation and gene mapping

To map the independent SNPs to genes, both positional and eQTL mappings were used, as implemented in FUMA^22^. Positional mapping was carried out using ANNOVAR annotations within FUMA and using a window of 10kb. For eQTL mapping, SNPs were mapped to genes for SNPs that are likely to affect the expression of those genes up to 1Mb, either in blood or brain tissues. Following default settings in FUMA, only SNP-gene pairs were considered, applying a false discovery rate of <5%. Identified variants were annotated for their deleteriousness and mapped genes for their intolerance to functional mutations. Chromatin interaction mapping was also carried out; however, mapped genes were only used for gene-based analysis (GENE2FUNC in FUMA) and not for gene identification. Additionally, we conducted a look-up of GWA loci for known genes and known phenotypic associations. All analyses are described in full in Supplementary Note 4.

#### Gene expression patterns, tissue specificity and molecular functions

Genes mapped by positional and eQTL mappings, as well as chromatin interaction mapping, were annotated using GENE2FUNC (version 1.3.5) within FUMA for gene expression patterns and tissue specificity (differentially expressed genes, DEG) and overrepresentation in specific gene sets.

Additionally, we conducted a look-up of whole-brain cortical expression signatures of the genes with potential biological implications using the Allen Human Brain Atlas (AHBA; https://human.brain-map.org, N=6 post-mortem brains, aged 24-58 years)^58^. Moreover, we extracted gene expression data from the spatio-temporal gene expression data set available from the BrainSpan Atlas of the Developing Human Brain (http://www.brainspan.org/)^59^. This data covers transcriptome profiles of 16 different brain regions, spanning a time frame from embryonic development to late adulthood, for both males and females. All analyses are described in full in Supplementary Note 4.

### Gene-based, gene-set and gene-property analyses

Gene-based, gene-set, and gene-property analyses were conducted with MAGMA (version 1.08)^60^. SNPs were mapped to protein-coding genes from Ensembl (build 85) using a window of 1kb upstream and 0kb downstream. A genome-wide, gene-based significance threshold was set to 2.62x10^-6^ (0.05/19,039 protein-coding genes tested). The multiple testing adjusted p-value threshold for gene-set analysis was set to 1.05x10^-5^ (0.05/4,767 gene sets) and the multiple testing threshold for gene-property analyses was set to 4.03x10^-4^ (0.05/124 tissue types). A detailed description is provided in Supplementary Note 4.

### GWAS selection for FUMA and MAGMA analysis

To provide a parsimonious overview of the biological insights from the 76 discovery GWAS summary statistics, FUMA and MAGMA analyses were carried out for the strongest independent genome-wide hits only (Supplementary Figure 32). To identify the top hits, first, we retained all GWAS with at least one genome-wide signal (N_GWAS_=22). Second, across these 22 GWAS, there were a total of 38 SNPs covering 6 independent genomic loci (Table 1). For each of the 6 loci, we interrogated the GWAS with the strongest association signal run with FUMA, specifically, these were self-reported LPB at 9 years, parent-reported LPB at 4 and 6 years, parent-reported PSP at 2 and 6 years and early childhood PSP (Supplementary Figure 32).

### Conditional analyses

To confirm the independence of genome-wide hits, we conducted conditional analysis using GCTA-COJO^21^ (GCTA v1.94) on the six selected GWAS, where we conditioned the association of the lead SNP on the other SNPs that passed the genome-wide threshold (p<5x10^-8^) and were located on the same chromosome.

### Genetic architecture of social behaviour

#### SNP heritability and genetic correlation analysis

To study the overarching genetic architecture of the derived GWAS summary statistics (Supplementary Table 5), we estimated SNP-based heritability (h^2^_SNP_) and genetic correlations (r_g_) using LDSC regression (version 3)^26^ and LDSC correlation^27^, based on a Genomic-relatedness-based restricted maximum likelihood (GREML) approach. All analyses were based on HapMap 3 SNPs only, and precalculated LD scores from the European 1000 Genomes reference cohort^53^ were used.

To compute h^2^_SNP_, LDSC estimates the proportion of phenotypic variance tagged by SNPs by regressing the genome-wide X^2^-statistics on the proportion of genetic variation captured by each SNP^26^. The intercept of the regression estimates the contribution of confounding biases, such as cryptic relatedness and population stratification, and is more powerful at detecting these biases compared to the genomic control^26^ method.

To compute r_g_, LDSC estimates the extent of shared genetic influences between two traits, as captured by SNPs, by obtaining the slope of the regression of the product of the effect sizes for the two traits against their LD scores^27^. Specifically, we applied unconstrained LDSC correlation, which allows for sample overlap. The LDSC-r_g_ intercept reflects the shared sources of population stratification and the potential sample overlap between the two investigated traits^27^.

#### Genomic factor analysis across social behaviour

To study the relationships across the 66 discovery GWAS summary statistics (high-resolution: across social domain, reporter and age), we carried out a data-driven and hypothesis-free approach, following a similar pipeline as applied in our previous work^28^. The aim of this approach is to identify clusters of variables sharing genetic variation. Specifically, we carried out a combination^28^ of principal component analysis (PCA), EFA and CFA, where the CFA model is fitted with genomicSEM^29^ software, an approach that has been described in detail in Supplementary Note 5.

### Polygenic prediction of social behavioural differences

PGS for LPB and PSP across reporters and ages were constructed in four independent cohorts (N_ind_=16,305), described in the Supplementary Notes (Supplementary Note 7), including extensive validation analyses in the Millennium Cohort Study (MCS, ages 3-17, 18 phenotypes)^30^ as well as further analyses in two European-ancestry cohorts and one African-ancestry cohort (Supplementary Note 6, Supplementary Table 16). Detailed information on phenotype and genetic data, consent and ethical approval for each of these cohorts is available in Supplementary Note 6 and Supplementary Table 16.

PGS were calculated with PRS-CS^61^ (version June 2021) for European-ancestry cohorts and PRS-CSx software^62^ (version November 2024) for the admixed African-ancestry cohort. PRS-CS is a method that applies a continuous-shrinkage parameter to adjust the effect sizes of the genetic markers for local LD patterns. As a reference LD panel, we used the 1000 Genomes Phase 3 imputed genetic data. The default software settings were used: auto-option for phi, gamma-gamma prior parameter a is 1, gamma-gamma prior parameter b is 0.5, 1000 Markov Chain Monte Carlo iterations, 500 burn-in iterations, and a Markov chain thinning factor of 5. Using the re-estimated effect sizes, PGS were generated with PLINK^63^ (v1.9) and, subsequently, Z-standardised.

PGS were used as predictors in linear models of quantitative outcomes, which were adjusted for covariates (age, sex, and the first ten ancestry-informative principal components). The variation explained by the PGS was captured as incremental-R^2^, which represents the difference in R^2^ between a model with PGS and covariates and a baseline model with only the covariates.

To correct for multiple testing, we estimated the number of independent measures using Matrix Spectral Decomposition^64^ based on the phenotypic correlation matrix. This resulted in a threshold of p<0.0037 (0.05/14 independent measures) in the MCS, and for other cohorts as detailed in the Supplementary Notes (Supplementary Note 7).

### Genetic correlation analyses with population-based traits and NNCs

To study the genetic overlap of social behaviour with other population-based traits (N=51) and clinically-defined NNCs (N=5, ADHD, autism, MDD, schizophrenia and BD), we computed bivariate genetic correlations using LDSC software^27^. Population-based traits included a broad range of developmental and adult phenotypes, including externalising-related, internalising-related, social, wellbeing, personality, cognition and EA, language, and anthropometric and birth outcomes (Supplementary Table 23). Respective summary statistics were obtained from existing publications (Supplementary Table 23) or the UK biobank GWAS repository (http://www.nealelab.is/uk-biobank/). The multiple testing threshold (Bonferroni adjusted) was set to p<8.89x10^-4^ (0.05/56 traits). Within this study, we refer to the strength of a correlation when reporting the p-value and refer to the magnitude of the correlation when reporting the size of the correlation, which can be low (<0.3), moderate (0.3 to 0.6) or strong (>0.6).

### Genomic factor analysis across social behaviour and NNCs

In addition to genetic correlations, we studied the multivariate architecture across the five NNCs and social behaviour. Specifically, we used the meta-regression-derived medium-resolution social behaviour GWAS differentiating across social domain and developmental stages with sufficient power (h^2^_SNP_ Z-score>3, LPB_EarlyCh_, PSP_EarlyCh_, LPB_MidCh_, PSP_MidCh_, LPB_Adol_, PSP_Adol_) and clinically defined NNCs as studied for correlation analyses (Supplementary Table 23): ADHD, autism, MDD, schizophrenia and BD (Supplementary Note 8). We refrained from including further NNCs into the model to ensure computational feasibility of fitting a model across traits with differing h^2^_SNP_ estimates (h^2^_SNP_ range: 2.3%, 21%). To identify a genomic factor model, we adopted a data-driven approach^28^ using a combination of PCA, EFA and CFA, where the CFA model was fitted with genomicSEM^29^ software, as described above (Supplementary Note 5). In addition, to further characterise the identified NNC factors, we subsequently mapped EA onto the factor structure of the best-fitting CFA model Supplementary Note 8.

## DATA AVAILABILITY

**Social behaviour GWAS:** Meta-regression-derived GWAS summary statistics will be made available upon publication of the manuscript via a public data repository.

**1958 birth cohort.** Phenotype and genetic data are provided by the UK Data Service at the University of Essex after registration (https://doi.org/10.5255/UKDA-Series-2000032). Information about data access is described in the CLS Data Access Framework: https://cls.ucl.ac.uk/wp-content/uploads/2017/02/CLS_Data_Access_Framework.pdf.

**ABCD-NL**. Due to ethical considerations, the individual data from the ABCD study cannot be deposited in a public repository. Any researcher can request the data by submitting a general publication proposal form to the ABCD Project Team as outlined at https://www.amc.nl/web/abcd-studie-2/abcd-studie/voor-onderzoekers-.htm under the section “Use of Data” (Dutch: Gebruik van data), by. Our data access policy can be found at https://www.amc.nl/web/abcd-studie-2/abcd-studie/voor-onderzoekers-.htm.

**ABCD-US**. The ABCD data used in this study came from the ABCD Data Release 4.0. The raw data are available at https://nda.nih.gov/edit_collection.html?id=2573. Instructions on how to create an NDA study are available at https://nda.nih.gov/nda/tutorials/creating_an_nda_study.

**ALSPAC**. The data used are available through a fully searchable data dictionary (http://www.bristol.ac.uk/alspac/researchers/our-data/). Access to ALSPAC data can be obtained as described within the ALSPAC data access policy (http://www.bristol.ac.uk/alspac/researchers/access/).

**BREATHE**. Access to BREATHE data can be obtained by contacting.

**CATSS**. The data used from the CATSS cohort are not publicly available to protect sensitive phenotype information for participating children. Access to CATSS can be obtained by submitting a data request.

**COPSAC**. Data from the COPSAC cohort is classified as sensitive, personally identifiable data and thus cannot be made publicly available, but can potentially be made available upon request under a data transfer agreement.

**DCHS**. The Drakenstein Child Health Study is committed to the principle of data sharing. De-identified data will be made available to requesting researchers as appropriate. Requests for collaborations are welcome. More information can be found on our website http://www.paediatrics.uct.ac.za/scah/dclhs.

**ELVS**. The data used from the ELVS dataset are not readily available due to the ethical governance set in place for this study. The data are shared following application to investigators of the ELVS study. Requests to access the genetic datasets should be directed to Angela Morgan.

**FinnTwin12**. In accordance with the Finnish Biobank Act, the data used in the analysis are deposited in the Biobank of the Finnish Institute for Health and Welfare (https://thl.fi/en/web/thl-biobank/forresearchers). They are available to researchers from academia and companies after a written application and following the relevant Finnish legislation. To ensure the protection of privacy and compliance with national data protection legislation, a data use/transfer agreement is needed, the content and specific clauses of which will depend on the nature of the requested data.

**GenR**. Data from the Generation R cohort are not publicly available due to legal and ethical restrictions; however, access can be requested via the Generation R administration. Requests for Generation R data access are evaluated by the Generation R Management Team.

**GINILISA**. Due to data protection reasons, the datasets generated and/or analysed during the current study cannot be made publicly available. The datasets are available to interested researchers from the corresponding author upon reasonable request (e.g. reproducibility), provided the release is consistent with the consent given by the GINIplus and LISA study participants. Ethical approval might be obtained for the release, and a data transfer agreement from the legal department of the Helmholtz Zentrum München must be accepted.

**GLAKU**. Researchers can obtain a de-identified GLAKU dataset after having obtained approval from the GLAKU Study Board.

**HRS**. The phenotype survey data are provided by the health and retirement study (HRS) without charge, after registration. The genomic data are provided by the health and retirement study after IRB approval and approval by NCBI-dbGaP, without charge. Sensitive data, including genetic information and biomarkers, are only available through an approved data application process (Health and Retirement Study, DUA Review Committee, 426 Thompson Street, Ann Arbor, Michigan 48104-2321). By requesting access to the HRS genetic information via NCBI-dbGaP, the authors agreed that the data are only shared within the working group permitted to work with the data (St Pourcain group) (https://hrs.isr.umich.edu/sites/default/files/HRS-Genetic-Data-Access-Agreement.pdf). Information about the data access is described here: https://hrs.isr.umich.edu/data-products/.

**INMA**. INMA data (INMA-GSA and INMA-Omni1) are available by request from the INfancia y Medio Ambiente Executive Committee for researchers who meet the criteria for access to confidential data. Access to INMA data can be obtained by contacting.

**INSchool**. Raw data from INSchool are not publicly available because of limitations in ethical approvals, and the summary data will be available by contacting.

**LSAC**. Data from the LSAC cohort can be obtained by completing an online form, following instructions from the LSAC data custodian, The Australian Institute of Family Studies. Information about data access is described in https://growingupinaustralia.gov.au/data-and-documentation/accessing-lsac-data.

**MCS**. Phenotype and genetic data from the Millennium Cohort Study are provided by the UK Data Service at the University of Essex after registration (https://doi.org/10.5255/UKDA-Series-2000031).

Information about data access is described in the CLS Data Access Framework: https://cls.ucl.ac.uk/wp-content/uploads/2017/02/CLS_Data_Access_Framework.pdf.

**MoBa**. Data from the Norwegian Mother, Father and Child Cohort Study and the Medical Birth Registry of Norway used in this study are managed by the national health register holders in Norway (Norwegian Institute of Public Health). The consent given by the participants does not allow for storage of data on an individual level in repositories or journals. Researchers who want access to data sets for replication should apply through helsedata.no. Access to data sets requires approval from the Regional Committee for Medical and Health Research Ethics in Norway (REC) and an agreement with MoBa. In addition, compliance with the EU General Data Protection Regulation (GDPR) is required.

**MUSP**. MUSP data are available from a third party upon reasonable request. Contact details can be found here: https://social-science.uq.edu.au/mater-university-queensland-study-pregnancy?p=5#5.

**NFBC1986**. NFBC data are available from the University of Oulu, Infrastructure for Population Studies. Permission to use the data can be applied for research purposes via an electronic material request portal. In the use of data, we follow the EU General Data Protection Regulation (679/2016) and the Finnish Data Protection Act. The use of personal data is based on a cohort participant’s written informed consent in their latest follow-up study, which may cause limitations to its use. Please, contact the NFBC project center and visit the cohort website (www.oulu.fi/nfbc) for more information.

**NTR**. The data used from the NTR cohort are not publicly available to protect sensitive phenotype information for participating children. Access to NTR can be obtained by submitting a data request.

**The Raine Study**. Restrictions apply to the availability of data generated or analysed during this study to preserve patient confidentiality. On request (https://ross.rainestudy.org.au), details of the restrictions and any conditions under which data access is granted may be provided.

**SYS**. SYS data are available upon request and should be addressed to Dr Zdenka Pausova and Dr Tomas Paus. Further details about the protocol can be found at http://www.saguenay-youth-study.org/.

**TEDS**. Researchers can apply for access to the TEDS data through their data access mechanism (www.teds.ac.uk/researchers/teds-data-access-policy).

**TRAILS**. TRAILS data are not publicly available to researchers outside the TRAILS consortium, but can be obtained by submitting a publication proposal. For more details, please see www.trails.nl.

## CODE AVAILABILITY

This study used openly available software and codes, specifically abagen toolbox (v0.1.1, https://github.com/rmarkello/abagen), FUMA (v1.6.0, https://fuma.ctglab.nl/), GCTA-COJO (v1.94, https://yanglab.westlake.edu.cn/software/gcta/), GEMMA (https://github.com/genetics-statistics/GEMMA), multivariate GWAMA (https://github.com/baselmans/multivariate_GWAMA), LDSC (v1.0.1, https://github.com/bulik/ldsc), METAL (v2011-03-25, https://csg.sph.umich.edu/abecasis/metal/), MAGMA (v1.08, https://ctg.cncr.nl/software/magma), PLINK (v1.9, https://www.cog-genomics.org/plink/1.9/), ProbABEL (https://github.com/GenABEL-Project/ProbABEL), PRScs (Jun-04-2021, https://github.com/getian107/PRScs), PRScsx (https://github.com/getian107/PRScsx), SNPTEST (https://mathgen.stats.ox.ac.uk/genetics_software/snptest/snptest_v2.4.1.html). We used the following R packages: cerebroviz v1.0, EasyQC v9.2, fsbrain v0.5.5, genomicSEM v0.05, lavaan v0.6-10, LDlinkR v1.4.0, Matrix v1.7-0, metafor v3.0-2, nFactors v2.4.1. SFARI (https://gene.sfari.org/database/human-gene/), Scripts used in this study will be made available via GitLab upon publication.

## ACKNOWLEDGEMENTS

We thank all the children, twins, families, and participants who took part and are taking part in the 29 cohorts whose data contributed to the GWAS meta-analyses or follow-up analysis for making this work possible. We also thank all employees of the cohorts included in this study. This project was embedded within the EAGLE Consortium, and follow-up analyses were also embedded in the R2D2 Consortium. Cohort-specific acknowledgements and funding information can be found in Supplementary Note 12. In addition, we thank all cohorts and researchers who made their summary statistics available to us. This includes the GenLang Consortium, Social Science Genetic Association Consortium, Early Growth Genetics Consortium, EAGLE Consortium, and the Psychiatric Genomics Consortium. We would like to extend our thanks and acknowledge the input of the R2D2-MH Adult Co-Creation Group for their support and contribution to the design of the study materials. We thank Wolfgang Viechtbauer for helpful discussions about conducting meta-regressions using metafor software.

L.d.H., F.S., D.A.G.A., S.J., E. Verhoef, S.v.d.B., S. Scatolin, E.E., N.Y.T.N., S.E.F., and B.S.P. were supported by the Max Planck Society. S.J., A.K., A.J., E.B., H.S., M.H.B., T.R., C.E., S.B., T.B., and B.S.P. were supported by the R2D2-MH, which has been funded by Horizon Europe (grant agreement no. 101057385), by UK Research and Innovation (UKRI) under the UK government’s Horizon Europe funding guarantee [grant no. 10039383] and by the Swiss State Secretariat for Education, Research and Innovation (SERI) under contract number 22.00277. B.S.P. is also supported by the Radboud University Donders RSF 2025. T.G.M.V. was supported by ZonMW (TOP 40–00812–98–11010). C.A.M.C. is supported by the Horizon Europe Research and Innovation Programme (FAMILY, grant agreement no. 101057529, also to A.N.; HappyMums, grant agreement no. 101057390) and the European Research Council (TEMPO; grant agreement no. 101039672, also to A.N.). A.N. and H.T. are supported by a grant from the Dutch Ministry of Education, Culture, and Science and the Netherlands Organisation for Scientific Research (NWO grant no. 024.001.003, Consortium on Individual Development). The work of H.T. is further supported by the European Union’s Horizon 2020 research and innovation program (Contract grant no. 633595, DynaHealth) and an NWO-VICI grant (NWO-ZonMW: 016.VICI.170.200). M.L. was supported by the scholarship from the China Scholarship Council (201706990036). This work of M.L. was further supported by the Ter Meulen Grant of the Royal Netherlands Academy of Arts and Sciences (KNAW). J.L. has received research support from the Strategic Research Council (SRC) established within the Academy of Finland (decision no. 352700). S.A. acknowledges her Miguel Servet contract (CP22/00026) awarded by the Instituto de Salud Carlos III and co-funded by the European Union Fund (Fondo Social Europeo Plus, FSE+). A. Havdahl was supported by funding from the Research Council of Norway (RCN) (#274611, #336085) and the South-Eastern Norway Regional Health Authority (HSØ) (#2020022 & #2018059). E.C.C. was supported by funding from the RCN (#274611) and HSØ (#2021045) and is a member of the MRC Integrative Epidemiology Unit at the University of Bristol which is supported by the Medical Research Council and the University of Bristol (MC_UU_00032/1). T.R-K. was supported by funding from the RCN (#274611). E.Y. was supported by funding from the RCN (#288083 and #336078) and the European Union (Grant agreement No. 101045526 and No. 818425). P.C. has received funding from the University of Oulu and the Academy of Finland Profi6 (decision number AF 336449). S. Sebert acknowledges funding support by The Horizon Europe framework programme (STAGE [Grant N° 101137146]; IHEN [Grant N°101137317], OBELISK Grant [N° 101080465], OBCT Grant N°[101080250], TRIGGER [Grant N°101057739]), The Horizon 2020 framework programme (LongITools [Grant N°874739], EarlyCause [Grant N°848158]) and the research council of Finland [Grant N°356888]. M-R.J. acknowledges funding support by The Horizon 2020 framework programme (LongITools [Grant N°874739], EarlyCause [Grant N°848158]) and by the MRC Centre for Environment and Health funded by the Medical Research Council, UK (MR/S019669/1). D.I.B. acknowledges the Royal Netherlands Academy of Science Professor Award (PAH/6635). A. Ronald received funding from the Simons Foundation (Award ID: 724306). K. Rimfeld is supported by a Sir Henry Wellcome Postdoctoral Fellowship (213514/Z/18/Z). N.V-T. is supported by the Spanish Ministry of Science and Innovation - State Research Agency grant RYC2022-038136-I, cofunded by the European Union FSE+, and grant PID2022-143106OA-I00, cofunded by the European Union FEDER. Additionally, N.V-T. is supported by the William H. Gates Sr. Fellowship from the Alzheimer’s Disease Data Initiative. TEDS is supported by a programme Grant to R.P. from the UK Medical Research Council (Grant Nos. MR/V012878/1 and previously MR/M021475/1), with additional support from the US National Institutes of Health (Grant No. AG046938). The research leading to these results has also received funding from the European Research Council under the European Union’s Seventh Framework Programme (FP7/2007-2013)/ grant agreement n° 602768. V.W. is supported by funding from the Simons Foundation for Autism Research Initiative, the Wellcome Trust (214322\Z\18\Z), Horizon-Europe R2D2-MH (grant agreement number 101057385), and UKRI (10063472). J.K.B. has been supported by the EU-AIMS (European Autism Interventions) and AIMS-2-TRIALS programmes which receive support from Innovative Medicines Initiative Joint Undertaking Grant No. 115300 and 777394, the resources of which are composed of financial contributions from the European Union’s FP7 and Horizon2020 Programmes, and from the European Federation of Pharmaceutical Industries and Associations (EFPIA) companies’ in-kind contributions, and AUTISM SPEAKS, Autistica and SFARI; and by the Horizon2020 supported programme CANDY Grant No. 847818).

The funders had no role in the design of the study; in the collection, analyses, or interpretation of data; in the writing of the manuscript, or in the decision to publish the results. This publication is the work of the authors, who serve as the guarantors for the contents of this paper. Views and opinions expressed are those of the authors only and do not necessarily reflect those of the European Union, the European Research Council Executive Agency, or the other funders. Neither the European Union nor the granting authority can be held responsible for them.

## COMPETING INTERESTS

H.L. reports receiving grants from Shire Pharmaceuticals; personal fees from and serving as a speaker for Medice, Shire/Takeda Pharmaceuticals and Evolan Pharma AB; all outside the submitted work. H.L. is editor-in-chief of JCPP Advances. J.A.R-Q. was on the speakers’ bureau and/or acted as a consultant for Biogen, Idorsia, Casen-Recordati, Janssen-Cilag, Novartis, Takeda, Bial, Sincrolab, Neuraxpharm, Novartis, BMS, Medice, Rubió, Uriach, Technofarma and Raffo in the last 3 years. J.A.R-Q. also received travel awards (air tickets + hotel) for taking part in psychiatric meetings from Idorsia, Janssen-Cilag, Rubió, Takeda, Bial and Medice. The Department of Psychiatry, chaired by J.A.R-Q., received unrestricted educational and research support from the following companies in the last 3 years: Exeltis, Idorsia, Janssen-Cilag, Neuraxpharm, Oryzon, Roche, Probitas and Rubió. E.D.R. has served as a speaker for Shire Sweden AB, a Takeda Pharmaceutical Company, outside of this work. S.B. discloses that he has in the last 3 years acted as a consultant or lecturer for Medice, Takeda, and LinusBio. S.B. receives royalties for textbooks and diagnostic tools from Hogrefe, UTB, Ernst Reinhardt, Kohlhammer, and Liber. S.B. is a partner in NeuroSupportSolutions International AB.

All other authors declare no conflict of interest.

