## Supplementary Figures for "Genome-wide analysis of social behaviour in context: a meta-regression approach across social domains, reporters and developmental stages"

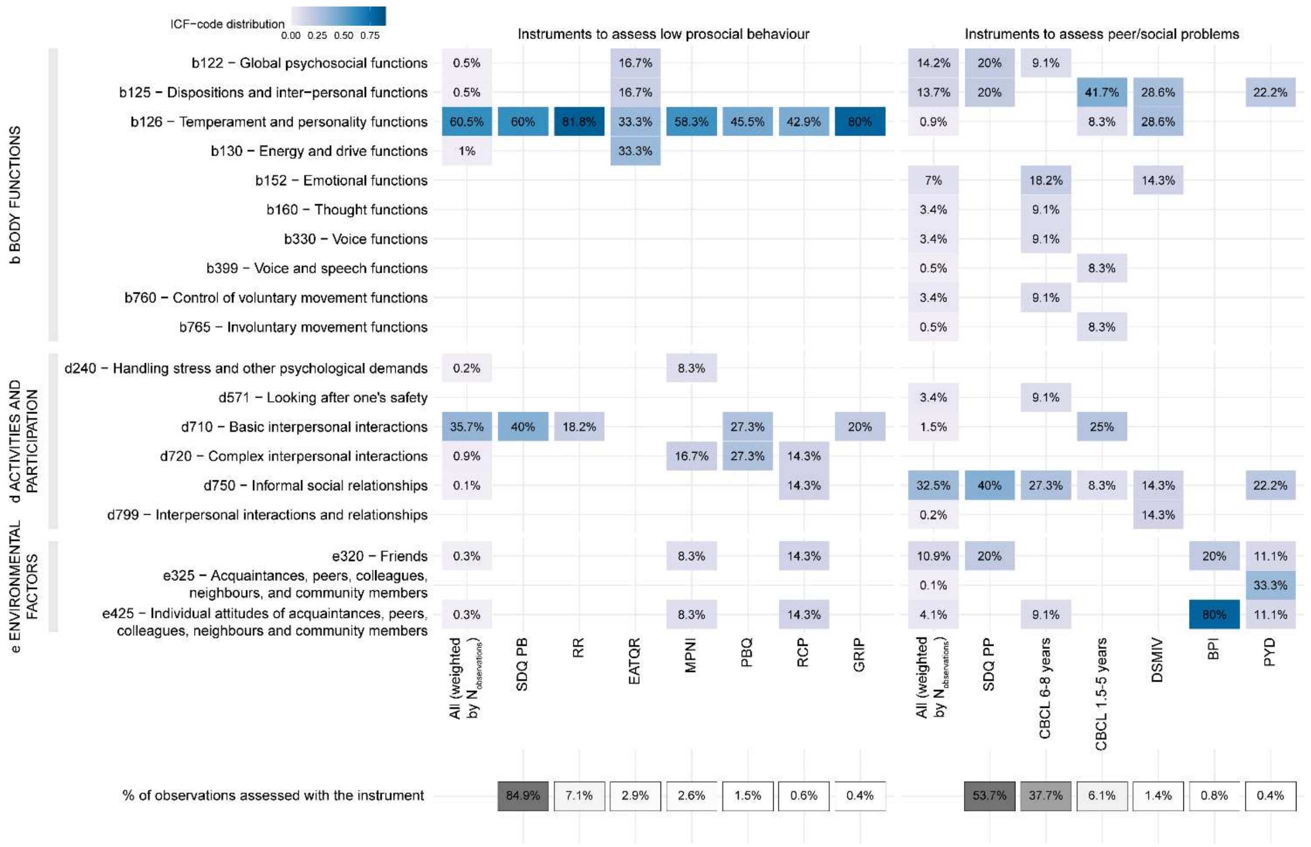

**Supplementary Figure 1.** ICF alignment of social behaviour measures

The y-axis shows the different codes from the ICF-CY grouped according to the ICF category. The x-axis shows for low prosocial behaviour (left) and peer/social problems (right) the distribution of the ICF codes across the questionnaires used to assess each social domain, as well as the overall contribution weighted by $N_{observations}$.

Abbreviations: ICF (WHO's International Classification of Functioning, Disability and Health, Children & Youth Version)

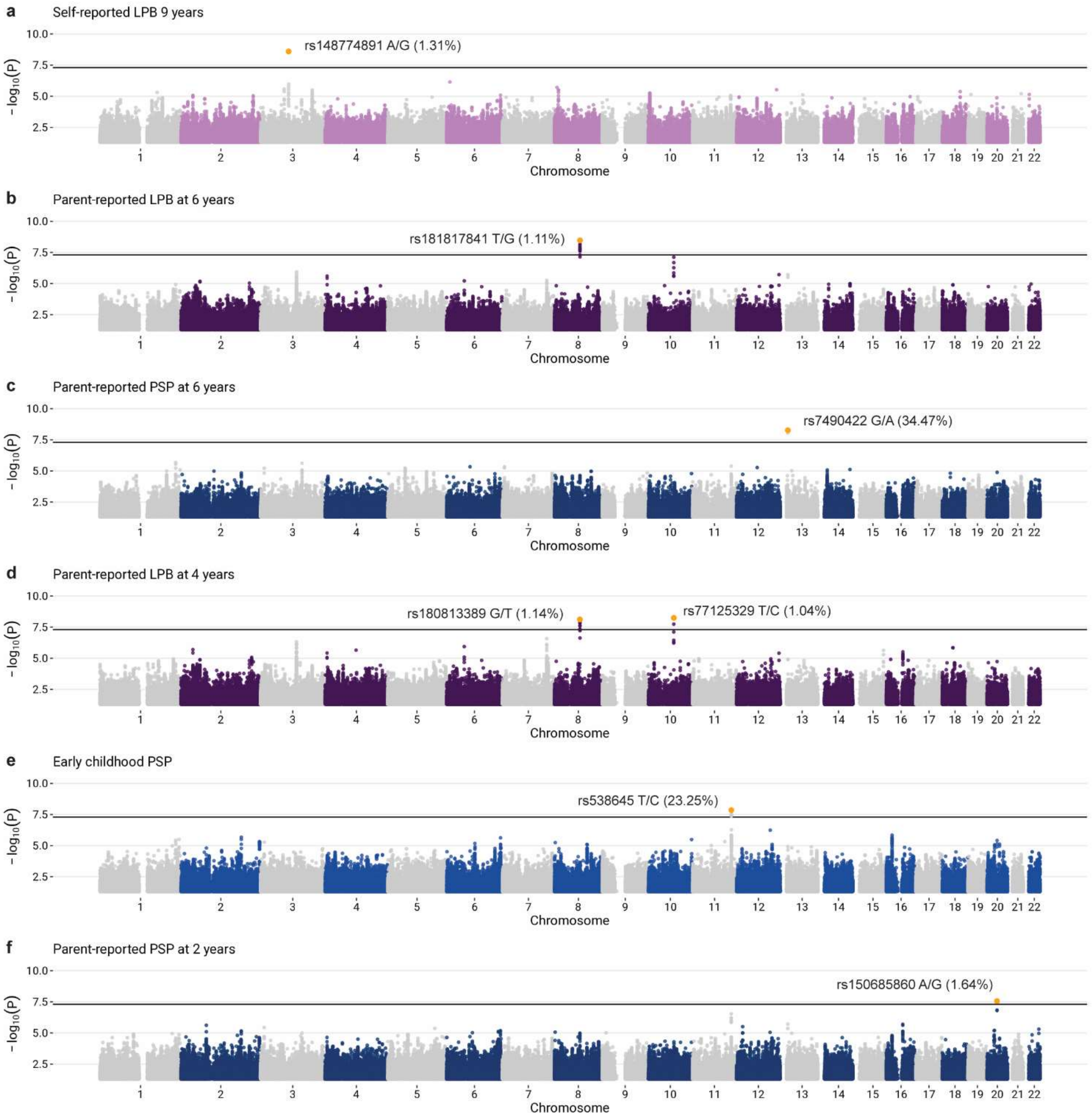

**Supplementary Figure 2.** Manhattan plots

Each dot represents a SNP. The -log10(p) is shown on the y-axis, while the x-axis indicates its genomic coordinates. A genome-wide significance threshold was set to $5x10^{-8}$ (0.05/1,000,000) and indicated by the solid horizontal black line. For each GWAS, the SNPs passing this threshold are annotated with a yellow dot along with their alleles and their minor allele frequency.

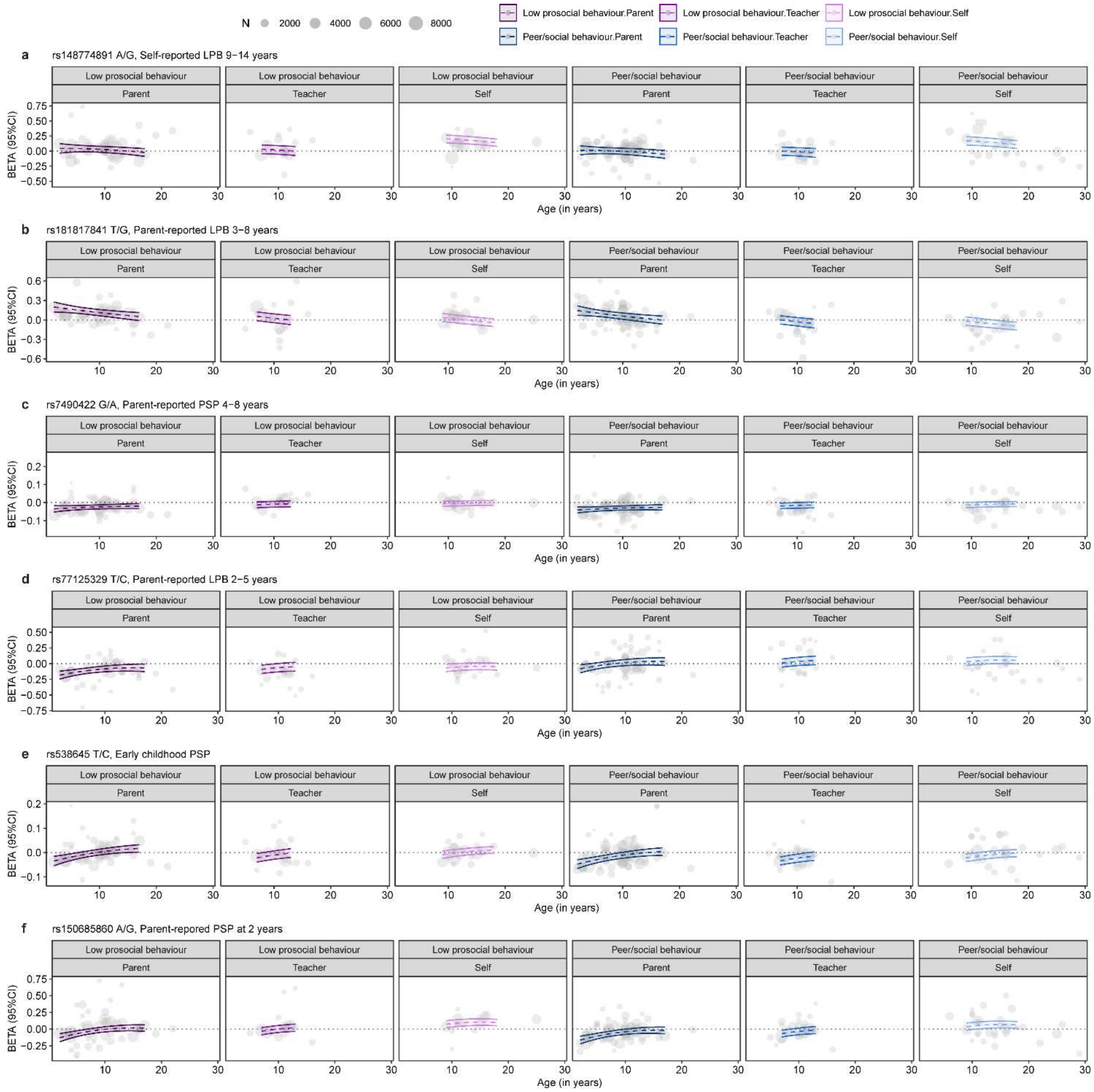

**Supplementary Figure 3.** Predicted effect sizes for genome-wide associated loci

The dashed lines represent the predicted effect size (y-axis) across age (x-axis), and the coloured area represents the associated 95% CI for lead SNPs of genome-wide associated loci. These effect sizes were estimated using random-effects meta-regression. Grey dots indicate the effect size for each SNP in the cohort-level GWAS, with the size of each dot proportional to the sample size. Data is presented across age, stratifying by social domain and reporter.

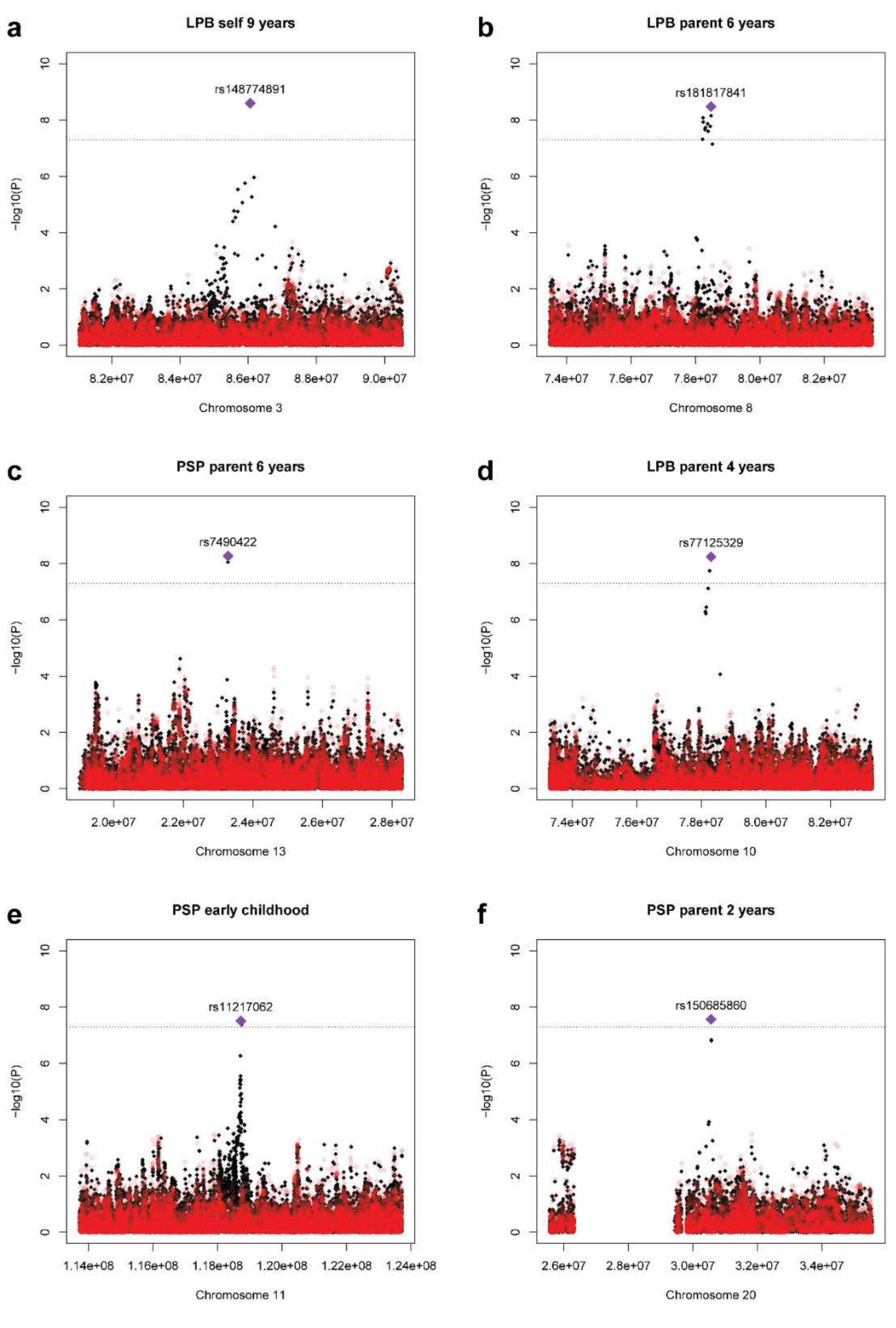

**Supplementary Figure 4.** Conditional and joint analysis

GCTA-COJO analysis was run for each of the selected GWAS and genomic loci: (**a**) Self-reported LPB at 9 years, (**b**) Parent-reported LPB at 6 years, (**c**) Parent-reported PSP at 6 years, (**d**) Parent-reported LPB at 4 years, (**e**) Early childhood PSP and (**f**) Parent-reported PSP at 2 years. Black dots indicate the original -log10(p) of the GWAS, red dots indicate the conditional -log10(p), and the lead SNP is indicated by a purple rhomb. Dashed grey line indicates the genome-wide significance threshold after Bonferroni correction: $p=5x10^{-8}$.

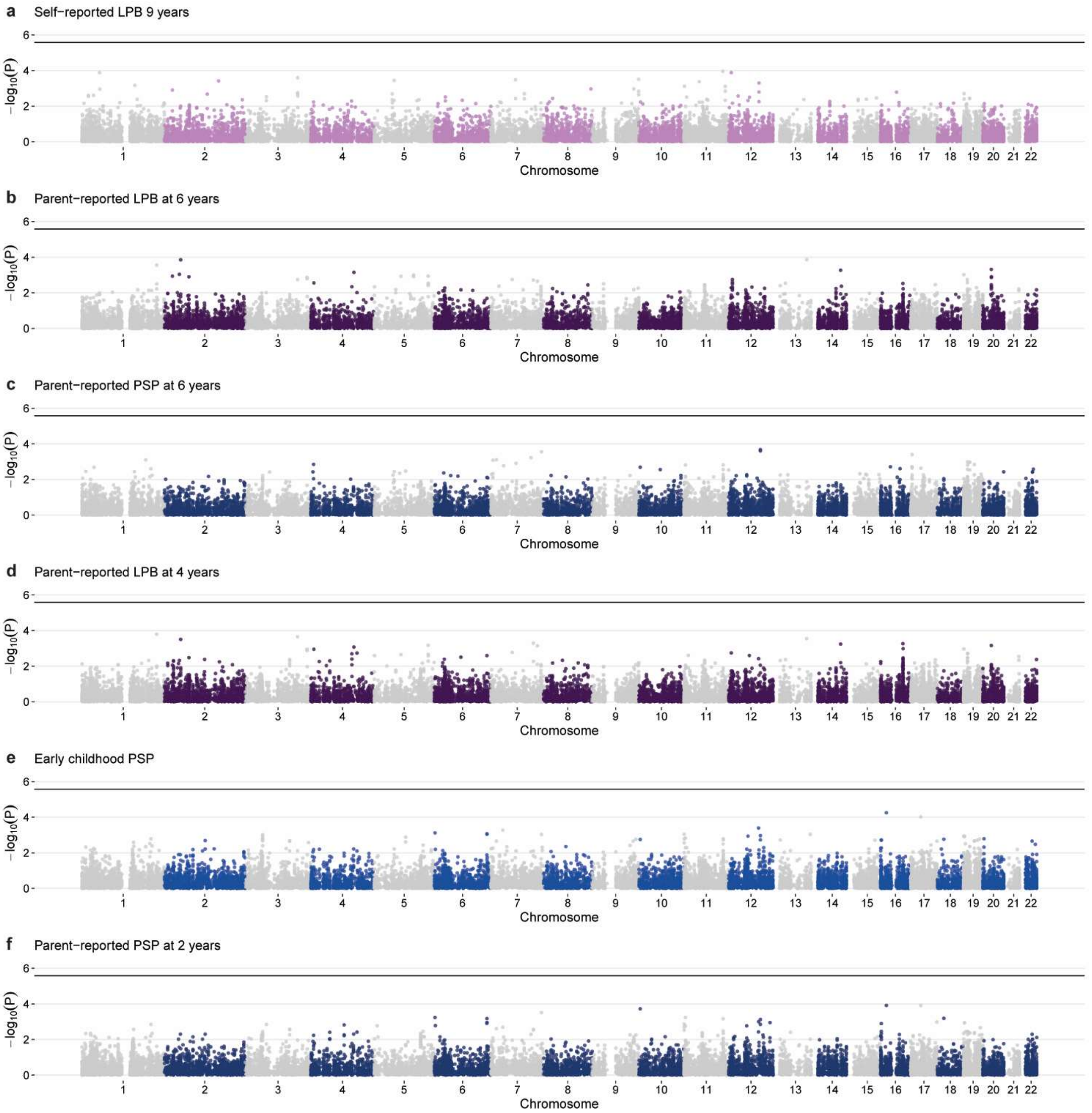

**Supplementary Figure 5.** Gene-based Manhattan plots

Each dot represents a gene. The -log10(p) is shown on the y-axis, while the x-axis indicates its genomic coordinates. A genome-wide gene-based significance threshold was set to $2.62 \times 10^{-6}$ (0.05/19,039 protein-coding genes tested) and indicated by the solid horizontal black line.

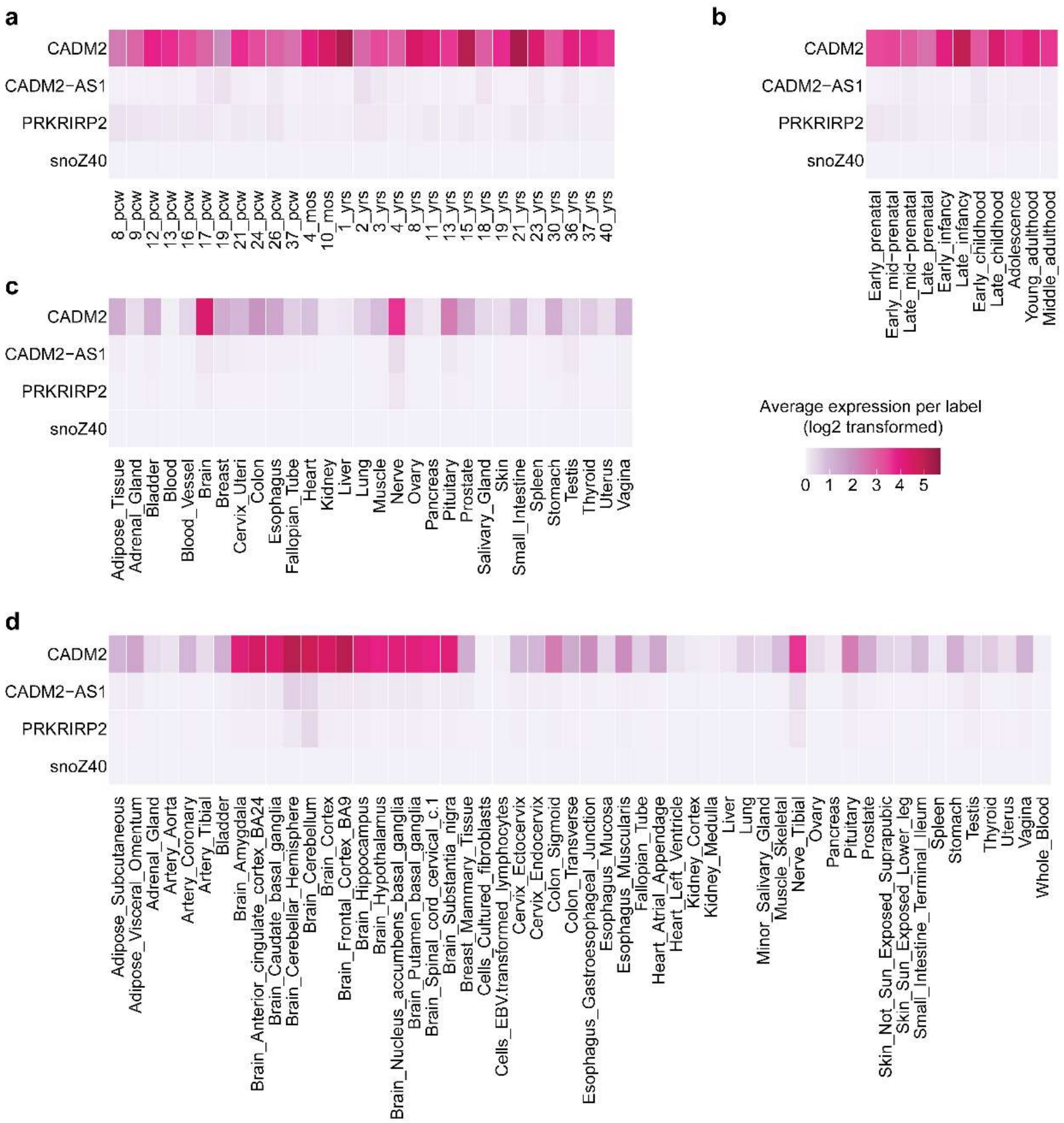

**Supplementary Figure 6.** Gene expression heatmaps across BrainSpan ages and GTEx tissues for genes mapped to locus rs148774891 on chromosome 3

(**a**) 29 BrainSpan ages, (**b**) 11 BrainSpan developmental stages, (**c**) 30 broad GTEx tissue types and (**d**) 54 specific GTEx tissue types. Average expression (log2 transformed) of the gene in each label (i.e. BrainSpan developmental stage or GTEx tissue) is shown, where darker values indicate higher expression compared to cells with lighter values. Expression value is expressed as RPKM (Reads per kilobase per million) for BrainSpan and as TPM (transcripts per million) for GTEx.

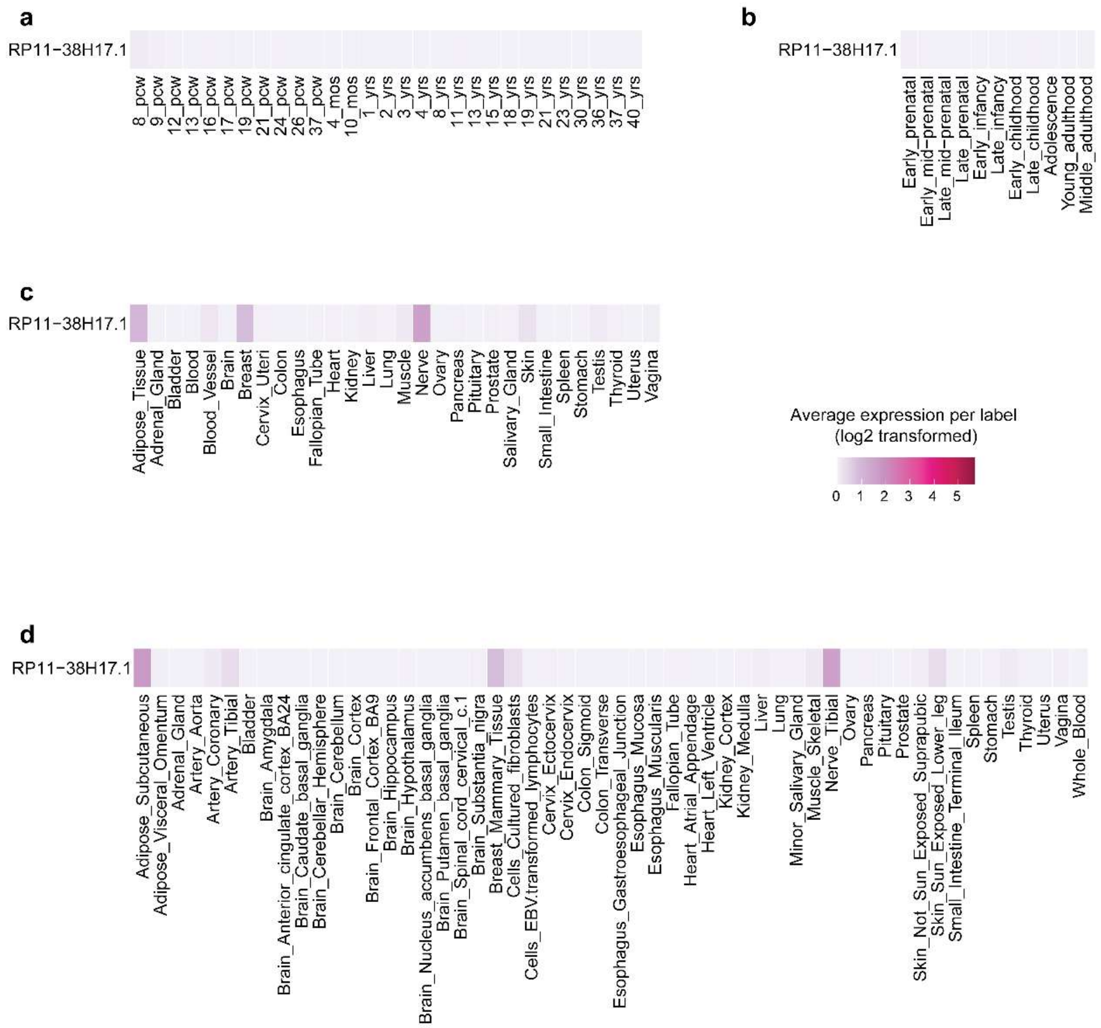

**Supplementary Figure 7.** Gene expression heatmaps across BrainSpan ages and GTEx tissues for genes mapped to locus rs181817841 on chromosome 8

(**a**) 29 BrainSpan ages, (**b**) 11 BrainSpan developmental stages, (**c**) 30 broad GTEx tissue types and (**d**) 54 specific GTEx tissue types. Average expression (log2 transformed) of the gene in each label (i.e. BrainSpan developmental stage or GTEx tissue) is shown, where darker values indicate higher expression compared to cells with lighter values. Expression value is expressed as RPKM (Reads per kilobase per million) for BrainSpan and as TPM (transcripts per million) for GTEx.

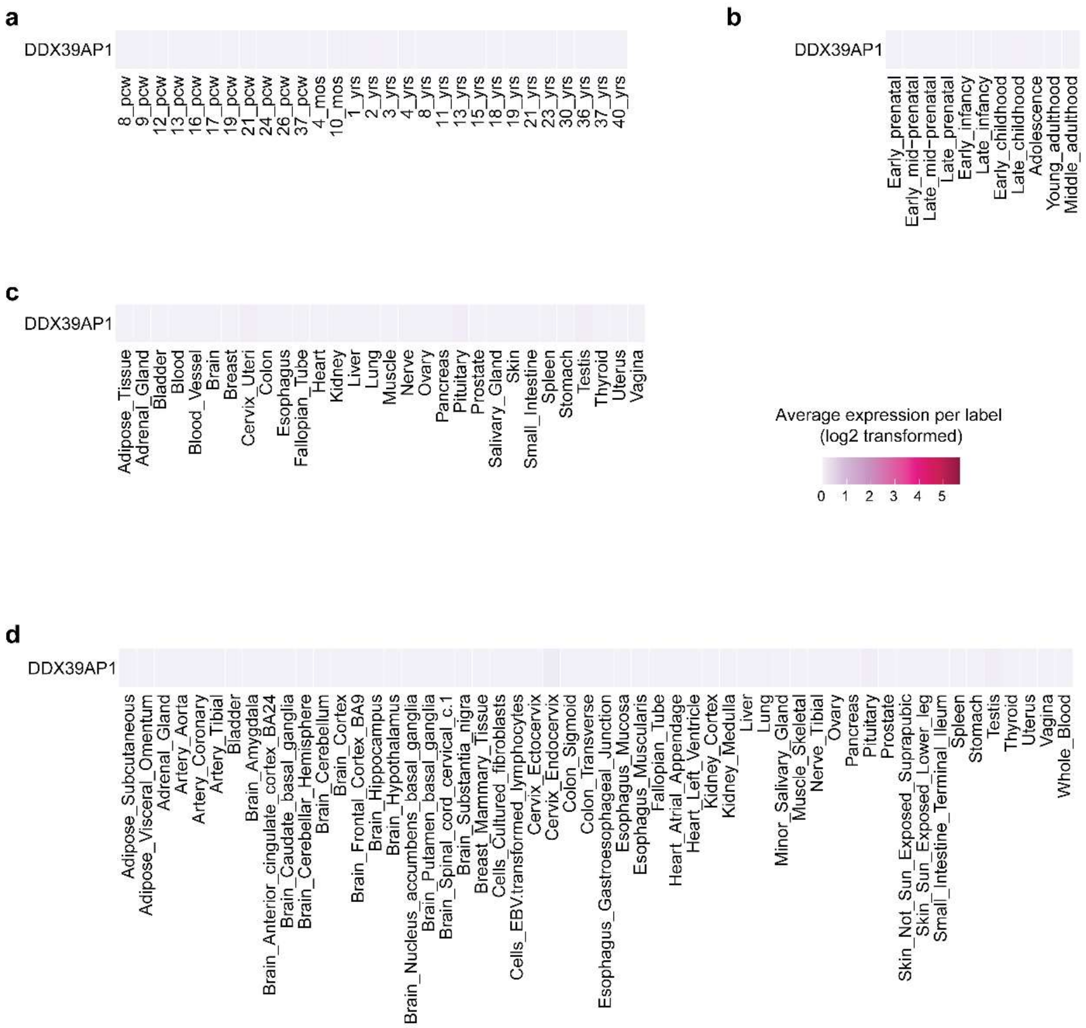

**Supplementary Figure 8.** Gene expression heatmaps across BrainSpan ages and GTEx tissues for genes mapped to locus rs7490422 on chromosome 13

(**a**) 29 BrainSpan ages, (**b**) 11 BrainSpan developmental stages, (**c**) 30 broad GTEx tissue types and (**d**) 54 specific GTEx tissue types. Average expression (log2 transformed) of the gene in each label (i.e. BrainSpan developmental stage or GTEx tissue) is shown, where darker values indicate higher expression compared to cells with lighter values. Expression value is expressed as RPKM (Reads per kilobase per million) for BrainSpan and as TPM (transcripts per million) for GTEx.

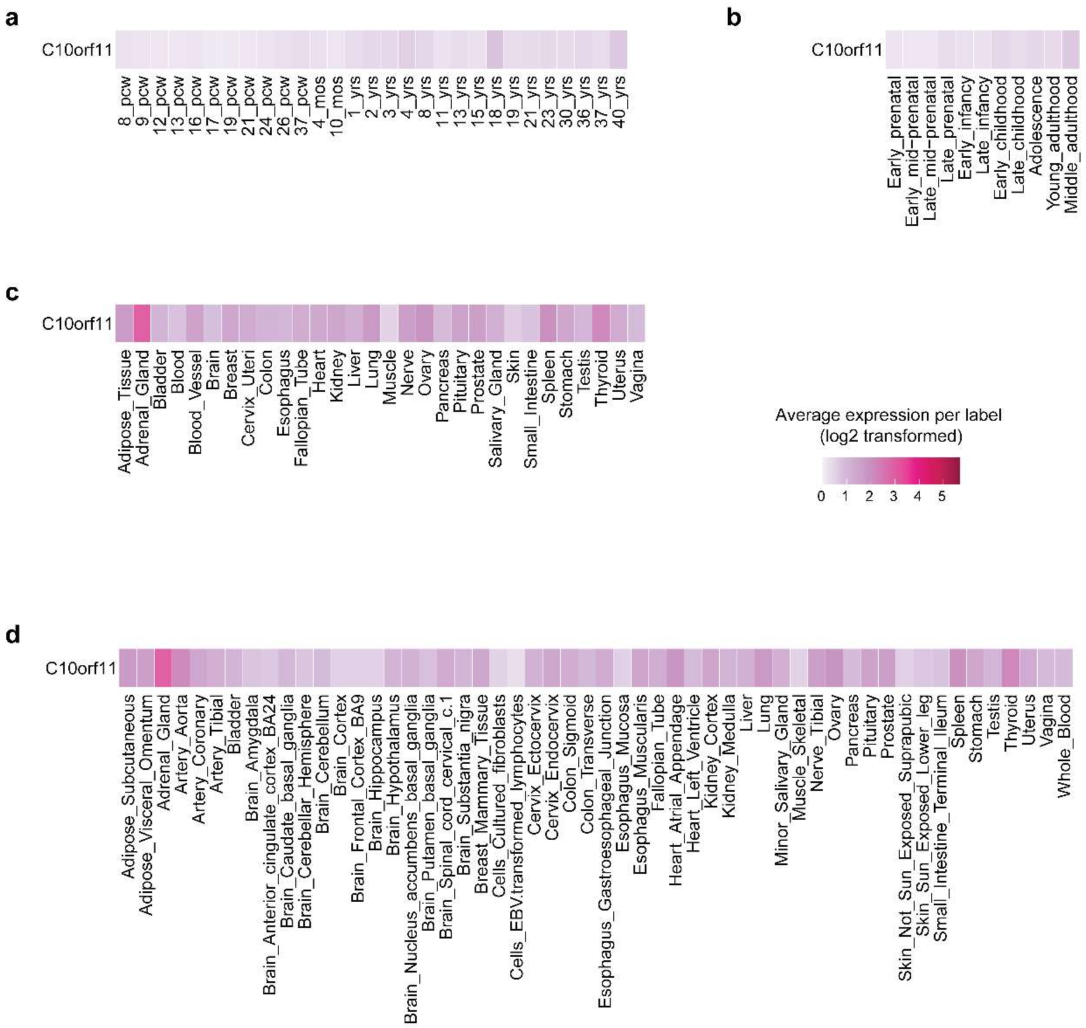

**Supplementary Figure 9.** Gene expression heatmaps across BrainSpan ages and GTEx tissues for genes mapped to locus rs77125329 on chromosome 10

(**a**) 29 BrainSpan ages, (**b**) 11 BrainSpan developmental stages, (**c**) 30 broad GTEx tissue types and (**d**) 54 specific GTEx tissue types. Average expression (log2 transformed) of the gene in each label (i.e. BrainSpan developmental stage or GTEx tissue) is shown, where darker values indicate higher expression compared to cells with lighter values. Expression value is expressed as RPKM (Reads per kilobase per million) for BrainSpan and as TPM (transcripts per million) for GTEx.

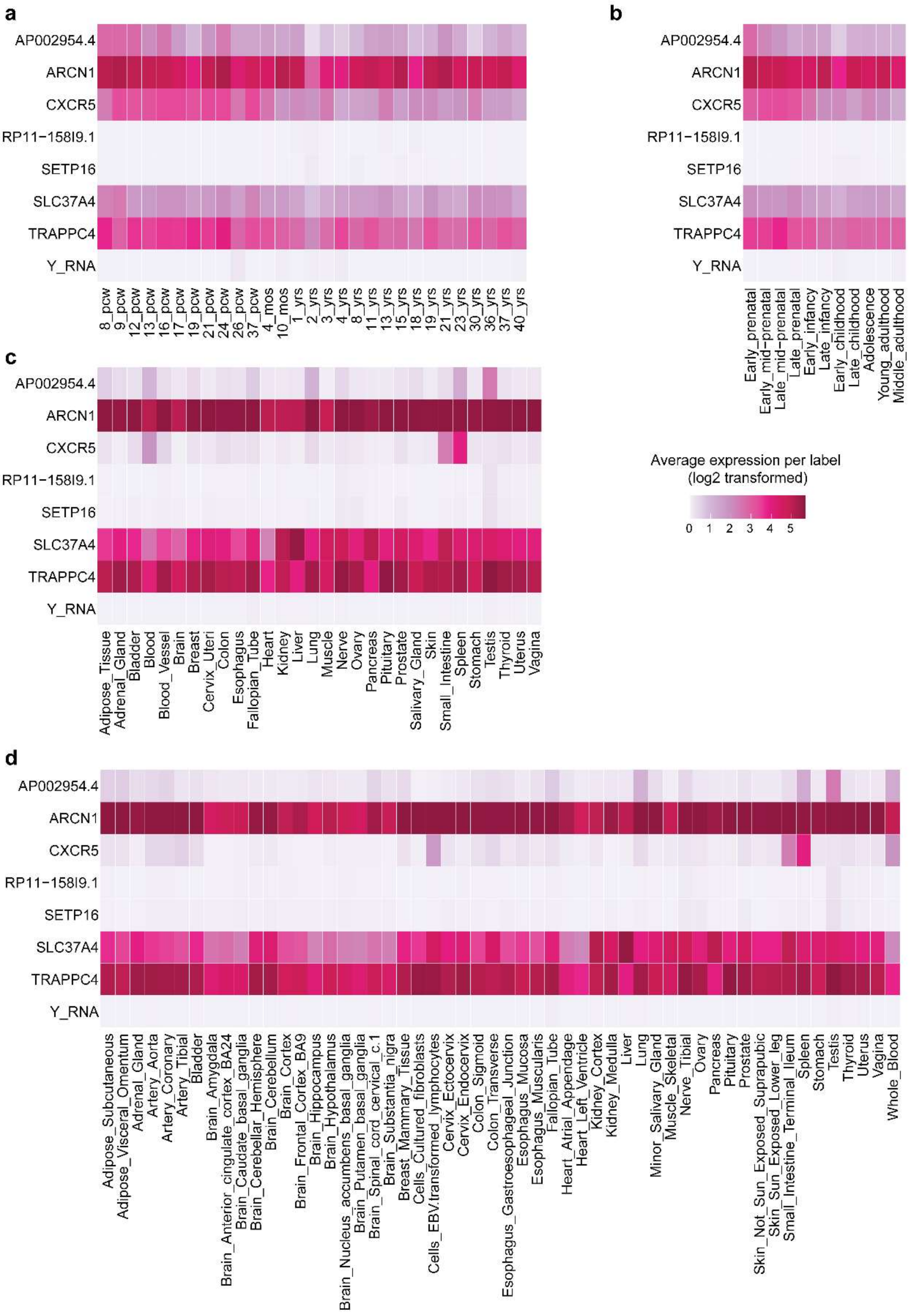

**Supplementary Figure 10.** Gene expression heatmaps across BrainSpan ages and GTEx tissues for genes mapped to locus rs538645 on chromosome 11

(**a**) 29 BrainSpan ages, (**b**) 11 BrainSpan developmental stages, (**c**) 30 broad GTEx tissue types and (**d**) 54 specific GTEx tissue types. Average expression (log2 transformed) of the gene in each label (i.e. BrainSpan developmental stage or GTEx tissue) is shown, where darker values indicate higher expression compared to cells with lighter values. Expression value is expressed as RPKM (Reads per kilobase per million) for BrainSpan and as TPM (transcripts per million) for GTEx.

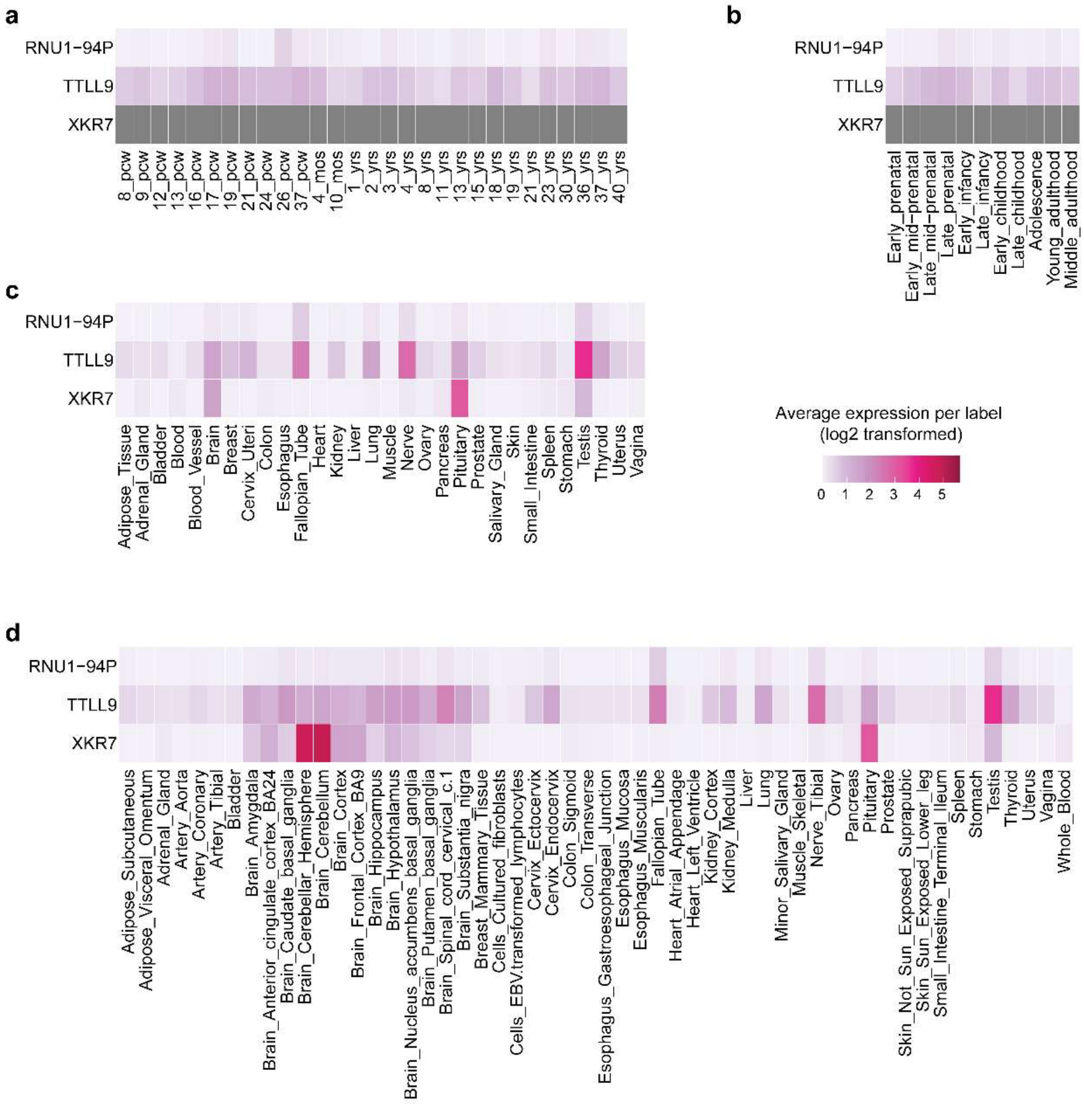

**Supplementary Figure 11.** Gene expression heatmaps across BrainSpan ages and GTEx tissues for genes mapped to locus rs150685860 on chromosome 20

(**a**) 29 BrainSpan ages, (**b**) 11 BrainSpan developmental stages, (**c**) 30 broad GTEx tissue types and (**d**) 54 specific GTEx tissue types. Average expression (log2 transformed) of the gene in each label (i.e. BrainSpan developmental stage or GTEx tissue) is shown, where darker values indicate higher expression compared to cells with lighter values. Expression value is expressed as RPKM (Reads per kilobase per million) for BrainSpan and as TPM (transcripts per million) for GTEx.

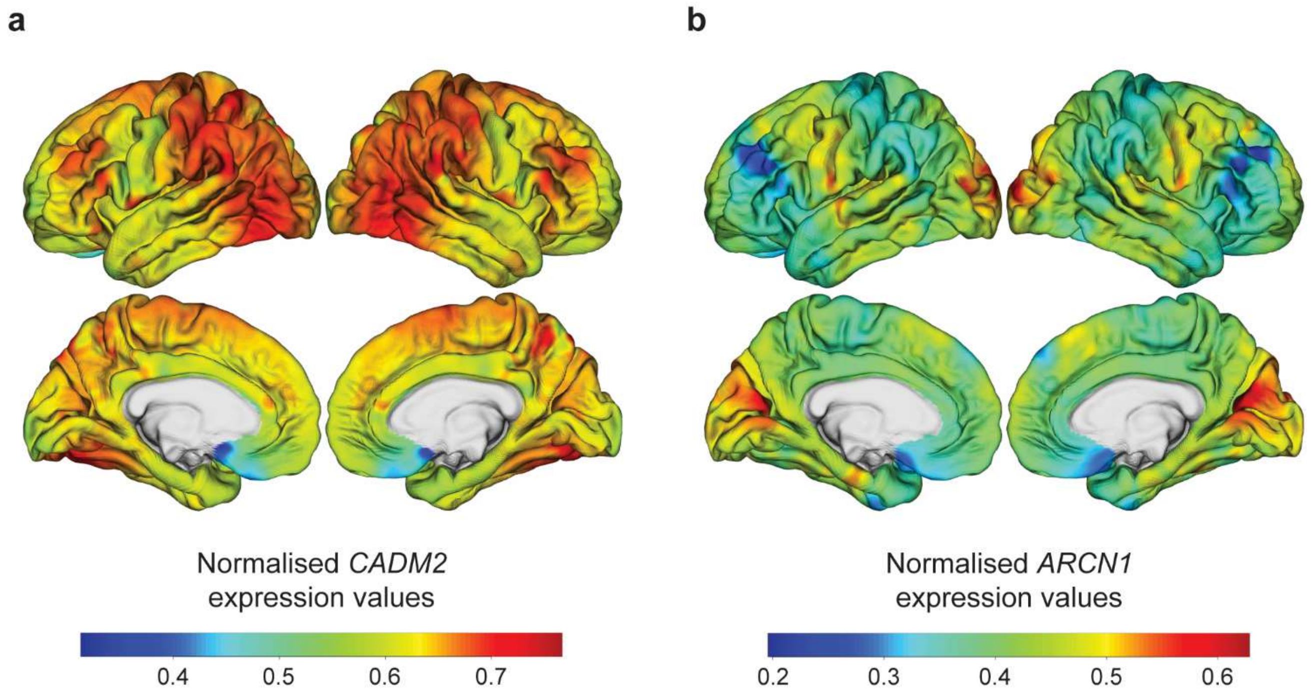

**Supplementary Figure 12.** Whole-brain cortical expression signatures of genes with potential biological implications.

Vertex-wise cortical expression of (**a**) CADM2 and (**b**) ARCN1 with standardised expression values. Whole-brain cortical expression signatures were derived from the Allen Human Brain Atlas (AHBA; https://human.brain-map.org), and samples were assigned to the FreeSurfer *fsaverage6* standard-space surface template, spatially interpolated and smoothed to create a gene-by-vertex matrix of standardised mRNA expression across the entire cortical surface.

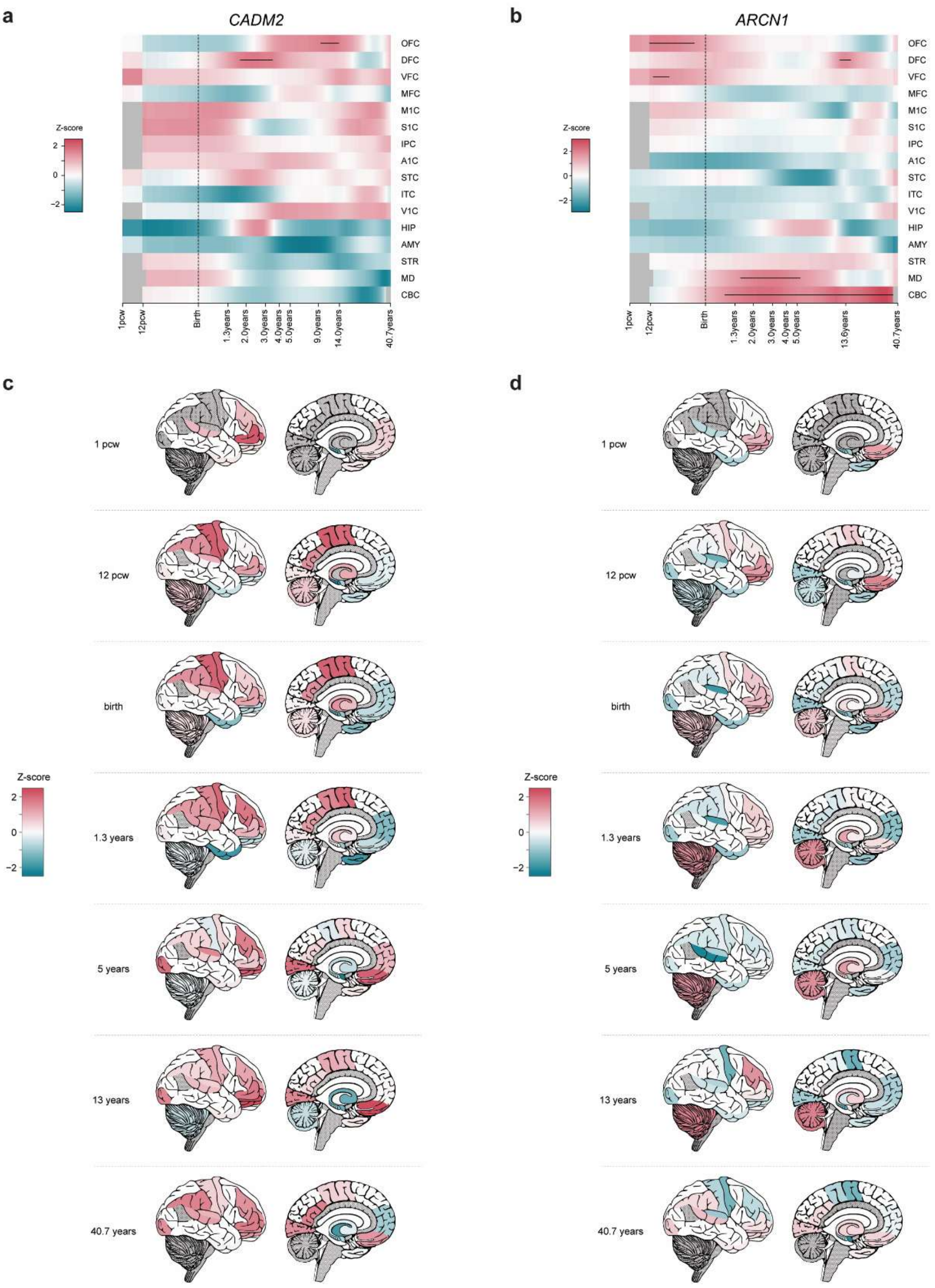

**Supplementary Figure 13.** Spatiotemporal gene expression patterns of genes with potential biological implications.

Heatmap of spatiotemporal expression profiles of **(a)** *CADM2* and **(b)** *ARCN*1 using BrainSpan data. The heatmap on the x-axis shows the developmental time frame, and the y-axis shows the different brain regions. The colour of the heatmap represents the gene expression values transformed into z-scores. Z-scores above 1.65 are indicated with a black line. The colour range of dark cyan to moderate red shows a range of lowest to highest expression values, respectively. Illustrations of brain regions corresponding to the expression data for **(c)** *CADM2* and **(d)** *ARCN1*. Z-scores of gene expression values across brain regions are visualised as sagittal views (left) and exterior views (right), ranging from 1-week post conception (top) to 40.7 years old (bottom).

Abbreviations. OFC: Orbital prefrontal cortex; DFC: Dorsolateral prefrontal cortex; VFC: Ventrolateral prefrontal cortex; MFC: Medial prefrontal cortex; M1C: Primary motor (M1) cortex; S1C: Primary somatosensory (S1) cortex; IPC: Posterior inferior parietal cortex; A1C: Primary auditory (A1) cortex, STC: Superior temporal cortex; ITC: Inferior temporal cortex; V1C: Primary visual (V1) cortex; HIP: Hippocampus; AMY: Amygdala; STR: Striatum; MD: Mediodorsal nucleus of the thalamus; CBC: Cerebellar cortex.

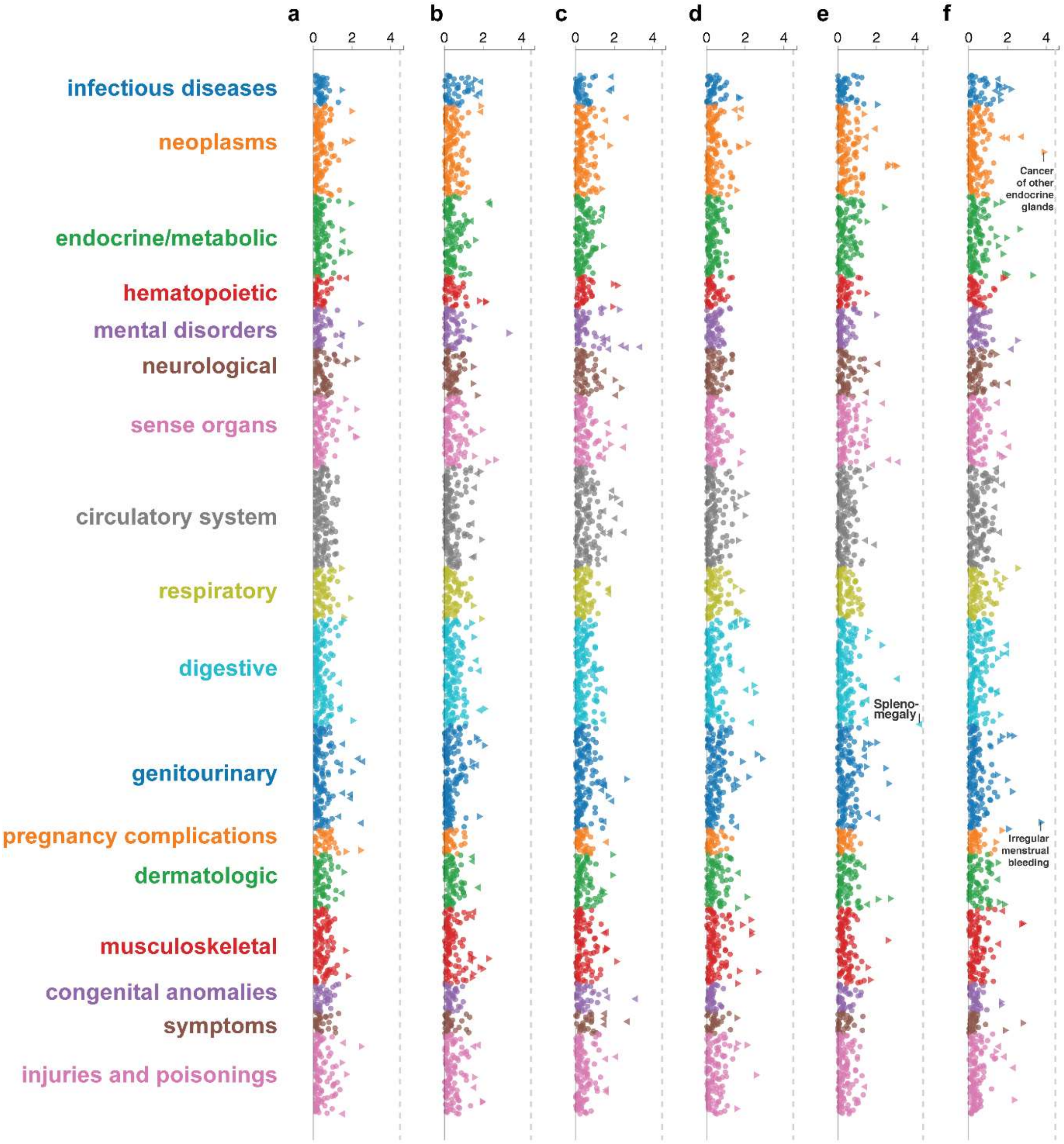

**Supplementary Figure 14.** PheWeb screen of lead SNPs

A PheWeb screen (https://pheweb.org/UKB-TOPMed/) for the genome-wide loci (**a**) rs148774891, (**b**) rs181817841, (**c**) rs7490422, (**d**) rs77125329, (**e**) rs538645, (**f**) rs150685860, was carried out for 1400 UKBiobank PheWAS codes and 57 million TOPMed-imputed variants in 400k white British individuals.

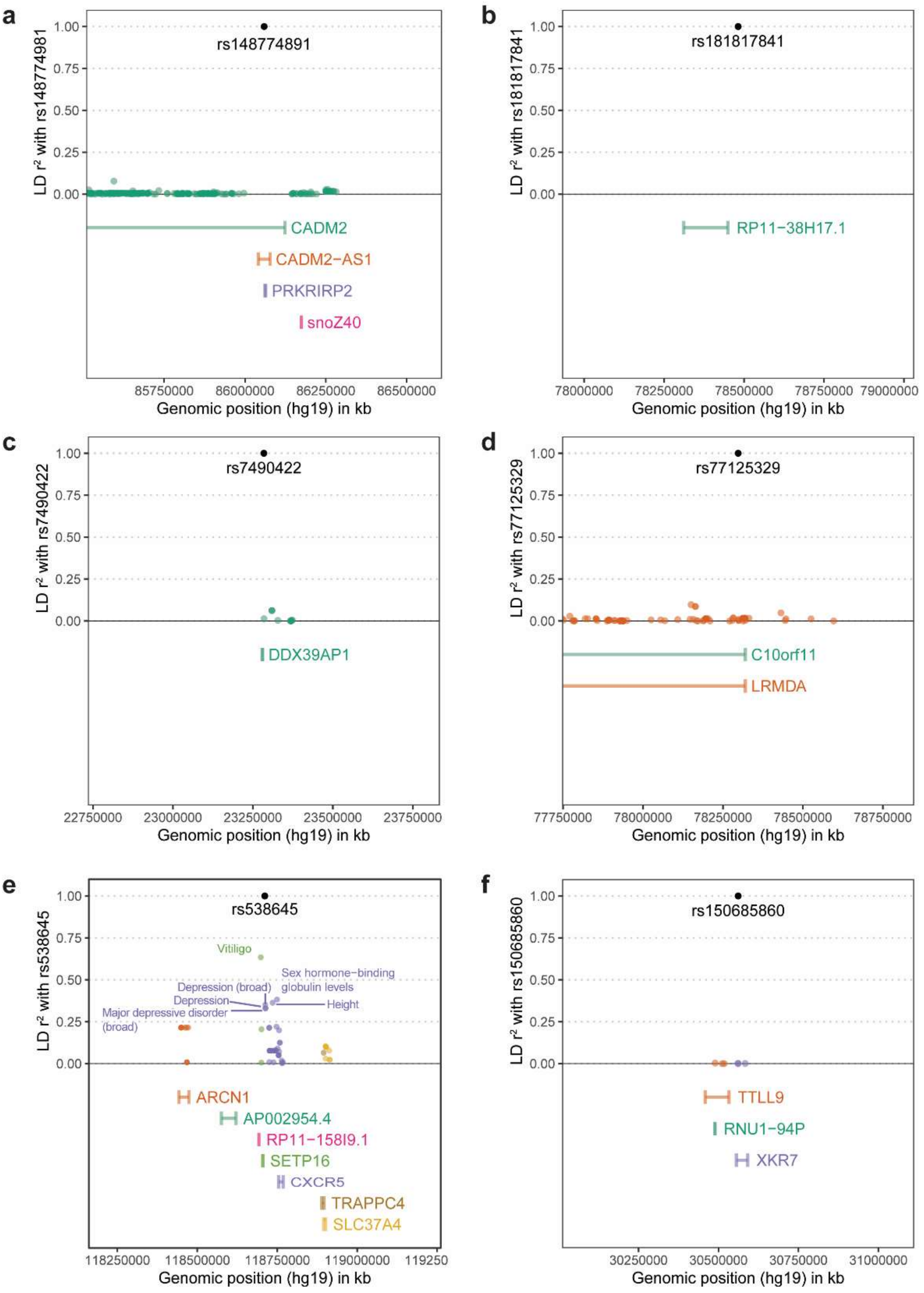

**Supplementary Figure 15.** GWAS Catalog screen of lead SNPs

The GWAS Catalog database was screened for previously reported loci within 500kb up- and downstream of the identified lead SNP, including mapped genes for (**a**) rs148774891, (**b**) rs181817841, (**c**) rs740422, (**d**) rs77125329, (**e**) rs538645, and (**f**) rs150685860. LD-$r^2$ with the lead SNP is reported on the y-axis, and the SNP's genomic location (hg19 build) is reported on the x-axis. The lead SNP is shown in black, and the SNPs reported in the GWAS Catalog are reported with the colour of their associated gene.

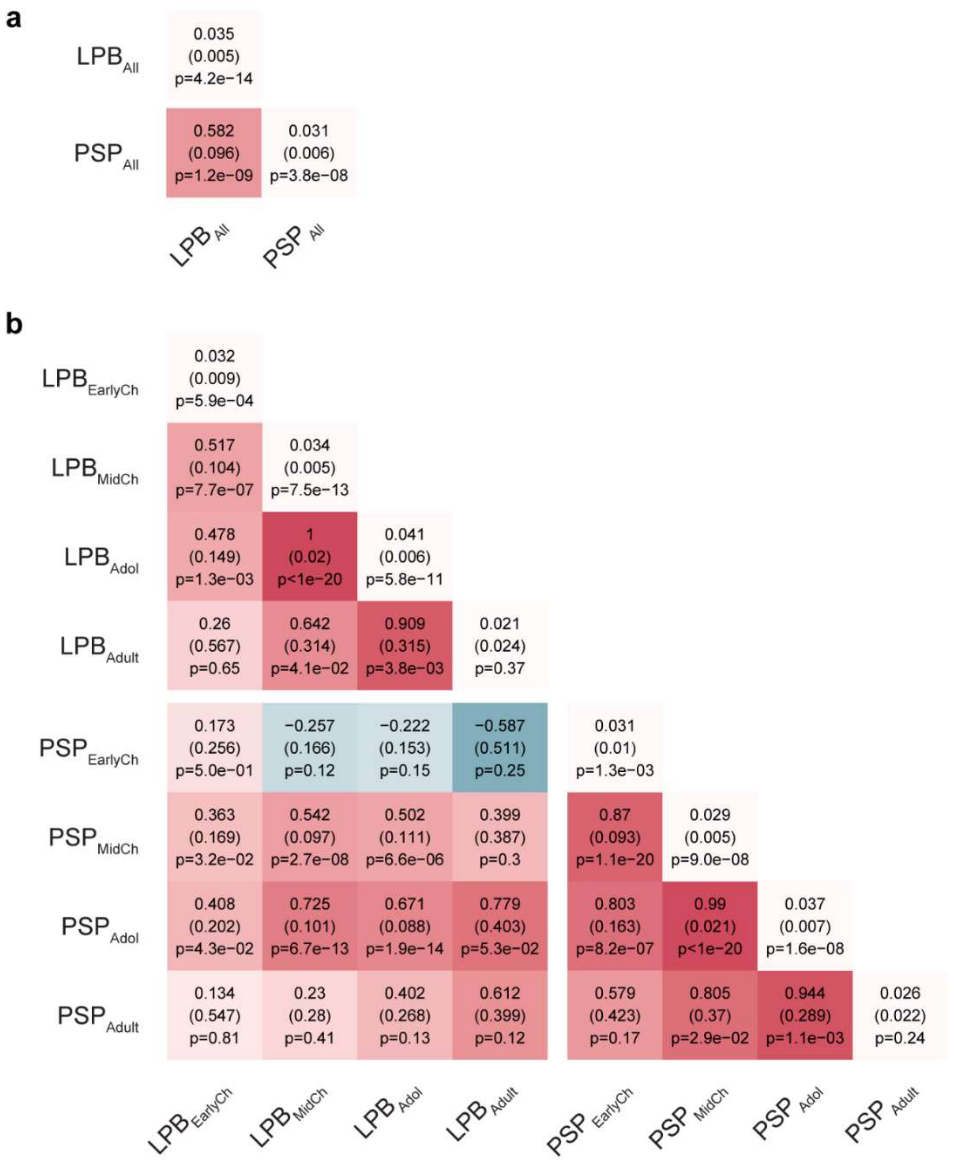

**Supplementary Figure 16.** LDSC heritability and genetic correlations for low-resolution and medium-resolution social behaviour GWAS

(**a**) LDSC $h^2_{SNP}$ and LDSC $r_g$ for low-resolution GWAS. (**b**) LDSC $h^2_{SNP}$ and LDSC $r_g$ for medium-resolution GWAS. Medium-resolution GWAS, including PSP and LPB GWAS across four developmental stages: early childhood (0-5 years), mid childhood (6-11 years), adolescence (12-17 years) and young adulthood (18-30 years). Diagonal values indicate LDSC $h^2_{SNP}$, and off-diagonal values indicate LDSC $r_g$, as estimated with linkage disequilibrium score (LDSC) regression and correlation, respectively.

Abbreviations: LPB (Low prosocial behaviour), $LPB_{EarlyCh}$ (Early childhood LPB), $LPB_{MidCh}$ (Mid childhood LPB), $LPB_{Adol}$ (Adolescence LPB), $PSP_{Adult}$ (Young adulthood PSP), PSP (Peer and social problems), $PSP_{EarlyCh}$ (Early childhood PSP), $PSP_{MidCh}$ (Mid childhood PSP), $PSP_{Adol}$ (Adolescence PSP), $PSP_{Adult}$ (Young adulthood PSP)

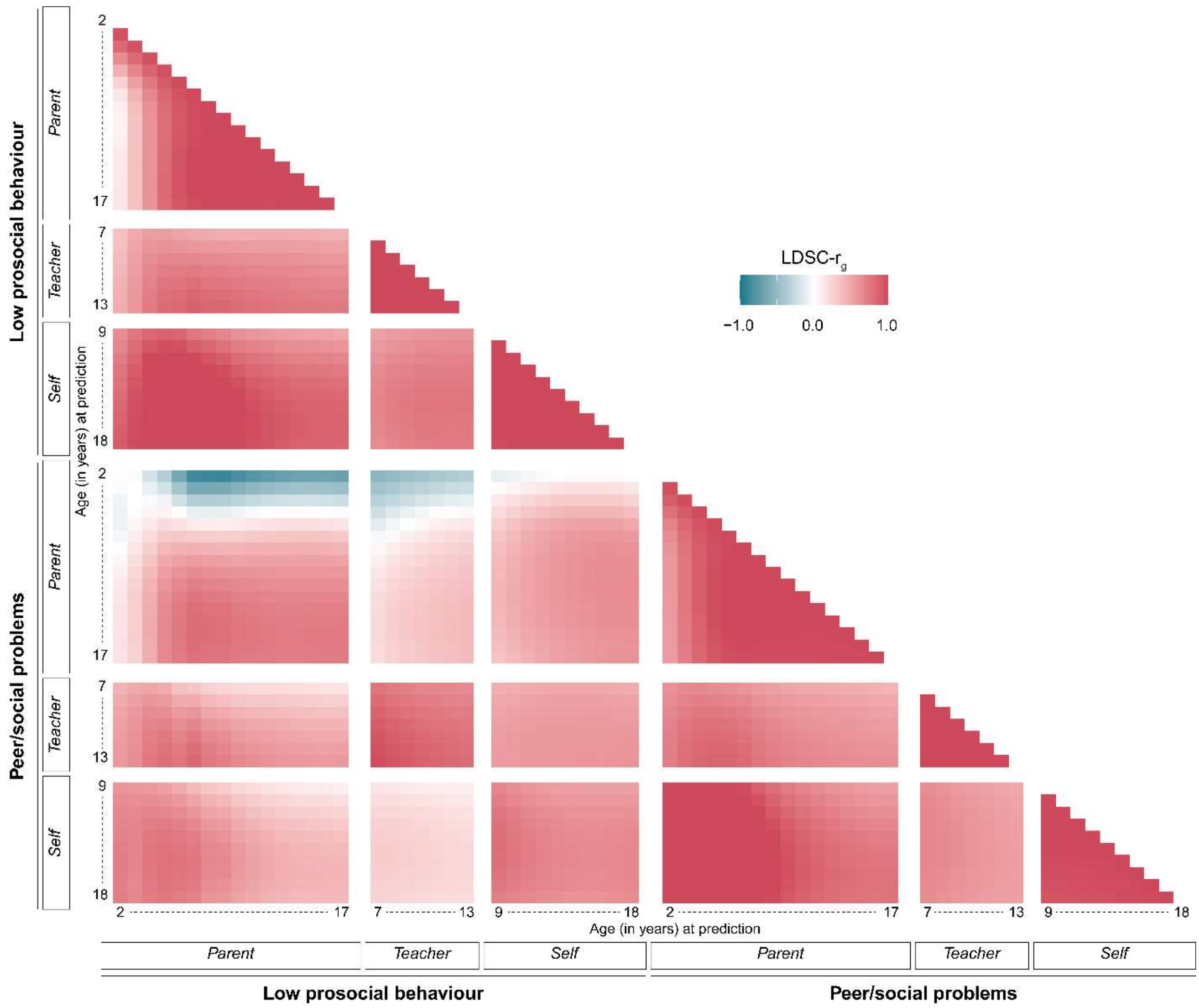

**Supplementary Figure 17.** LDSC genetic correlation matrix for high-resolution social behaviour GWAS across social domain, reporter, and age

Derived high-resolution social behaviour GWAS were restricted to the age ranges that had data on at least two cohorts or that had data on one cohort but with a sample size above 2,000. Parent-reported LPB and PSP GWAS were derived from 2 to 17 years (16 ages x 2 domains = 32 GWAS). Teacher-reported LPB and PSP GWAS were derived from 7 to 13 years (7 ages x 2 domains = 14 GWAS). For self-reported LPB and PSP GWAS, the range was set from 9 to 18 years (10 ages x 2 domains = 20 GWAS) (Methods). Genetic correlations were estimated with the linkage disequilibrium score (LDSC) correlation.

Abbreviations: LPB (Low prosocial behaviour), PSP (Peer and social problems)

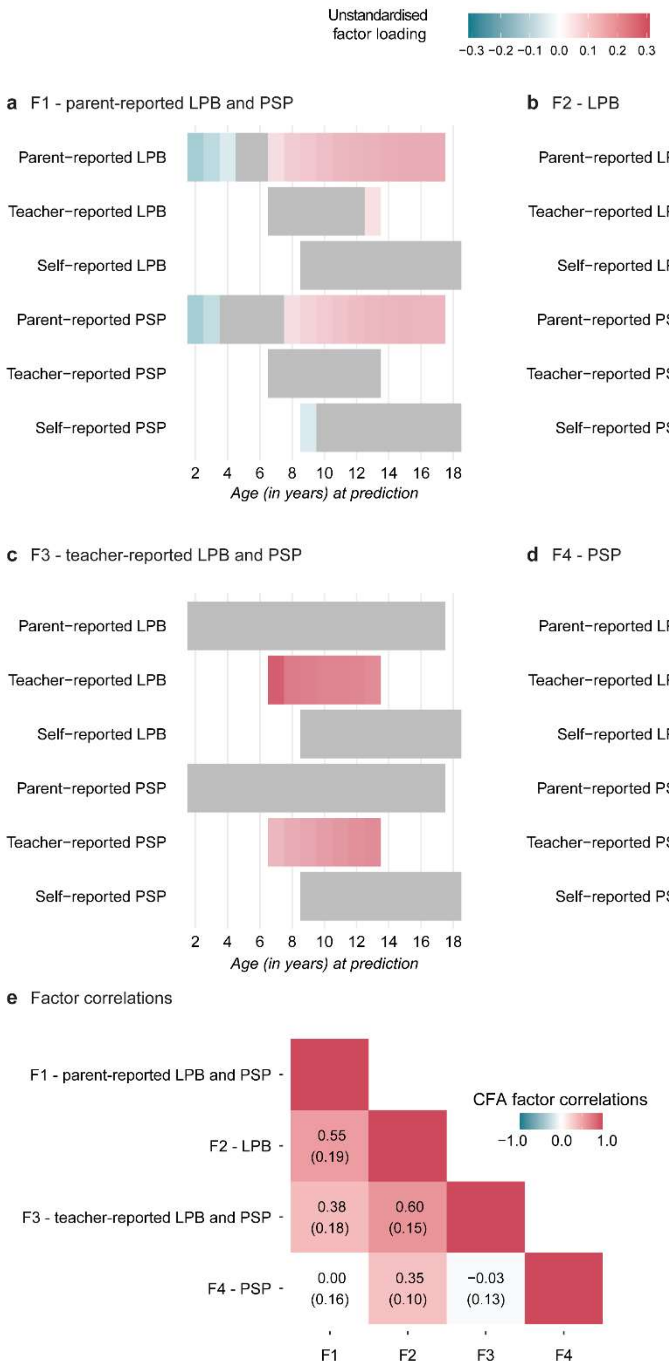

**Supplementary Figure 18.** Factor loadings of the genetic CFA model of social behaviour

Factor loadings, as estimated from an LDSC genetic covariance matrix across 66 high-resolution GWAS predictions, for (**a**) F1 – parent-reported LPB and PSP, (**b**) F2 - LPB, (**c**) F3 – teacher-reported LPB and PSP, and (**d**) F4 -PSP. The colour of the link between the factors (upper part of the plot) and the predicted GWAS summary statistics (lower part) represents the strength of the unstandardized factor loading. Factor loadings that were not estimated are shown in grey. (e) Correlations between the identified factors. Estimates are shown with their corresponding standard errors. Model fit parameters are shown in Supplementary Table 14, and standardised and unstandardised parameter estimates are shown in Supplementary Table 15. The genetic covariance was estimated with the linkage disequilibrium score (LDSC) correlation.

Abbreviations: EFA (Exploratory factor analysis), LPB (Low prosocial behaviour), PSP (Peer and social problems)

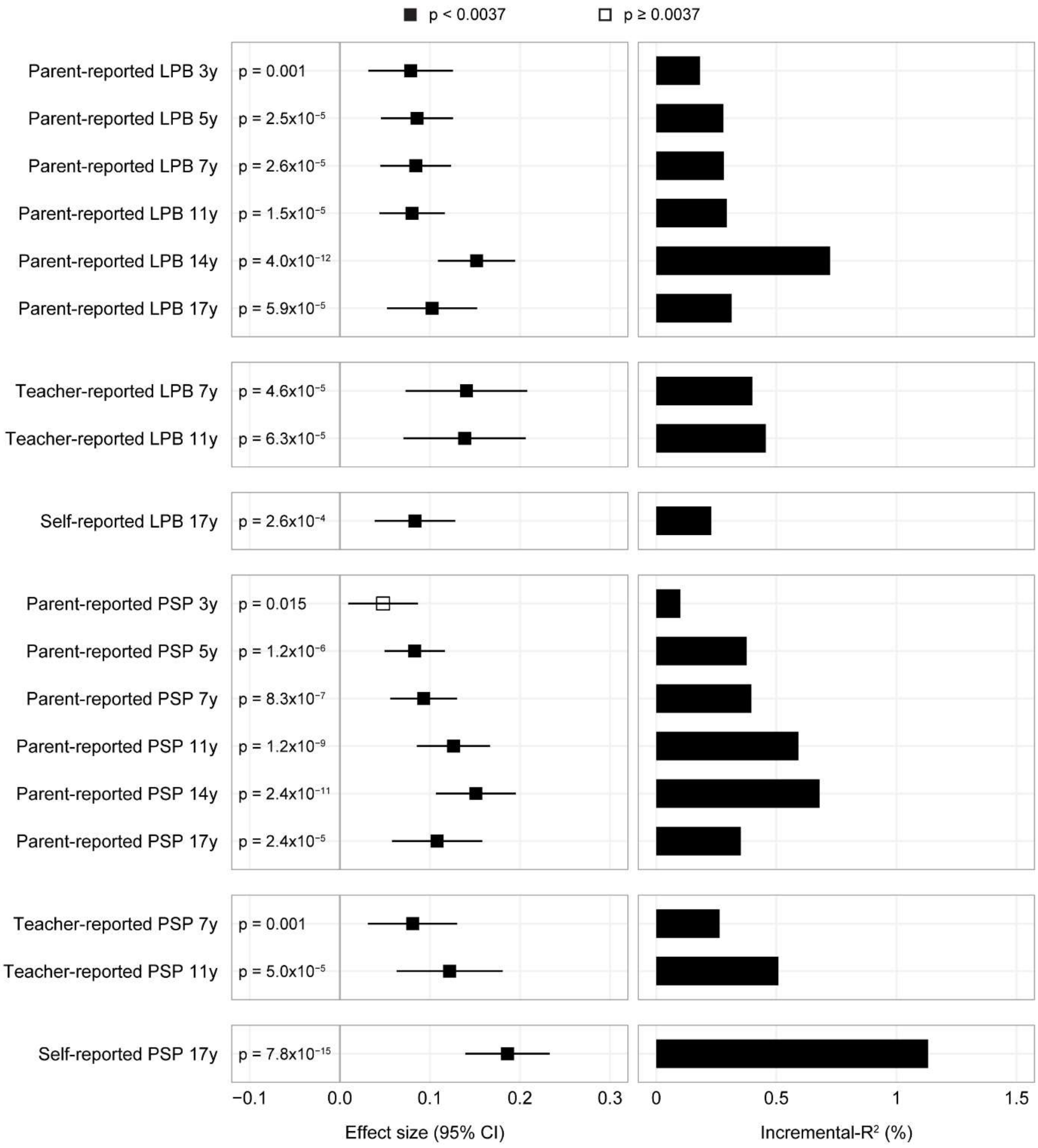

**Supplementary Figure 19.** Validation analysis of fully-matched social behaviour polygenic scores in the Millennium Cohort Study (MCS)

Polygenic scores were computed by matching discovery GWAS and target phenotypes (MCS cohort) according to social domain, reporter and age. Effect sizes are shown with their corresponding 95% confidence intervals, and the amount of variance explained is expressed in terms of incremental-$R^2$ (%). Associations passing the multiple-testing threshold ($p<0.0037$) are shown with a filled rectangle, and with an empty rectangle otherwise.

Abbreviations: LPB (Low prosocial behaviour), PSP (Peer and social problems)

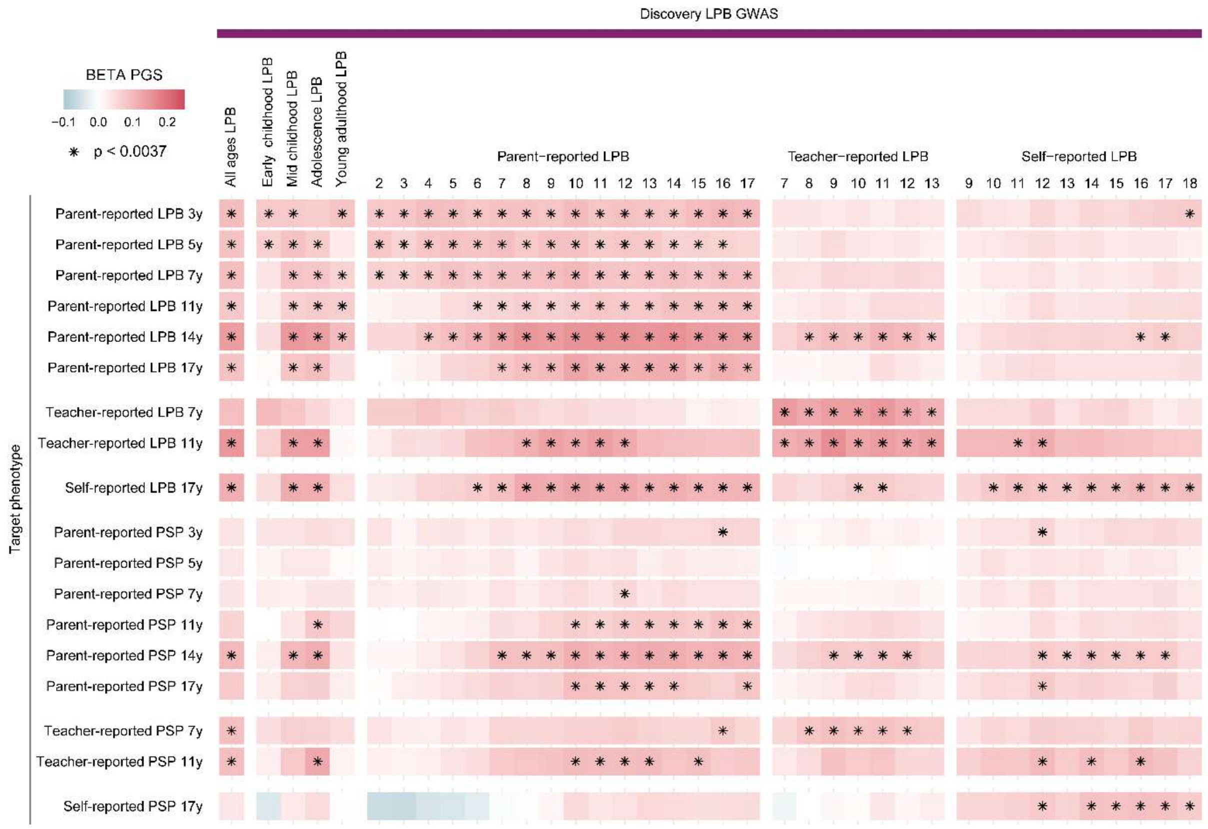

**Supplementary Figure 20.** Polygenic score analysis across all LPB discovery GWAS predicting social measures in the Millennium Cohort Study.

The discovery GWAS are shown on the x-axis and the MCS phenotypes on the y-axis, including low-resolution, medium-resolution and high-resolution social behaviour GWAS. Analyses are shown for LPB GWAS across all ages and reporters, across each developmental stage and across reporter and age. Associations passing the multiple-testing threshold (p<0.0037) are marked with an asterisk. Details for each PGS association analysis are reported in Supplementary Table 17.

Abbreviations: LPB (Low prosocial behaviour), PGS (Polygenic score)

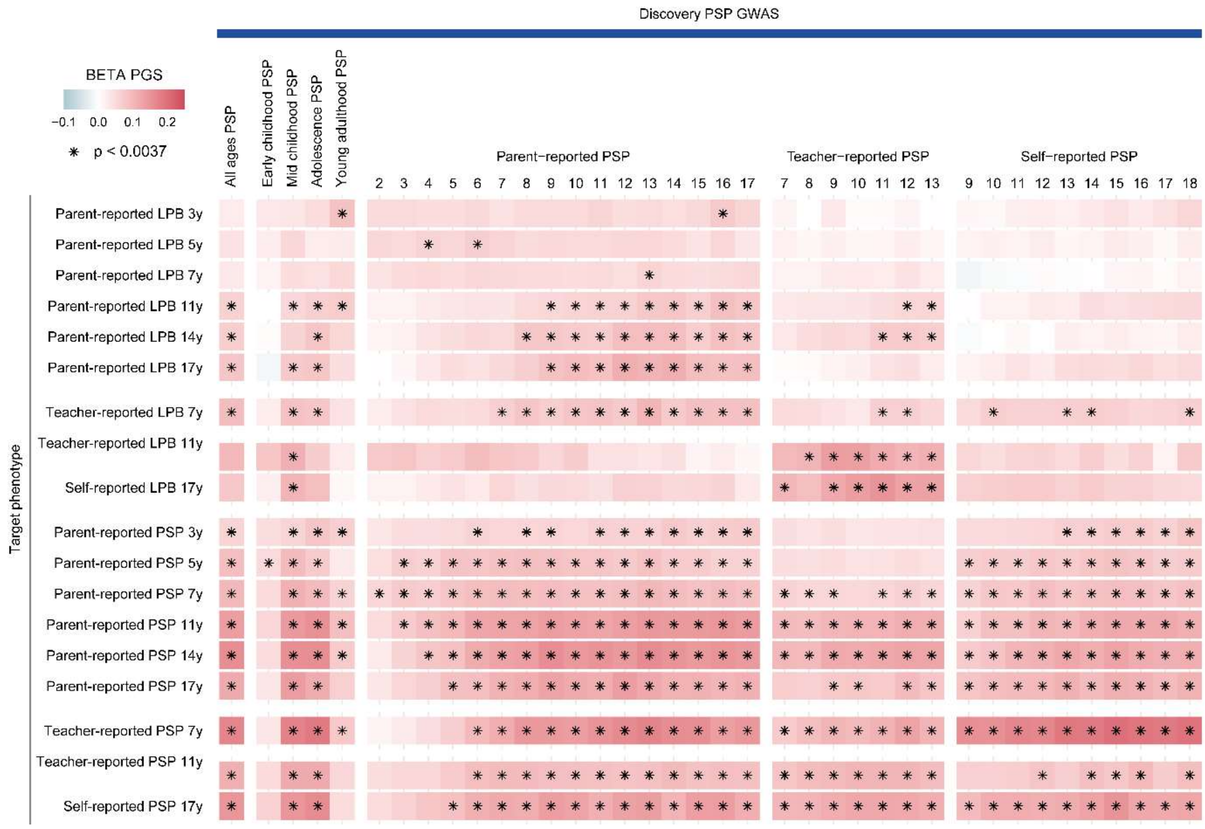

**Supplementary Figure 21**. Polygenic score analysis across all PSP discovery GWAS predicting social measures in the Millennium Cohort Study.

The discovery GWAS are shown on the x-axis and the MCS phenotypes on the y-axis, including low-resolution, medium-resolution and high-resolution social behaviour GWAS. Analyses are shown for PSP GWAS across all ages and reporters, across each developmental stage and across reporter and age. Associations passing the multiple-testing threshold ($p<0.0037$) are marked with an asterisk. Details for each PGS association analysis are reported in Supplementary Table 17.

Abbreviations: PSP (Peer and social problems), PGS (Polygenic score)

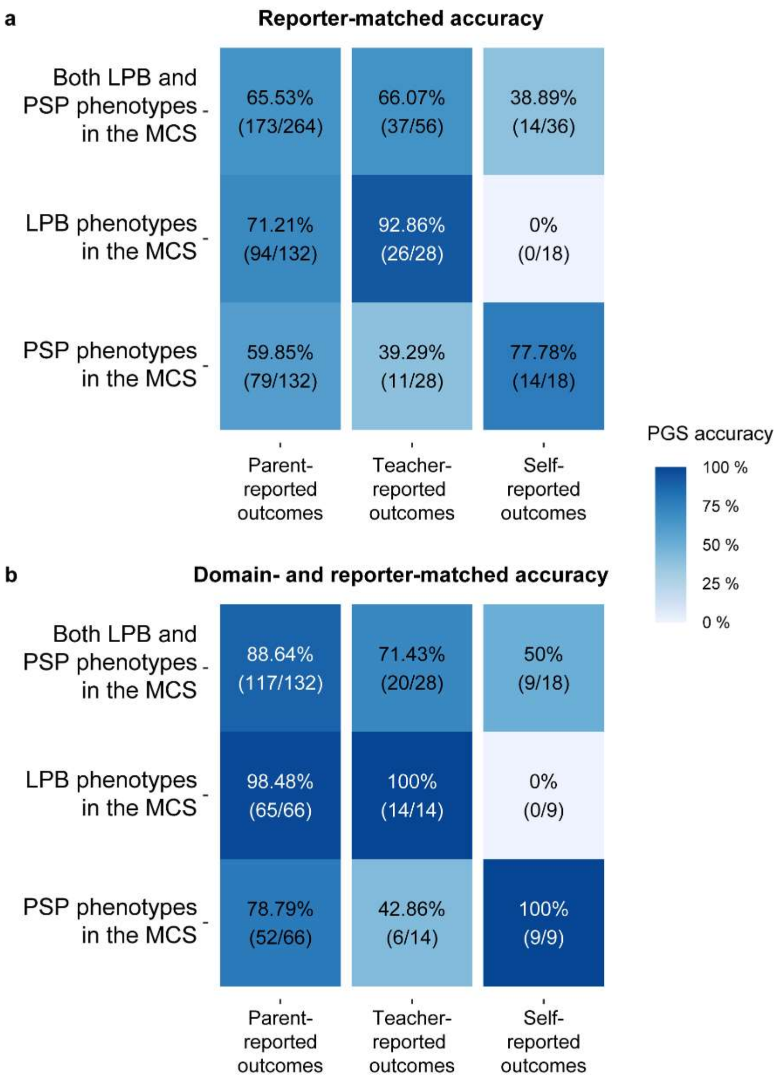

**Supplementary Figure 22.** Reporter-accuracy of PGS predictions in the Millennium Cohort Study

For each reporter, the (**a**) reporter-matched accuracy and (**b**) the combined social domain and reporter-matched accuracy are shown. The predictive accuracy of social behaviour PGS was assessed using 66 PGS generated from high-resolution social behaviour GWAS and 18 MCS social behavioural outcomes. For reporter-matched PGS compared to non-matched PGS, the signal was considered accurate if the incremental $R^2$ was higher for the matched PGS, for signals passing the multiple testing threshold. Full table comparing each measure across reporters is available at Supplementary Table 19.

Abbreviations: LPB (Low prosocial behaviour), MCS (Millennium Cohort Study), PGS (Polygenic score), PSP (Peer and social problems)

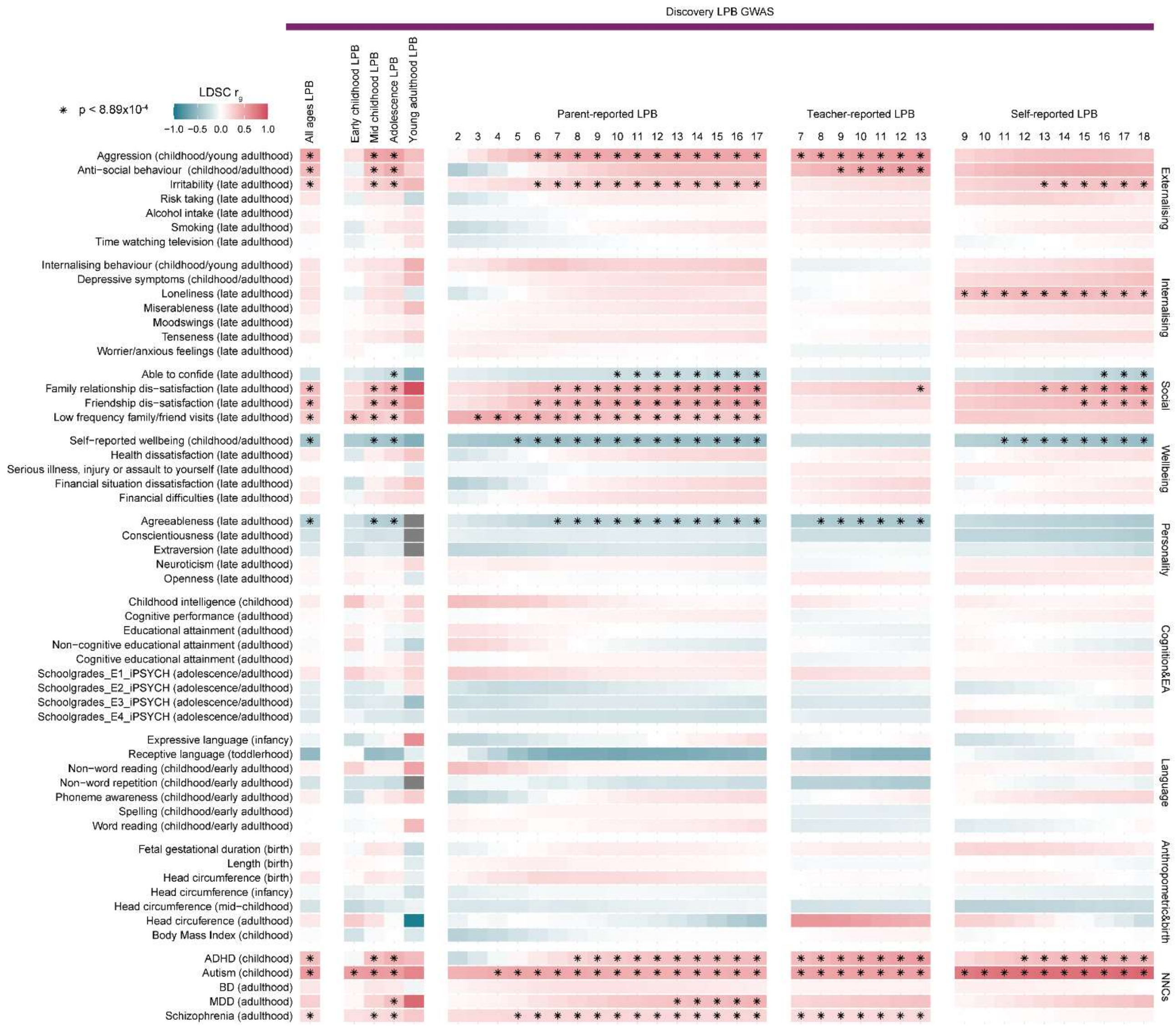

**Supplementary Figure 23.** Genetic correlations of LPB with population-based traits and NNCs

Linkage disequilibrium score (LDSC) genetic correlations with low-resolution, medium-resolution and high-resolution LPB GWAS are shown as a heatmap. Genetic correlations passing the multiple-testing threshold ($p<8.93x10^{-4}$) are marked with an asterisk. Details are reported in Supplementary Table 25.

Abbreviations: LPB (Low prosocial behaviour), NNCs (Neurodevelopmental and neuropsychiatric conditions)

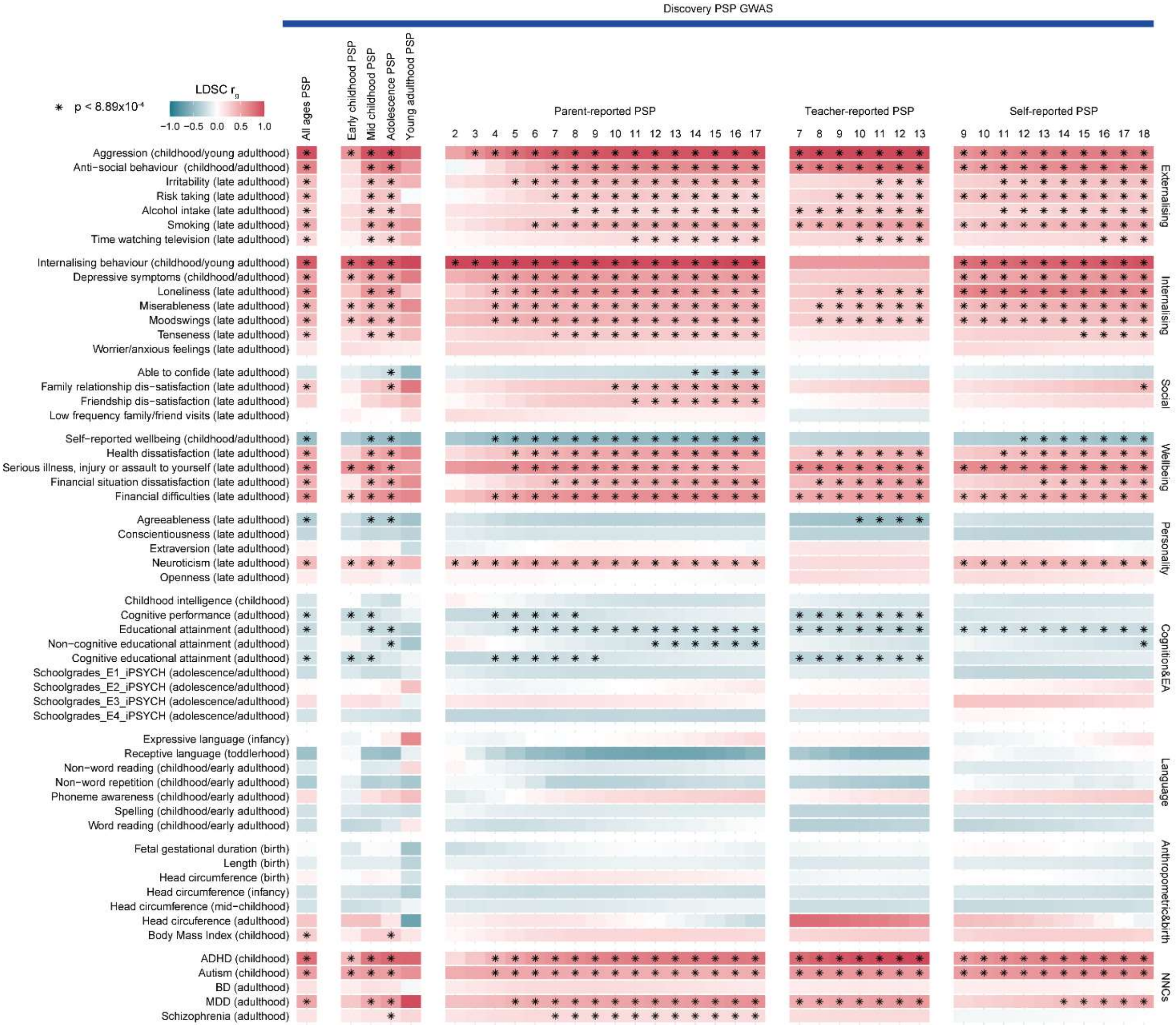

**Supplementary Figure 24.** Genetic correlations of PSP with population-based traits and NNCs

Linkage disequilibrium score (LDSC) genetic correlations with low-resolution, medium-resolution and high-resolution PSP GWAS are shown as a heatmap. Genetic correlations passing the multiple-testing threshold ($p<8.93x10^{-4}$) are marked with an asterisk. Details are reported in Supplementary Table 25.

Abbreviations: PSP (Peer and social problems), NNCs (Neurodevelopmental and neuropsychiatric conditions)

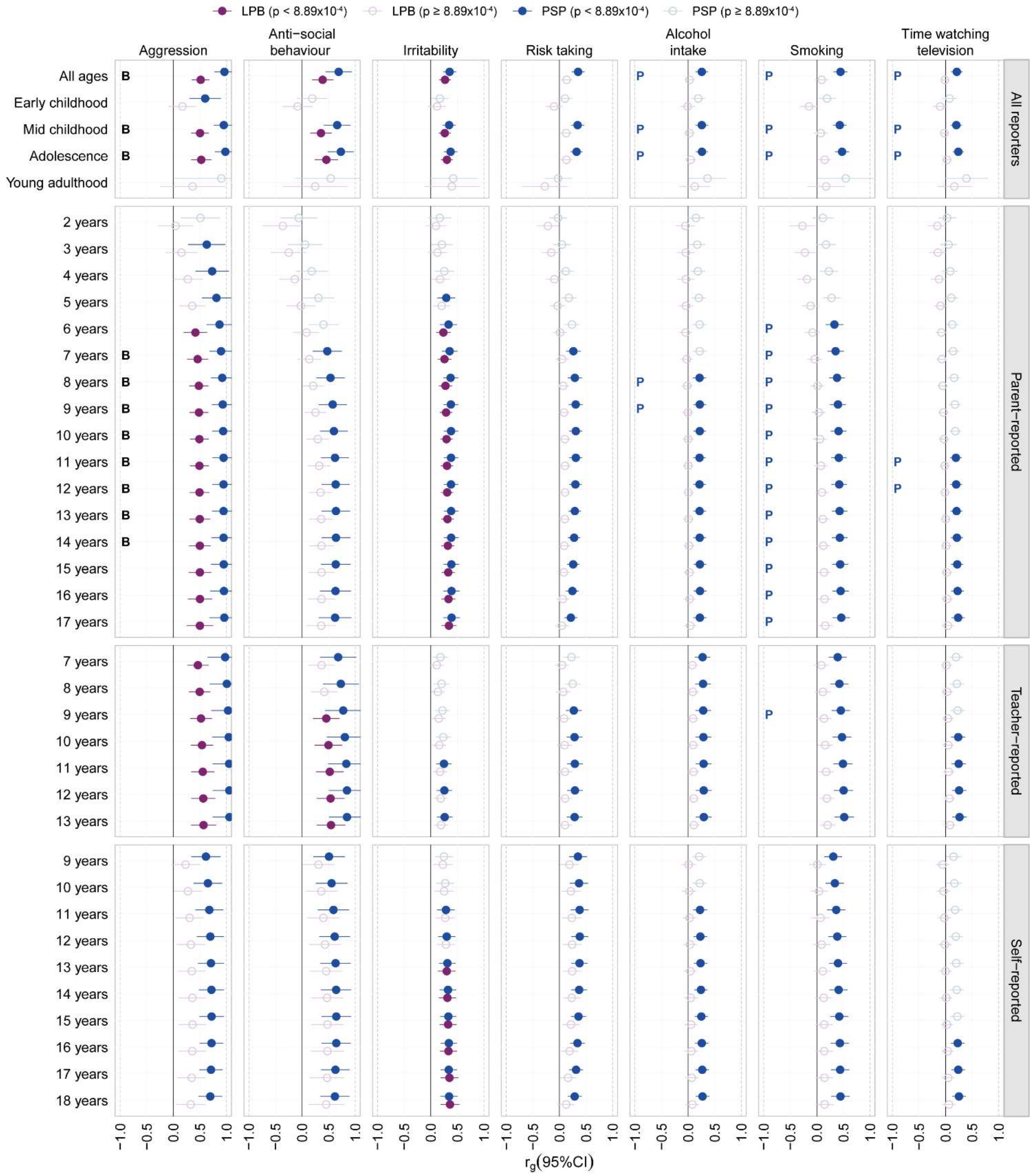

**Supplementary Figure 25.** Contextual genetic correlations of social behaviour with externalising behaviour phenotypes

Linkage disequilibrium score (LDSC) genetic correlations with low-resolution, medium-resolution and high-resolution LPB and PSP GWAS are shown. Filled circles indicate linkage disequilibrium score (LDSC) correlations passing the multiple-testing threshold ($p<8.89x10^{-4}$). Differences in PSP and LPB genetic correlations at different moderator levels, indicating specific effects based on non-overlapping 95% confidence bands, are shown for PSP correlations (blue P), LPB correlations (purple L) or both (black B), once they passed the multiple-testing threshold.

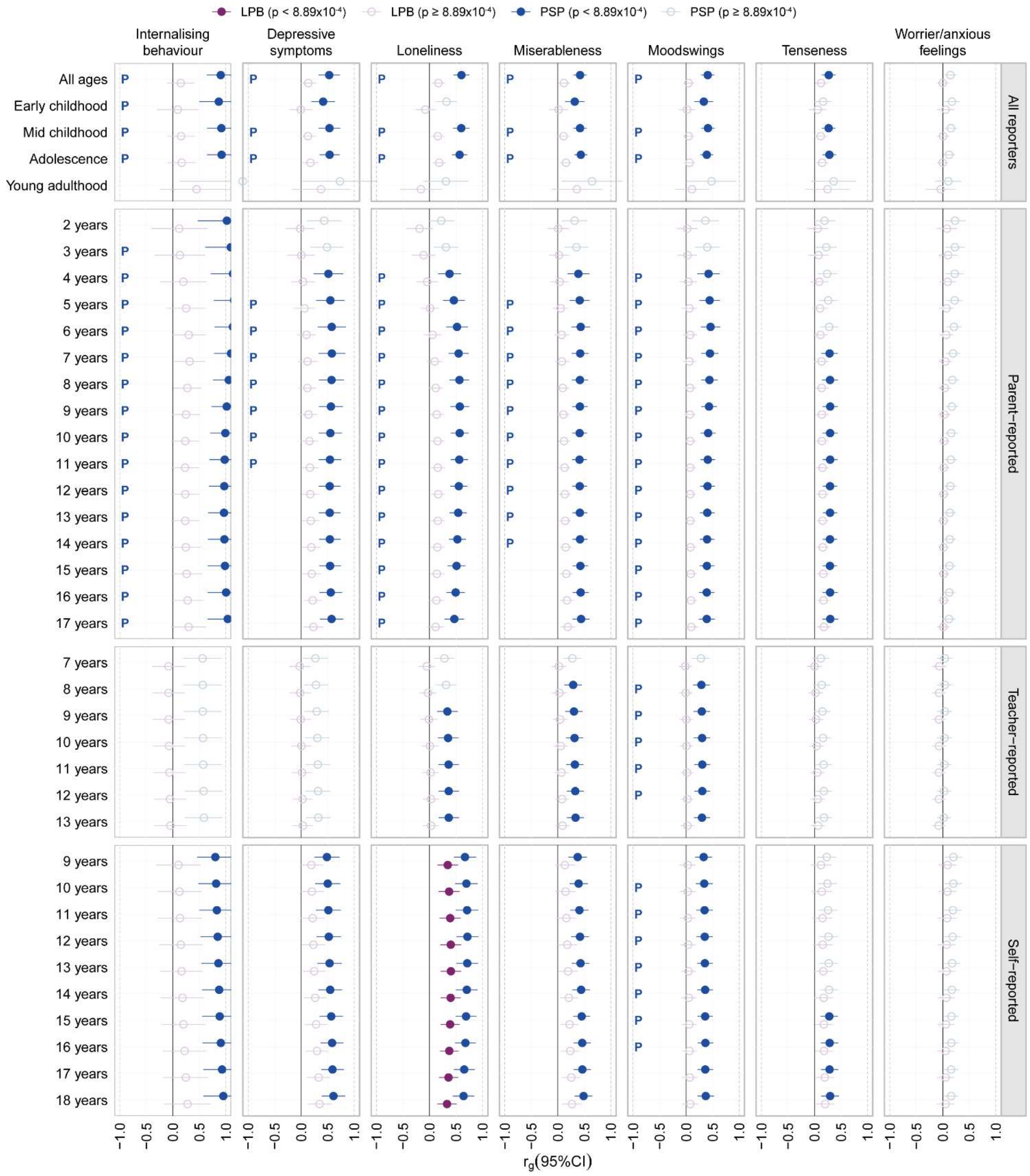

**Supplementary Figure 26.** Contextual genetic correlations of social behaviour with internalising behaviour phenotypes

Linkage disequilibrium score (LDSC) genetic correlations with low-resolution, medium-resolution and high-resolution LPB and PSP GWAS are shown. Filled circles indicate correlations passing the multiple-testing threshold ($p<8.89x10^{-4}$). Differences in PSP and LPB genetic correlations at different moderator levels, indicating specific effects based on non-overlapping 95% confidence bands, are shown for PSP correlations (blue P), LPB correlations (purple L) or both (black B), once they passed the multiple-testing threshold.

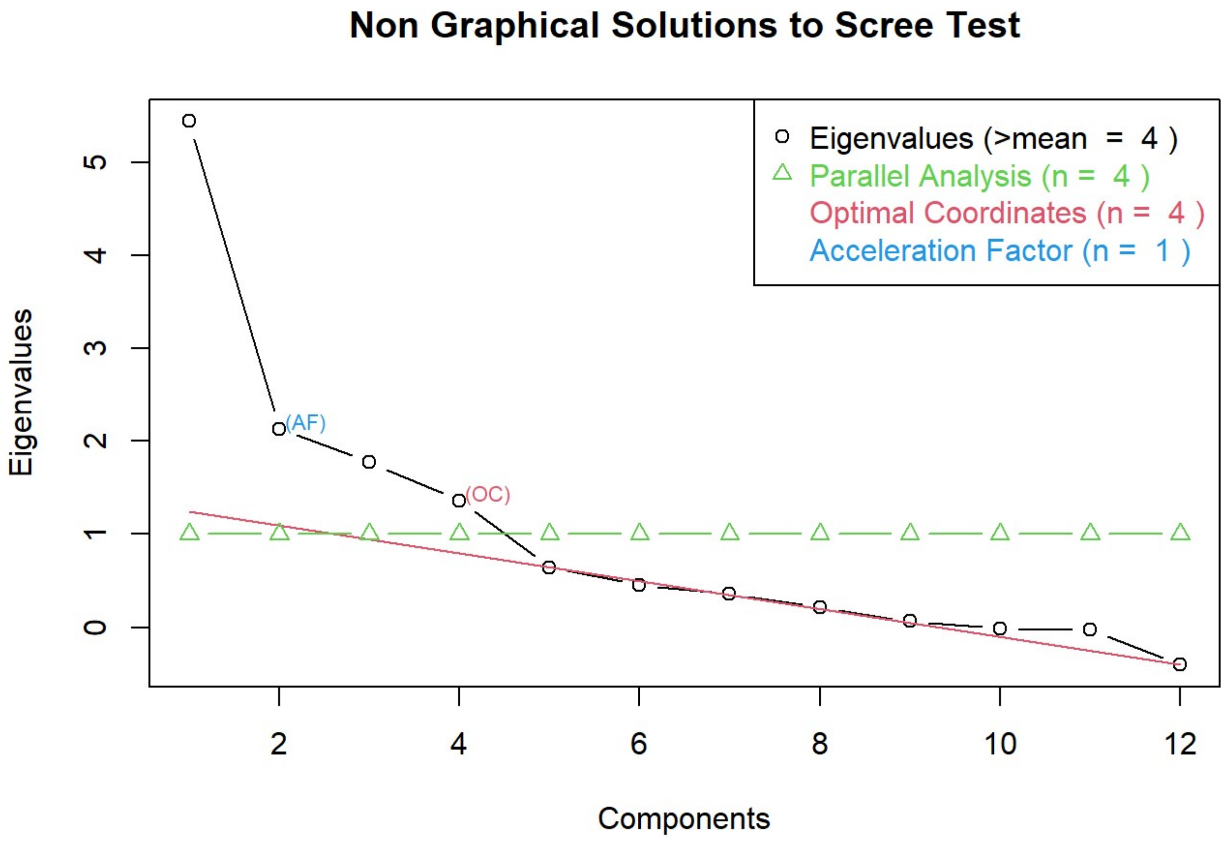

**Supplementary Figure 27.** Eigenvalue decomposition of the LDSC genetic correlation matrix between five NNCs and medium-resolution social behaviour GWAS across developmental stages.

Analyses were restricted to social behaviour GWAS statistics with sufficient power ($h^2_{SNP}$ Z-score >3).

Abbreviation: LDSC (Linkage disequilibrium score correlation), NNCs (Neurodevelopmental and neuropsychiatric conditions).

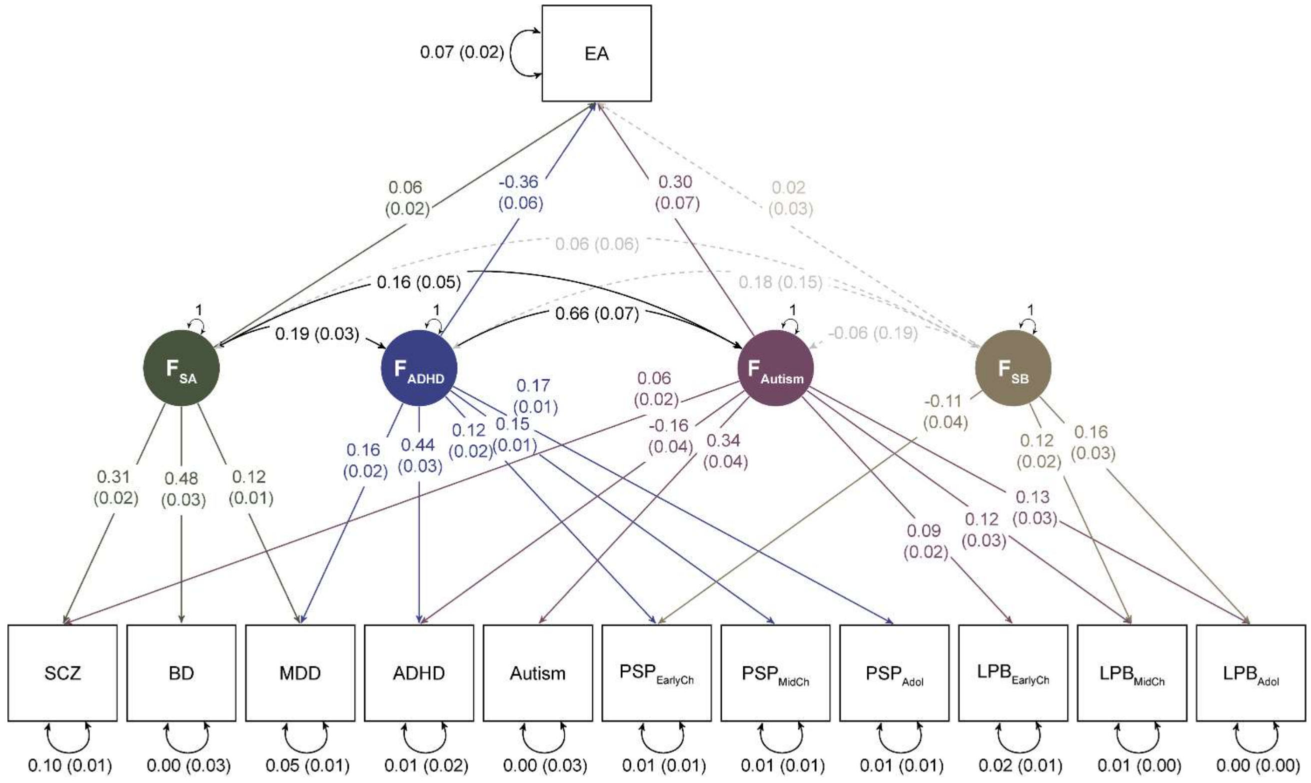

**Supplementary Figure 28**. Genetic CFA across medium-resolution social behaviour and NNCs with EA mapped.

EA was mapped onto the CFA factor structure (AIC=523.81, CFI=0.97, SRMR=0.069). Observed measures are represented by squares and latent variables by circles. Coloured, single-headed arrows define unstandardised factor loadings (shown with their corresponding SEs). Factor loadings with p ≤ 0.05 are solid, and dashed otherwise. Double-headed arrows define factor correlations (shown with their corresponding SEs). Factor correlations are represented with a black solid line if p ≤ 0.05 and with a dashed grey line otherwise. Note that unstandardised factor loadings reflect relationships at the level of the captured $h^2_{SNP}$. Analyses were restricted to social behaviour GWAS statistics with sufficient power ($h^2_{SNP}$ Z-score >3).

Model fit indices are shown in Supplementary Table 32, and standardised and unstandardised parameter estimates are shown in Supplementary Table 34.

Abbreviations: ADHD (Attention-Deficit/Hyperactivity Disorder), BD (bipolar disorder), EA (Educational attainment), $F_{ADHD}$ (ADHD genetic factor), $F_{Autism}$ (autism genetic factor), $F_{SA}$ (schizoaffective genetic factor), $F_{SB}$ (social behaviour genetic factor), LPB (Low prosocial behaviour), $LPB_{EarlyCh}$ (Early childhood LPB), $LPB_{MidCh}$ (Mid childhood LPB), $LPB_{Adol}$ (Adolescence LPB), MDD (major depressive disorder), NNCs (Neurodevelopmental and neuropsychiatric conditions), PSP (Peer and social problems), $PSP_{EarlyCh}$ (Early childhood PSP), $PSP_{MidCh}$ (Mid childhood PSP), $PSP_{Adol}$ (Adolescence PSP), SCZ (schizophrenia)

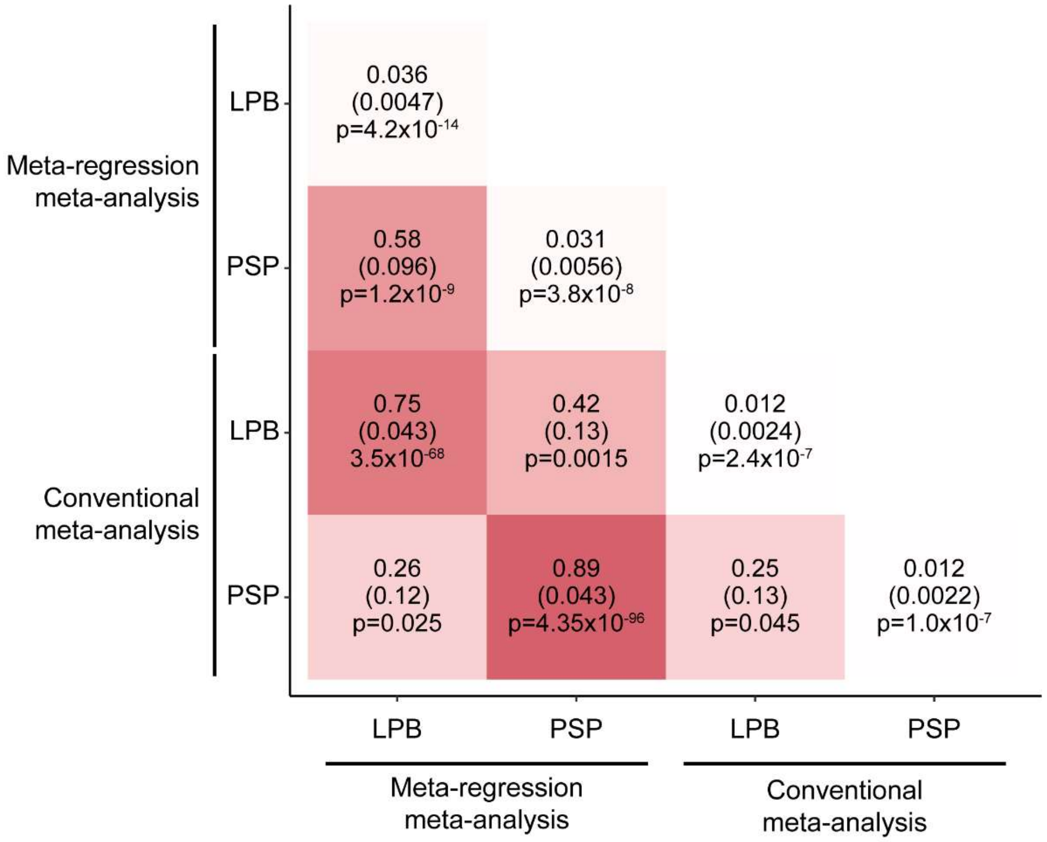

**Supplementary Figure 29.** Sensitivity analysis. LDSC heritability and genetic correlations for low-resolution social behaviour GWAS across meta-regression GWAS and conventional GWAS methods

Diagonal values indicate LDSC $h^2_{SNP}$ and off-diagonal values LDSC $r_g$ as estimated with linkage disequilibrium score (LDSC) regression and correlation, respectively. As part of a sensitivity analysis, we compared the genetic architecture between low-resolution LPB and PSP GWAS as estimated with a random-effects meta-regression GWAS and a conventional meta-GWAS (multivariate N-GWAMA). Details are provided in Supplementary Table 35.

Abbreviations: LPB (Low prosocial behaviour), PSP (Peer and social problems)

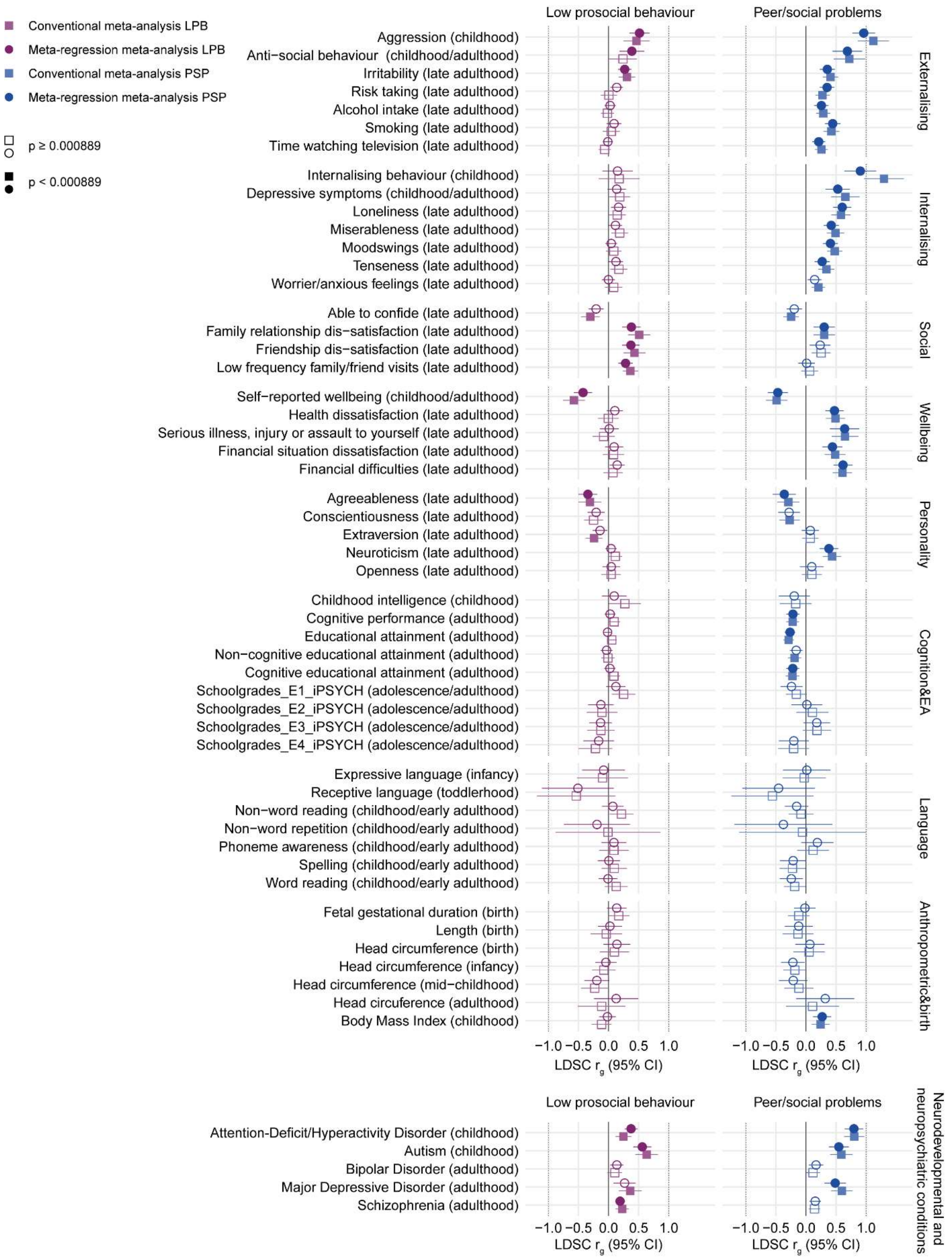

**Supplementary Figure 30.** Comparison of genetic correlations of low-resolution social behaviour with population-based traits and NNCs across meta-regression GWAS and conventional GWAS methods.

Linkage disequilibrium score (LDSC) genetic correlations with $LPB_{All}$ and $PSP_{All}$ GWAS are shown. Details are provided in Supplementary Table 36.

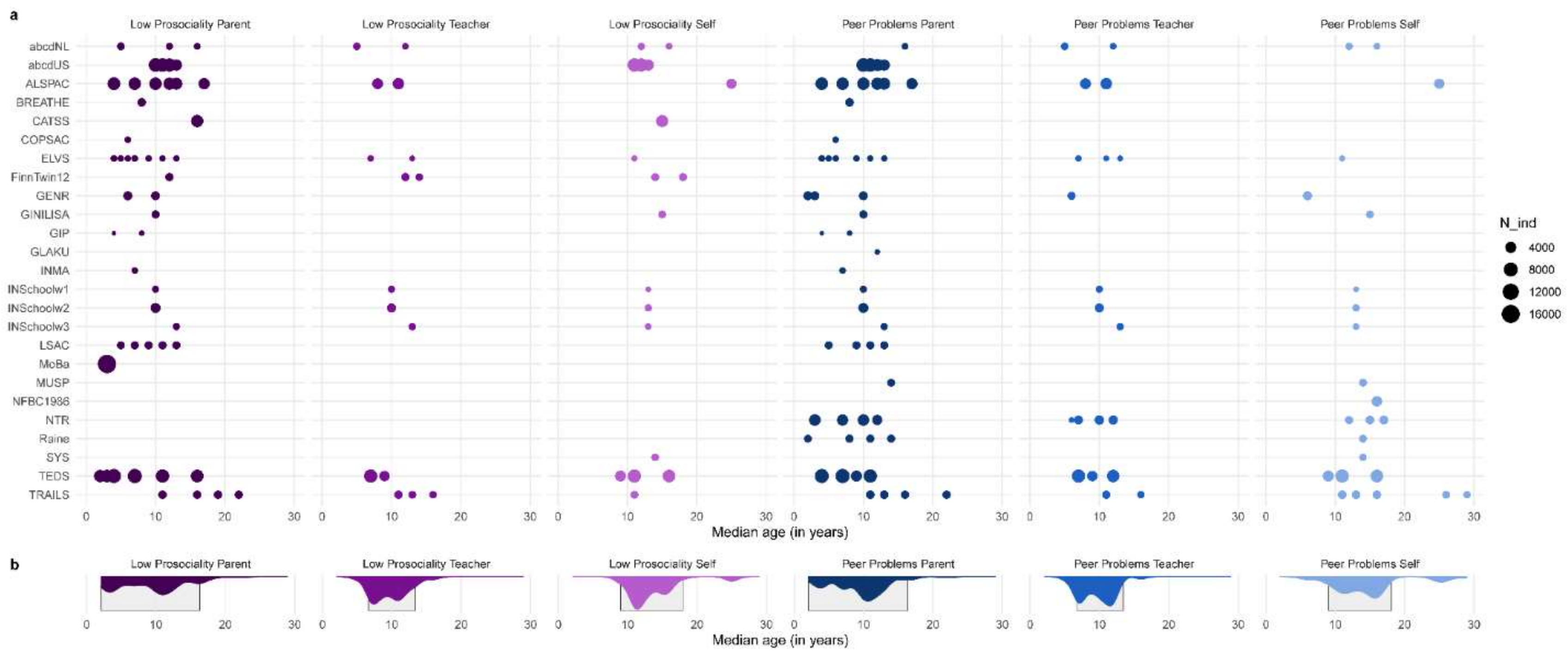

**Supplementary Figure 31.** Data availability and prediction ranges

(**a**) Overview of available data across the 25 cohorts per social domain (LPB or PSP) and reporter (parent, teacher or self) spanning ages 2 to 29 years. Each dot represents a GWAS summary statistics (total $N_{GWAS}$=195), and its size represents the number of individuals included in that GWAS. (**b**) Distributions across all cohorts are shown by age for each social domain and reporter. Additionally, prediction ranges for meta-regression GWAS are shown as squares. We derived parent-reported LPB and PSP GWAS from 2-17 years, teacher-reported LPB and PSP GWAS from 7-13 years and self-reported LPB and PSP GWAS from 9-18 years.

Abbreviations: GWAS (genome-wide association study), LPB (Low prosocial behaviour), PSP (Peer and social problems)

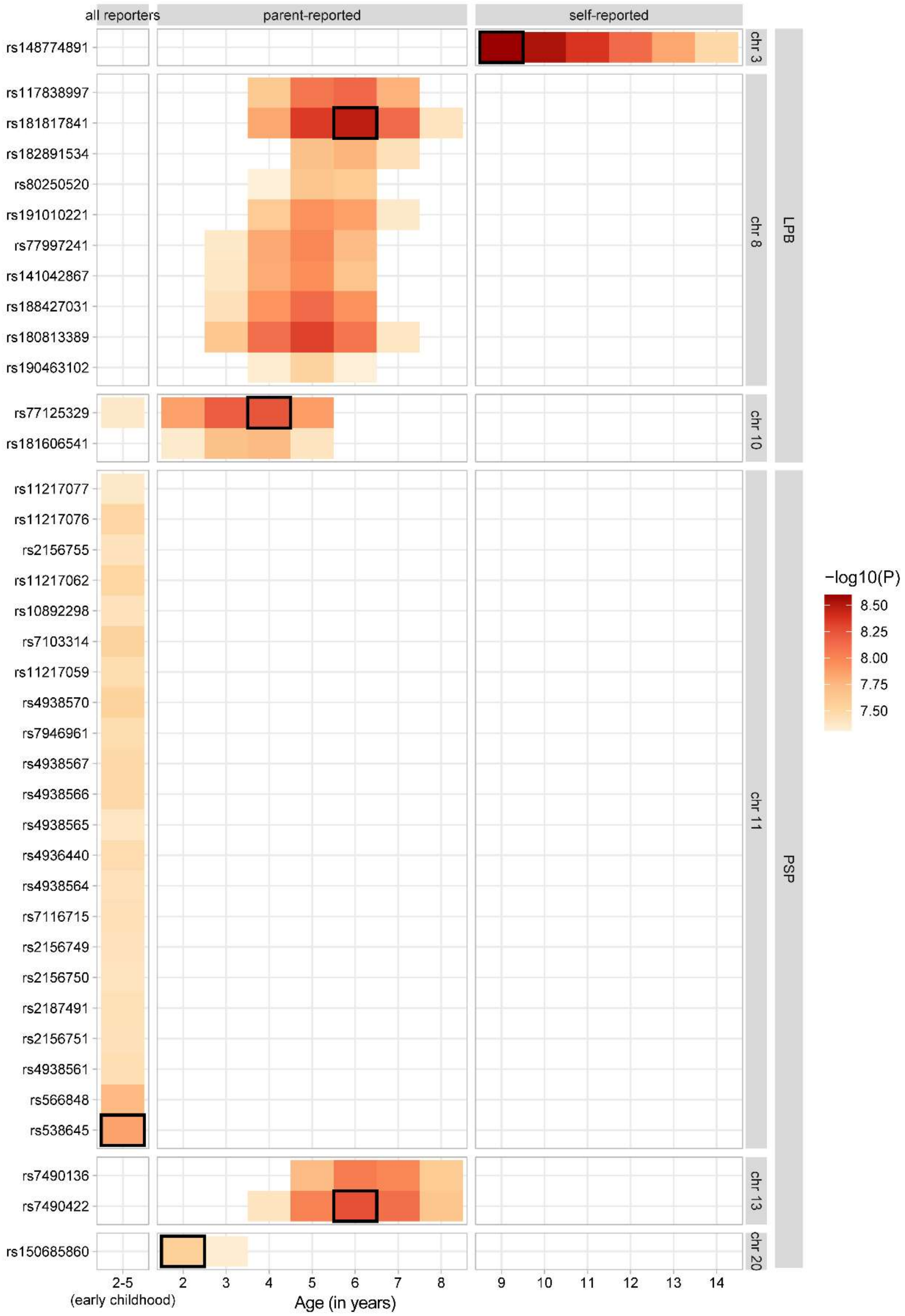

**Supplementary Figure 32.** Social behaviour GWAS selection for FUMA and MAGMA analysis

SNPs passing the genome-wide significance threshold are shown on the y-axis. Each square represents a GWAS summary statistic with a genome-wide signal with a -log10 p-value as indicated by the colour code. Black boxes indicate the summary statistics with the strongest signal at each genome-wide locus, as selected for FUMA and MAGMA analysis.

| | | Cohort-level data | | Seed-based meta-analysis (METAL) | Developmental window-based meta-analysis (GWAMA) | Social domain-based meta-analysis (GWAMA) |
|---|---|---|---|---|---|---|
| LPB | Early Childhood (0-5y) | ALSPAC, ELVS, GIP, LSAC, MoBa, TEDS | $N_{ind}$=33,062 $N_{obs}$=50,171 | 3 seeds $N_{ind}$=13,960 – 22,239 | $N_{SNPs}$= 8,848,838 $N_{effective}$=35,433 | $N_{SNPs}$= 13,982,147 $N_{effective}$=154,763 |
| | Mid Childhood (6-11y) | ABCD-NL, ABCD-US, ALSPAC, BREATHE, COPSAC, ELVS, GENR, GINILISA, INMA_GSA, INMA_Omni1, INSchoolw1, INSchoolw2, LSAC, TEDS, TRAILS | $N_{ind}$=36,214 $N_{obs}$=87,394 | 5 seeds $N_{ind}$=17,307 – 17,579 | $N_{SNPs}$= 13,409,195 $N_{effective}$=59,902 | |
| | Adolescence (12-17y) | ABCD-NL, ABCD-US, ALSPAC, CATSS, ELVS, FinnTwin12, GINILISA, INSchoolw1, INSchoolw2, INSchoolw3, LSAC, SYS, TEDS, TRAILS | $N_{ind}$=34,354 $N_{obs}$=100,358 | 6 seeds $N_{ind}$=16,535 – 16,823 | $N_{SNPs}$= 13,317,789 $N_{effective}$=57,488 | |
| | Young adulthood (18-30y) | ALSPAC, FinnTwin12, TRAILS | $N_{ind}$=5,819 $N_{obs}$=7,263 | 2 seeds $N_{ind}$=2,782 – 4,481 | $N_{SNPs}$= 9,531,777 $N_{effective}$=6,231 | |
| PSP | Early Childhood (0-5y) | ALSPAC, ELVS, GENR, INMA_GSA, LSAC, NTR, The Raine Study, TEDS | $N_{ind}$=23,709 $N_{obs}$=30,699 | 3 seeds $N_{ind}$=8,613 – 12,397 | $N_{SNPs}$= 9,245,774 $N_{effective}$=22,412 | $N_{SNPs}$= 14,859,523 $N_{effective}$=161,163 |
| | Mid Childhood (6-11y) | ABCD-NL, ABCD-US, ALSPAC, BREATHE, COPSAC, ELVS, GENR, GINILISA, INMA_GSA, INMA_Omni1, INSchoolw1, INSchoolw2, LSAC, NTR, The Raine Study, TEDS, TRAILS | $N_{ind}$=41,948 $N_{obs}$=108,783 | 7 seeds $N_{ind}$=15,289 – 15,820 | $N_{SNPs}$= 13,458,404 $N_{effective}$=75,189 | |
| | Adolescence (12-17y) | ABCD-NL, ABCD-US, ALSPAC, ELVS, GINILISA, GLAKU, INSchoolw1, INSchoolw2, INSchoolw3, LSAC, MUSP, NFBC1986, NTR, The Raine Study, SYS, TEDS, TRAILS | $N_{ind}$=35,374 $N_{obs}$=100,059 | 7 seeds $N_{ind}$=14,130 – 14,550 | $N_{SNPs}$= 14,097,988 $N_{effective}$=51,130 | |
| | Young adulthood (18-30y) | ALSPAC, TRAILS | $N_{ind}$=4,478 $N_{obs}$=6,519 | 3 seeds $N_{ind}$=1,201 – 3,877 | $N_{SNPs}$= 9,016,994 $N_{effective}$=5,002 | |

**Supplementary Figure 33.** Pipeline for a conventional GWAS meta-analysis

The description of a 3-step pipeline including a combination of METAL and multivariate N-GWAMA software to conduct a conventional GWAS meta-analysis is provided in Supplementary Note 11. Each GWAS seed included a partial meta-analysis across a subset of GWAS statistics based on independent individuals. Multivariate N-GWAMA analyses combined GWAS seeds, accounting for sample overlap. The effective sample size was estimated with multivariate N-GWAMA software.
