## Supplementary Notes for "Genome-wide analysis of social behaviour in context: a meta-regression approach across social domains, reporters and developmental stages"

### SUPPLEMENTARY NOTE 1. COHORT DESCRIPTIONS AND ETHICAL APPROVAL

#### ABCD-NL

The ABCD study (Amsterdam Born Children and their Development) is a large prospective community-based birth cohort in the Netherlands[1]. Regarding the DNA collection and analysis, an opt-out procedure was used (METC approval 2002_039#B2013531). The inclusion criteria for the genotyping were as follows: unrelated children who were of Dutch ancestry, availability of a blood sample, good quality of extracted DNA for genotyping, and the willingness to participate in genetic investigations. This resulted in 1,192 participants. Participants were excluded based on the quality: genetic quality control (due to call rate <95%; heterozygosity, phenotype-genotype gender mismatch, and relatedness; n=45), resulting in 1147 remaining participants.

All ABCD participants gave written informed consent for data collection of the phenotypes. The ABCD study protocol was approved by the Central Committee on Research Involving Human Subjects in the Netherlands, the medical ethics review committees of the participating hospitals, and the Registration Committee of the Municipality of Amsterdam.

#### ABCD-US

The ABCD study is a multisite, population-based, longitudinal study of brain development and child health in the United States designed to recruit more than 10,000 children aged 9-10 years and follow them over 10 years into early adulthood[2]. The current study is based on ABCD Data Release 4.0. Starting at the ages of 9-10 years, the ABCD study follows participants for 10 years at 21 data acquisition sites across the United States. Participants (N=11,875) were recruited through school systems sampling across gender, race and ethnicity, socioeconomic status, and urbanicity[3].

Parents or guardians provided written consent, while children provided written assent[4]. The Institutional Review Board (IRB) at the University of California, San Diego, approved all aspects of the ABCD study[5].

#### ALSPAC

Pregnant women resident in Avon, UK, with expected dates of delivery between 1st April 1991 and 31st December 1992 were invited to take part in the study[6,7]. The initial number of pregnancies enrolled was 14,541 and, of these, 13,988 children were alive at 1 year of age. When the oldest children were approximately 7 years of age, an attempt was made to bolster the initial sample with eligible cases who had failed to join the study originally. As a result, when considering variables collected from the age of seven onwards (and potentially abstracted from obstetric notes) there are data available for more than the 14,541 pregnancies mentioned above: The number of new pregnancies not in the initial sample (known as Phase I enrolment) that are currently represented in the released data and reflecting enrolment status at the age of 24 is 906, resulting in an additional 913 children being enrolled (456, 262 and 195 recruited during Phases II, III and IV respectively). The phases of enrolment are described in more detail in the cohort profile paper and its update. The total sample size for analyses using any data collected after the age of seven is therefore 15,447 pregnancies, resulting in 15,658 foetuses. Please note that the study website contains details of all the data that are available through a fully searchable data dictionary and variable search tool (http://www.bristol.ac.uk/alspac/researchers/our-data/).

Ethical approval for the study was obtained from the ALSPAC Ethics and Law Committee and the Local Research Ethics Committees. Consent for biological samples has been collected following the Human Tissue Act (2004). Informed consent for the use of data collected via questionnaires and clinics was obtained from participants following recommendations of the ALSPAC Ethics and Law Committee at the time.

#### BREATHE

The BREATHE project (European Commission: FP7-ERC-2010-AdG, ID 268479) is a population-based cohort of primary school children designed to analyse the association between air pollution and behaviour, cognitive function and brain morphology[8] (https://www.isglobal.org/ca/-/breathe-brain-development-and-air-pollution-ultrafine-particles-in-school-children). Thirty-six of the 416 schools in Barcelona were selected based on modelled levels of traffic-related nitrogen dioxide. Thirty-eight schools were located in Barcelona, and one school was in an adjacent municipality, Sant Cugat del Vallés. All families of children without special needs who were enrolled in 2nd, 3rd, and 4th grades at the selected schools were invited to participate in the study (2012). A total of 2897 children aged 7 to 11 years accepted the invitation and participated in the project. Genotype data were available for 1667 children of European genetic ancestry.

All parents or legal guardians provided written informed consent. The study was approved by the IMIM-Parc de Salut Mar Research Ethics Committee (No. 2010/41221/I), Barcelona, Spain, and the FP7-ERC-2010-AdG Ethics Review Committee (268479-22022011).

#### CATSS

The Child and Adolescent Twin Study in Sweden (CATSS) is an ongoing longitudinal twin study targeting all twins born in Sweden since 1992[9,10]. Parents of twins were interviewed regarding the twins' somatic and mental health in connection with their 9$^{th}$ and 12$^{th}$ birthdays, and both parents and twins completed questionnaires in connection with their 15$^{th}$ and 18$^{th}$ birthdays. DNA samples (from saliva) were obtained from the CATSS participants at study enrolment. The twins in CATSS are predominantly of European ancestry.

Informed consent was provided by participants in CATSS. The study and analyses of genetic data were approved by the Karolinska Institute Ethical Review Board and the Swedish Ethical Review Authority.

#### COPSAC

The Copenhagen Prospective Studies on Asthma in Childhood 2010 (COPSAC2010) is an ongoing longitudinal population-based clinical mother-child cohort[11] which includes a comprehensive cognitive and psychopathological assessment of the child at age 10 years (The Copenhagen Prospective Study on Neuro-PSYCHiatric Development [COPSYCH])[12]. The COPSAC2010 cohort[11] is an ongoing mother-child population-based longitudinal clinical cohort (N=700, 48.57% females, 95.7% of European ancestry, mean [SD] of maternal age at birth 32.28 [4.36]) recruited at pregnancy week 24 in the period between 2008 and 2010. The mothers participated in a randomised control trial of a high dose of vitamin D and fish oil supplementation in the third trimester of pregnancy as part of a longitudinal study to investigate the long-term effects of these supplements in the offspring, with asthma as the main outcome[11].

The participants took part in the study voluntarily and after written informed consent by the caretaker. The participating families are informed that the study is unlikely to benefit their child and that discoveries will only benefit future generations. The main study protocol for starting the cohort was approved by the Ethics Committee (H-B-2008-093) and the Danish Data Protection Agency (2008-41-2599).

#### ELVS

The Early Language in Victoria Study (ELVS) is a longitudinal, population-based cohort study from Melbourne, Victoria, Australia[13], which follows children from infancy (8 months) to adolescence (13 years). Parents of 1,910 infants living in six local government areas of varying socioeconomic status in Melbourne, Victoria, Australia, were recruited between September 2003 and April 2004. Data were collected for a wide range of speech, language, literacy, health and psychosocial measures.

Written informed consent was provided by the parents, and ethical approval was obtained from the Royal Children's Hospital (23018 and 27078) and La Trobe University Human Ethics Committee (03-32).

#### FinnTwin12

The target population was all twins in Finland born between 1983 and 1987, all are of European ancestry. They and their parents were asked to participate in the study when the twins were aged 11-12 years, and they have been asked to take part in subsequent waves of data collection at ages 14 and 17 years, and as young adults. For details see [14].

The participants provided written informed consent prior to blood or saliva sampling for DNA as young adults (wave 4 data collection). The ethics committee of the Department of Public Health of the University of Helsinki (Helsinki, Finland), the ethics committee of the Helsinki University Central Hospital District (Helsinki, Finland, HUS/2226/2021), and the Institutional Review Board of Indiana University (Bloomington, Indiana, USA) approved the FinnTwin12 study protocol. HUS reviews the study annually, and 2023's statement is number 4/2023, dated February 1, 2023.

#### GenR

The Generation R Study is a population-based prospective cohort study[15]. All children were born between April 2002 and January 2006. This study is designed to identify early environmental and genetic predictors of growth, development, and health from foetal life until young adulthood[15]. GWAS were conducted on a maximum of 2,191 individuals of Northern European ancestry from Generation R.

Genotype data were derived from cord blood at birth or from venepuncture during a visit to the research centre, using Illumina 610K and 660K genotype arrays (Illumina, San Diego, CA, USA). SNP-level filtering included genotype call rate < 95%, minor allele frequency (MAF) < 1%, and Hardy-Weinberg Equilibrium (HWE) p < .000001. Individuals were filtered based on individual call rate < 95%, outlying heterozygosity, non-European ethnicity, missing phenotype, and relatedness (pairwise IBD >0.185). Genotype data that passed quality control were subsequently imputed using the Haplotype Reference Consortium (HRC) release 1.1 as the reference panel. An extensive description of the calling procedures and subsequent quality control has been described elsewhere[16].

Participants in GenR have given written informed consent. The study protocol has been approved by the Medical Ethical Committee of the Erasmus Medical Centre, Rotterdam.

#### GINILISA

Data were combined from two German birth cohort studies, the German Infant Study on the Influence of Nutrition Intervention plus Air pollution and Genetics on Allergy Development (GINIplus) and the Influence of Lifestyle factors on Development of the Immune System and Allergies in East and West Germany (LISA) study. For GINIplus, a total of 5,991 mothers and their newborns were recruited between September 1995 and June 1998 in Munich and Wesel. Infants with at least one allergic parent and/or sibling were allocated to the interventional study arm investigating the effect of different hydrolysed formulas for allergy prevention in the first year of life. All children without a family history of allergic diseases and children whose parents did not give consent for the intervention were allocated to the non-interventional arm[17]. For LISA, a total of 3,094 healthy, full-term neonates were recruited between 1997 and 1999 in Munich, Leipzig, Wesel and Bad Honnef. The participants were not preselected based on family history of allergic diseases[18]. DNA was collected at the ages of 6 and 10 years in both studies, and 1,511 children of European ancestry from the Munich study centre were genotyped.

For both studies, written consent from participants' families and approval by the local Ethics Committees (Bavarian Board of Physicians, Medical Faculty of the University of Leipzig, Board of Physicians of North-Rhine-Westphalia) were obtained.

#### GLAKU

The adolescents of the GLAKU (Glycyrrhizin in Licorice) cohort came from an urban community-based cohort comprising 1,049 infants born between March and November 1998 in Helsinki, Finland[19]. In 2009–2011, initial cohort members who had given permission to be contacted and whose addresses were traceable (N=920, 87.7% of the original cohort in 1998) were invited to a follow-up, of which 692 (75.2%) could be contacted by phone (mothers of the adolescents). Of them, 451 (65.2% of those who could be contacted by phone, 49% of the invited) participated in a follow-up at a mean age of 12.3 years (SD = 0.5, range 11.0–13.2 years).

Informed consent was obtained from all participants. The study protocol was approved by the ethical committees of the City of Helsinki and the Uusimaa Hospital District.

#### INMA

The INfancia y Medio Ambiente – Environment and Childhood (INMA) Project (http://www.proyectoinma.org/)[20] is a network of prospective population-based, mother-child birth cohorts in Spain that aims to study the role of environmental pollutants in air, water, and diet during pregnancy and early childhood in relation to child growth, health, and development. Data were collected between 2003 and 2008, and children have been followed up from birth to puberty. Within this study, we used data from two independent sub-cohorts: INMA_GSA and INMA_Omni1.

INMA_GSA. The INMA-Gipuzkoa cohort (INMA_GSA) forms an independent sub-cohort within INMA[20]. In the INMA_GSA sub-cohort, pregnant women were recruited between April 2006 and January 2008. Written informed consent was obtained from all participating parents. Genetic data for the INMA_GSA sub-cohorts were obtained with the Infinium Global Screening Array (GSA). The study was approved by the Institutional Boards of the recruiting centres and by the Basque Ethics Committee.

INMA_Omni1. Data for the INMA_Omni1 study come from the INMA-Sabadell and INMA-Valencia sub-cohorts of the INMA Project[20]. Pregnant women were recruited from November 2003 to June 2005 for the INMA-Valencia sub-cohort and from July 2004 to July 2006 for the INMA-Sabadell sub-cohort. Written informed consent was obtained from all participating parents. Genetic data for the INMA_ Omni1 sub-cohorts were obtained with the Illumina Human-Omni1 SNP array. The study was approved by the Ethical Committee of each participating centre.

#### INSchool

INSchool is a Spanish population-based sample of 6,500 children from primary and secondary schools in Catalonia, Spain[21]. All parents or surrogates completed the Child Behavior Checklist for ages 6-18 years (CBCL-6/18) or Youth Self-Report for ages 11-18 years (YSR-11/18) from Achenbach System of Empirically Based Assessment (ASEBA) and the Strengths and Difficulties Questionnaire (SDQ). ADHD symptoms were assessed with the Conners Rating Scale for parents and teachers (CPRS) or the ADHD Rating Scale-5 for Children and Adolescents. In the diagnostic phase, participants classified as positive according to the CBCL were clinically interviewed to confirm/discard the diagnosis of neurodevelopmental disorders using the Diagnostic Interview Schedule for Affective Disorders and Schizophrenia for School-Age Children 6-18 years–Lifetime Version DSM5 (K-SADS-PL). Genomic DNA was isolated from saliva using the Oragene DNA kit (OG-500; DNA Genotek Inc., Ottawa, ON, Canada), and genome-wide genotyping was performed with the Illumina Global Screening Array (GSA).

Written informed consent was obtained from participants' parents before inclusion into the study. The study was approved by the Clinical Research Ethics Committee of the Hospital Universitari

Vall d'Hebron. All methods were performed in accordance with the relevant guidelines and regulations.

#### LSAC

The Longitudinal Study of Australian Children study (LSAC) includes prospective birth (B) and kinder (K, not further considered in the EAGLE consortium) cohorts, which aimed to be broadly representative of the Australian population[22]. The B-cohort recruited 5,107 0-1-year olds in 2004, with continued in-home follow-up every 2 years up to 14-15 years of age (prior to the COVID-19 pandemic), and additional waves conducted remotely during the COVID-19 pandemic, with timing of future waves into early adulthood under consideration. The Child Health CheckPoint module was a once-off physical health and biomarkers assessment, nested between LSAC's 10-11 year and 12-13 year waves, for 1,874 B-cohort families[23]. Families either attended an assessment centre or received a home visit and provided biospecimens for molecular analyses and DNA extraction. CheckPoint comprises mainly (85%) participants of European ancestry[24].

Written informed consent and assent were obtained from all parents and children. The Australian Institute of Family Studies (AIFS) Ethics Committee approved each wave of LSAC and the CheckPoint (14-26), with additional approval from Melbourne's Royal Children's Hospital Human Research Ethics Committee (33225D).

#### MoBa

The Norwegian Mother, Father and Child Cohort Study (MoBa) is a population-based pregnancy cohort study conducted by the Norwegian Institute of Public Health. Participants were recruited from all over Norway from 1999-2008[25]. Women consented to participate in 41% of pregnancies. Blood samples were obtained from both parents during pregnancy and from mothers and children (umbilical cord) at birth[26]. The cohort includes approximately 114,500 children, 95,200 mothers and 75,200 fathers. The establishment of MoBa and initial data collection was based on a license from the Norwegian Data Protection Agency and approval from the Regional Committees for Medical and Health Research Ethics. The MoBa cohort is currently regulated by the Norwegian Health Registry Act. The present analyses used the genetic data from a subsample of 98,110 individuals genotyped in 10 batches[27,28]. Only 16,768 children with a measure of prosocial behaviour (measured by the SDQ) at three years of age were included in the current study. The Medical Birth Registry (MBRN)[29], a national health registry containing information about all births in Norway from 1967 onwards, was used to determine sex assigned at birth.

Pregnant women provided written consent before enrolment in the study. The current study was approved by the Regional Committees for Medical and Health Research Ethics (2016/1702).

#### MUSP

The MUSP (Mater Misericordiae Mothers' Hospital-University of Queensland Study of Pregnancy) study is of 7,223 women recruited early in pregnancy over the period 1981-1983, and the live singleton children to whom they subsequently gave birth[30,31]. There were follow-ups of the mothers at 5, 14, 21 and 27 years after recruitment. Children were also followed up in the mothers' questionnaire and independently at 21 and 30 years of age. The CBCL and YSR were administered to children at 5, 14 and 21 years of age. The cohort comprises both mothers (up to 27 years after the birth) and children (up to 30 years of age). For more details, see [30,31].

At all data-collection phases of the study, mothers provided written informed consent. The current study was approved by the Mater Hospital Ethics Committee and the University of Queensland Ethics Committee.

#### NFBC1986

NFBC1986 is a prospective longitudinal pregnancy-offspring cohort study consecutively recruited in early pregnancy[32,33]. The population is Finnish and therefore of European ancestry. All pregnant women whose expected date of birth was between July 1st 1985, and June 30th 1986, in the two northernmost provinces of Finland (Oulu and Lapland) were eligible. Participants comprised 99% of all deliveries occurring in the region during the recruitment period. Altogether, there were 9,362 deliveries, with 9,479 infants of whom 9,432 were live-born. Follow-up data collections occurred during the child's ages of 7, 8 and 15 / 16 years. The most recent data collection, including clinical examinations and questionnaires, was conducted when offspring were 33 years old. Data come from multi-informant postal questionnaires, health records, national register data, and clinical examinations[34]. More information about the cohort can be found here (http://www.oulu.fi/nfbc).

Written informed consent was provided by parents and adolescents. Studies were approved by the ethics committee of the Northern Ostrobothnia Hospital District in Oulu (EETTMK: 108/2017 (15.1.2018)) and the Oulu University, Faculty of Medicine, Oulu, Finland.

#### NTR

The Netherlands Twin Register (NTR, https://tweelingenregister.vu.nl/) is a population-based prospective cohort study from the Netherlands. The Young NTR includes newborn twins and their family members who were born since 1986[35,36]. The NTR focuses on studying individual differences in health, behaviour, development, and lifestyle. Phenotype data were collected longitudinally by surveys. Biological samples (buccal cells and blood) were collected for DNA isolation[37].

Written informed consent was provided by the parents prior to enrolment in the study. The study was approved by the Central Ethics Committee on Research Involving Human Subjects of the VU University Medical Centre, Amsterdam, an Institutional Review Board certified by the U.S. Office of Human Research Protections (IRB number IRB00002991 under Federal-wide Assurance-FWA00017598; IRB/institute codes, NTR 03-180).

#### The Raine Study

The Raine Study is a prospective pregnancy cohort of 2,900 mothers recruited between 1989–1991 (https://www.rainestudy.org.au/)[38]. Recruitment took place at Western Australia's major perinatal centre, King Edward Memorial Hospital, and nearby private practices. Women who had sufficient English language skills, an expectation to deliver at King Edward Memorial Hospital, and an intention to reside in Western Australia to allow for future follow-up of their child were eligible for the study. The primary carers (Gen1) completed questionnaires regarding their respective study child, and the children (Gen2) had physical examinations at ages 1, 2, 3, 5, 8, 10, 14, 17, 18, 20, 22, 27 and 28 years. This study included a subset of the original cohort that had genetic data and were of white European ancestry. All research was performed in accordance with the approved guidelines.

Parents, guardians and young adult participants provided written informed consent either before enrolment or at data collection at each follow-up. Ethics approval for the original pregnancy cohort and subsequent follow-ups was granted by the Human Research Ethics Committee of King Edward Memorial Hospital, Princess Margaret Hospital, the University of Western Australia, and the Health Department of Western Australia.

#### SYS

The Saguenay Youth Study (SYS) is an ongoing, population-based study of adolescents and their parents (N=1,029 adolescents and 962 parents) aimed at investigating trajectories of cardiometabolic and brain health[39]. Participants were recruited through regional high schools. Data collection took place from 2003 to 2012 in adolescents (full) and their parents (partial), and from 2012 to 2015 in parents (full). Phenotyped data of each participant cover a wide range of domains, including cognition, mental health and substance use, diet, physical activity and sleep, and family environment. Participants were genotyped using the Human610-Quad and

HumanOmniExpress BeadChips (Illumina, San Diego, CA). More information is available in the original publication[39].

Written consent of the parents and assent of the adolescents were obtained. The study protocol was approved by the Research Ethics Committee of the Chicoutimi Hospital and the Hospital for Sick Children in Toronto.

#### TEDS

Participants come from the Twins Early Development Study (TEDS)[40,41], which comprises 16,810 twin pairs born in England and Wales between 1994 and 1996 who have been assessed in multiple waves across development from approximately 18 months up until the present. Although there has been some attrition, more than 10,000 twin pairs remain actively involved in the study. The demographic characteristics of TEDS participants and their families are reasonably comparable to those of the population in England and Wales for this birth cohort[40].

Written informed consent was obtained from parents before data collection and from TEDS participants themselves past the age of 18 years. King's College London's Ethics Committee approved the project for the Institute of Psychiatry, Psychology and Neuroscience, PNM/09/10–104.

#### TRAILS

TRAILS (Tracking Adolescents' Individual Lives Survey; https://www.trails.nl/en/home) is a prospective cohort study of Dutch adolescents and young adults, with approximately triennial measurements from age 11 years onwards, which started in 2001[42,43]. TRAILS consists of a general population cohort (TRAILS-pop) and a clinical cohort (TRAILS-CC). Participants of the population cohort were selected from five municipalities in the Northern part of the Netherlands, including both urban and rural areas. Participants of the clinical cohort were selected from children who had been referred to a child psychiatric outpatient clinic in the same area at any time before the age of 11 years. DNA was extracted from blood samples or (in a few cases) buccal swabs. In total, 1,354 participants of the population cohort and 341 participants of the high-risk cohort, all of European ancestry, were genotyped with the Illumina Cyto SNP12 v2 array. For more details, please see [42,43].

All measurements were carried out with participants' adequate understanding and written consent. The study was approved by the Dutch Central Committee on Research Involving Human Subjects (CCMO).

### SUPPLEMENTARY NOTE 2. SOCIAL BEHAVIOUR MEASURES AND ICF ALIGNMENT

#### Phenotype definition

Across the cohorts, dimensions of social behaviour (LPB and PSP) were predominantly assessed with the SDQ[44] and the CBCL[45] (Supplementary Table 3). Both questionnaires are widely used, validated, and standardised instruments assessing psychosocial symptoms for research and clinical practice in both clinical and general population-based settings[46]. The SDQ is a brief screening instrument used to assess psychosocial symptoms across both a problem and a competency scale based on parent, teacher, and self-reports. It was derived from the Rutter questionnaire[47], and updated according to diagnostic criteria put forward in the Diagnostic and Statistical Manual of Mental Disorders IV[48] and the International Classification of Diseases-10[49]. The CBCL is an instrument for the in-depth assessment of child behavioural-emotional problems and competencies available in four versions: parent report (CBCL/6-18 for ages 6 to 18 years[50]; CBCL/1.5-5 for ages 1.5 to 5 years[51]), teacher report (Teacher Report Form, TRF, for ages 6 to 18 years[52]) and adolescent self-report (Youth Self-Report, YSR, for ages 11 to 18 years[53]).

#### LPB questionnaires and items

The LPB domain was assessed with 6 different questionnaires (62 items): Strengths and Difficulties Questionnaire (SDQ, 5 items), Revised Rutter Parent Scale for Preschool Children (RR, 11 items), Early Adolescent Temperament Questionnaire-Revised (EATQR, 6 items), Revised Class Play method of peer assessment (RCP, 7 items), Prosocial Behaviour Questionnaire (PBQ, 11 items), Groupe de Recherche sur l'Inadaptation Psychosociale (GRIP, 10 items), and Multidimensional Peer Nomination Inventory (MPNI, 12 items).

#### PSP questionnaires and items

The PSP domain was assessed with 9 different questionnaires: Strengths and Difficulties Questionnaire (SDQ, 5 items), Diagnostic and Statistical Manual of Mental Disorders IV (DSM-IV, 7 items), Positive Youth Development (PYD, 9 items), Child Behaviour Checklist for 6-18 year olds (CBCL/6-18, 11 items), Youth Self-Report of CBCL (YSR, 11 items), Teacher Report Form of CBCL (TRF, 11 items), Teacher's Checklist of Psychopathology (TCP, 6 items), Child Behaviour Checklist for 1.5-5 year olds (CBCL/1.5-5, 11 items), and Berkeley Puppet Interview (BPI, 5 items). Note that different versions of the CBCL[45] were included, assessing social problems from 6 years onwards: parent report (CBCL/6-18[50]), teacher report (TRF[52] and TCP[54]) and adolescent self-report (YSR[53]). Therefore, we used the CBCL/6-18 to reflect these other 3 questionnaires. Therefore, 6 different questionnaires were used to assess PSP, covering 49 items.

#### ICF code alignment

Dimensions of LPB and PSP were assessed with different questionnaires (Supplementary Table 3). To harmonise measures of social behaviour and align instruments, we linked questionnaire items to the World Health Organisation’s (WHO)  International Classification of Functioning (ICF) (Children & Youth Version)[55]. The ICF is a bio-psycho-social framework designed to capture functioning, defined as the interaction between an individual, their activities and participations, and their contexts. The ICF also includes a comprehensive classification system, comprising nearly 1700 codes across body function, body structure, activity, and participation and environmental domains. Established procedures outlined by the WHO and ICF research branch provide a method to systematically harmonise heterogeneous measures using the taxonomy of the ICF[56–58].

Item interpretation and alignment with ICF-codes followed published protocols[59] and were discussed among authors with expertise in the ICF system and who had received training from the WHO ICF Research Branch on ICF linking.

To calculate the contribution of ICF codes for each questionnaire, we conducted questionnaire-based weighting. Subsequently, we computed the contribution of ICF codes per social domain using domain-based weighting, where we account for the number of observations assessed with each questionnaire.

**Questionnaire-based weighting**. As questionnaires had a different number of items (Supplementary Table 4), we standardised the contribution of ICF codes within each questionnaire, using the following formula:

$$C_{ICF_k}^{(q)} = N_{ICF_k}^{(q)} \Big/ N_{total}^{(q)}$$

- $C_{ICF_k}^{(q)}$ is the contribution of ICF code k within questionnaire q
- $N_{ICF_k}^{(q)}$ is the number of items in questionnaire q that are aligned to ICF code k
- $N_{total}^{(q)}$ is the total number of items in questionnaire q

**Equation 1.** Questionnaire-based weighting

**Domain-based weighting** (weighted by the number of observations): To calculate the overall contribution of each ICF code within a social domain, we aggregated the questionnaire-level contributions, accounting for the number of observations assessed with each questionnaire (Supplementary Table 3), using the following formula:

$$C_{ICF_k}^{domain} = \sum_{q=1}^{Q_{domain}} C_{ICF_k}^{(q)} * \frac{N_{obs_q}}{N_{obs_{domain}}}$$

- $C_{ICF_k}^{domain}$ is the contribution of ICF code k within each social domain
- $C_{ICF_k}^{(q)}$ is the contribution of ICF code k within questionnaire q
- $N_{obs_q}$ number of observations assessed with questionnaire q
- $N_{obs_{domain}}$ number of observations for each social domain
- $Q_{domain}$ is the number of questionnaires for each social domain

**Equation 2.** Domain-based weighting

#### Limitations

All questionnaire items were classified following the linking rules provided by the ICF Research Branch, and in accordance with previous research[56,60], where the assigned ICF code captures characteristics of the questionnaire item's underlying behavioural theme. Given the nuanced nature of item content, multiple ICF codes could apply to the same questionnaire item, reflecting the possibility of diverse interpretations at the item level.

To enhance the consistency and reliability of the coding process, all assigned ICF codes were thoroughly discussed with co-authors who were trained specifically in the ICF classification system. This training supports greater consistency in interpretation across authors who were involved in the ICF code alignment and helps reduce potential bias. Nonetheless, we acknowledge that some variability in interpretation can still affect the results of these analyses, particularly given the inherent complexity of defining, interpreting and classifying social behaviour, which is a challenging topic to this day[61].

### SUPPLEMENTARY NOTE 3. HETEROGENEITY IN SNP EFFECTS AT GENOME-WIDE ASSOCIATED LOCI

We identified six loci (tagged by SNPs rs148774891, rs181817841, rs7490422, rs77125329, rs538645, and rs150685860) associated genome-wide with social behaviour (Table 1, Supplementary Table 6).

**rs148774891**. The top associated locus (tagged by rs148774891, A/G, chromosome 3, $p=2.51 \times 10^{-9}$, $\beta=0.202(SE=0.034)$, EAF=1.29%(SE=0.26%)) located inside an intronic region of the Cell Adhesion Molecule 2 (*CADM2*) gene (Figure 2a) and was associated with self-reported LPB from ages 9 to 14 years. Heterogeneity in SNP effects was captured by the fixed effect for self-report ($\theta_{Self}=0.17(SE=0.032)$, $p=1.2 \times 10^{-7}$, Supplementary Table 7).

**rs181817841 and rs77125329**. Two other loci were associated with parent-reported LPB (tagged by rs181817841 and rs77125329). The SNP rs181817841 (T/G, chromosome 8, $p=3.3 \times 10^{-9}$, $\beta=0.154(SE=0.026)$, EAF=1.25%(SE=0.29%)) was related to parent-reported LPB from ages 3 to 8 years. Variation in genetic effects was explained by fixed effects for social domain, teacher-report, self-report and age, the latter being the strongest moderator ($\theta_{Age}=-0.010(SE=0.003)$, $p=1.3 \times 10^{-3}$, Supplementary Table 7). The SNP rs77125329 (T/C, chromosome 10, $p=5.7 \times 10^{-9}$, $\beta=-0.15(SE=0.026)$, EAF=1.24%(SE=0.31%)) is located inside an intronic region of the Leucine Rich Melanocyte Differentiation Associated (*LRMDA* or *C10orf11*) gene and was associated with $LPB_{EarlyCh}$ and parent-reported LPB (ages 2-5 years). Heterogeneity in SNP effects was accounted for by fixed effects for age, age squared, and, especially, social domain ($\theta_{PSP}=0.10(SE=0.025)$, $p=7.7 \times 10^{-5}$, Supplementary Table 7).

**rs7490422 and rs150685860**. Parent-reported PSP was associated with two loci (tagged by rs7490422 and rs150685860). rs7490422 was associated with parent-reported PSP at ages 4 to 8 years. Heterogeneity in SNP effects at rs7490422 (G/A, chromosome 13, $p=5.37 \times 10^{-9}$, $\beta$=-

0.035(SE=0.006), EAF=35.04%(SE=1.27%)) was explained by fixed effects for self-report ($\theta_{Self}$=0.018(SE=0.007), p=1.1x$10^{-5}$, Supplementary Table 6). rs150685860 (A/G, chromosome 20, p=2.72x$10^{-8}$, β=-0.164(SE=0.030), EAF=2.09%(SE=0.58%)) is located inside an intronic region of the XK Related 7 (*XKR7,* Figure 2g) gene and was linked to parent-reported PSP at ages 2 and 3 years. Heterogeneity in SNP effects was accounted for by fixed effects for self-report and, especially, age and age squared ($\theta_{Age}$=0.009(SE=0.002), p=2.7x$10^{-4}$; $\theta_{Age^2}$=-0.001(SE<0.001), p=2.2x$10^{-4}$, Supplementary Table 7).

**rs538645**. Across all reporters, we identified a locus associated with early childhood PSP (rs538645, T/C, p=1.41x$10^{-8}$, β=-0.044(SE=0.008), EAF=76.43%(SE=1.02%)) though the signal was largely based on parent-report, given the scarcity of teacher- and self-reports at this age. Heterogeneity in genetic effects at this SNP was captured by fixed effects for social domain, age square, but especially age ($\theta_{Age}$=0.003(SE=0.001), p=1.0x$10^{-5}$, Supplementary Table 7).

### SUPPLEMENTARY NOTE 4. BIOLOGICAL ANNOTATION OF GENOME-WIDE ASSOCIATED LOCI

#### Functional annotation and gene mapping

To map the independent SNPs to genes, both positional and eQTL mappings were used, as implemented in FUMA[62]. Positional mapping was carried out using ANNOVAR annotations within FUMA across a window of 10 kb. For eQTL mapping, GWA variants at a locus were screened for eQTL affecting the expression of genes up to 1Mb, either in blood (Blood eQTL browser[63], BIOS QTL browser[64] and GTEx v8[65]) or brain (BRAINEAC[66] and GTEx v8[65]) tissues. Following default settings in FUMA, only SNP-gene pairs were considered that passed a false discovery rate of <5%. Identified variants were annotated for their deleteriousness using CADD scores (i.e. a deleteriousness score, the higher the more deleterious, with a proposed cut-off to indicate deleteriousness at CADD>12.37). Mapped genes were annotated for their intolerance to functional mutations using pLI scores (i.e. intolerance to functional mutations score, the higher the more intolerant, proposed cut-off to indicate intolerance pLI>0.90)[67].

Chromatin interaction mapping was also carried out; however, the identified genes using this mapping were only used for gene-based analysis (GENE2FUNC in FUMA) and not for gene identification. To carry out chromatin interaction mapping, we selected chromatin data from adult and foetal human cortex data[68] as well as brain tissue data (hippocampus, left and right ventricle, and dorsolateral prefrontal cortex)[69]. As per default settings, a cut-off of $p<1x10^{-6}$ was used to assess evidence for chromatin interactions.

Additionally, we conducted a look-up in the GWAS Catalog[70] (downloaded on September 22nd, 2024) and PheWeb[71] (accessed on October 10th, 2024) databases. For the PheWeb look-up, a browser search was conducted per lead SNP. For the GWAS Catalog look-up, we downloaded "all associations v1.0" and conducted two look-up steps. For the first look-up, we queried the entire database for previously reported associations with the six lead SNPs. For the second look-up, we searched the entire database for associations reported for the mapped genes (positional and eQTL mapping). Then, using the *LDlinkR* R package (version 1.4.0)[72] and the Central European population as a reference panel, we explored the LD-$r^2$ between the lead SNP and the reported associations across the mapped genes. All SNPs within a 500kb window around the lead SNP were included.

#### Gene expression patterns, tissue specificity and molecular functions

Genes mapped by positional and eQTL mappings, as well as chromatin interaction mapping, were studied using GENE2FUNC (version 1.3.5) within FUMA. Here, we explored gene expression patterns as well as tissue specificity (differentially expressed genes, DEG) in the GTEx v8 RNA-sequencing[65] (30 broad tissue types and 54 specific tissue types) and the BrainSpan[73] (11 developmental stages and 29 different ages of brain samples) databases. Finally, within FUMA, the mapped genes were tested for overrepresentation in any specific gene sets from WikiPathways[74] (v2023) and GWAS Catalog[70] (version e110_r2023-07-20) databases.

Following a previous analysis pipeline[75], we conducted a look-up of whole-brain cortical expression signatures of genes with potential biological implications using the Allen Human Brain Atlas (AHBA; https://human.brain-map.org, N=6 post-mortem brains, aged 24-58 years)[76]. Following quality assessment using the *abagen* toolbox[77] (v0.1.1), the AHBA provided mRNA data for 15,633 genes and 1,670 samples in cortical tissue, standardised across genes and donors. Samples were then assigned to the FreeSurfer *fsaverage6* standard-space surface template, spatially interpolated and smoothed to create a gene-by-vertex matrix of standardised mRNA expression across the entire cortical surface. Cortical gene expression was displayed using the *fsbrain* R package[78] (*R::fsbrain*, v0.5.5).

Additionally, we extracted gene expression data from the spatio-temporal gene expression data set available from the BrainSpan Atlas of the Developing Human Brain (http://www.brainspan.org/)[79]. This data covers transcriptome profiles of 16 different brain regions within a time frame ranging from embryonic development to late adulthood for males and females. We plotted the 2D heatmaps with brain anatomical structures over time using the *cerebroViz* R package (*R::cerebroViz*, v1.0)[80]. The gene expression values were transformed to Z-scores for better visualisation.

#### Gene-based, gene-set and gene-property analyses

Gene, gene-set, and gene-property analyses were conducted with MAGMA (version 1.08)[81].

In gene analysis, SNPs were mapped to protein-coding genes from Ensembl (build 85) using a window of 1kb upstream and 0kb downstream. A genome-wide, gene-based significance threshold was set to $2.62 \times 10^{-6}$ (0.05/19,039 protein-coding genes tested).

To conduct gene-set analysis, we included 4,767 gene-sets containing 10 to 200 genes from gene ontology (GO) biological processes from the Molecular Signatures Database (MSigDB v2023)[82]. To assess gene-set p-values, MAGMA runs a competitive model accounting for gene size, gene density and a gene-wise mean minor allele count[81]. The multiple testing adjusted p-value threshold for association was set to $1.05 \times 10^{-5}$ (0.05/4,767 gene sets).

Additionally, gene-property analyses were conducted to examine the relationship between identified MAGMA genes and their expression within a particular tissue. The mapped genes, based on gene-based analysis, were tested for differential expression using GTEx v8 RNA-sequencing[65] (30 broad tissue types and 54 specific tissue types) and the BrainSpan[73] (11 developmental stages and 29 different ages of brain samples) databases. The multiple testing threshold was set to $4.03 \times 10^{-4}$ (0.05/124 tests).

#### Results

Here, we report mapped genes (Supplementary Table 8-9), gene-based and gene-set association tests using MAGMA software (Supplementary Figure 5, Supplementary Table 10) as well as gene expression patterns, tissue specificity and molecular signatures of mapped genes (Supplementary Figure 6-Supplementary Figure 11, Supplementary Table 11) using *GENE2FUNC* in FUMA software. In addition, we examined the brain expression patterns of mapped genes in the Allen Human Brain Atlas[76] and BrainSpan samples[73] (Supplementary Figure 12, Supplementary Figure 13).

The six genome-wide associated loci were mapped to 18 genes (8 protein-coding genes) based on positional and eQTL mapping using FUMA[62] (Supplementary Table 8). Additionally, the loci were mapped to 36 other genes (8 protein-coding genes) based on chromatin interaction mapping (Supplementary Table 9). Three of the identified loci (rs148774891, rs150685860 and rs538645) are likely to have a biological impact and, thus, are discussed in detail below.

**rs148774891.** The strongest signal was observed for rs148774891, associated with self-reported LPB from 9 to 14 years. This low-frequency variant (EAF=1.29%) is located within the cell adhesion molecule 2 (*CADM2*) gene and *CADM2* Antisense RNA 1 gene (*CADM2-AS1*) (Figure 2a) on chromosome 3p12.1. *CADM2* has a high score for probability loss-of-function intolerance (pLI=0.98) and is important for synapse organisation. Furthermore, this gene has been previously linked to autism[83,84], with a Simons Foundation Autism Research Initiative (SFARI) database score of 2 (https://gene.sfari.org/database/human-gene/CADM2)[85], risk-taking behaviour[86], smoking and alcohol consumption[87–89], externalising behaviours[90], educational attainment (EA)[91], and general cognitive ability[92]. However, the *CADM2* locus reported within this study (rs1487748919) is novel and unrelated to previous GWAS findings (Supplementary Figure 15). *CADM2* is expressed throughout all BrainSpan developmental stages (Supplementary Figure 6a-b) and all GTEx brain tissues as well as the tibial nerve (Supplementary Figure 6c-d).

Furthermore, the *CADM2* gene is strongly expressed in parietal and occipital areas, including the trans-parietal junction (TPJ) and the posterior superior temporal sulcus (pSTS), which are integral parts of the 'social brain'[93], and also shows heightened cortical expression values (>0.6) in the Allen Human Brain Atlas dataset across the entire cortex (Supplementary Figure 12). Matching the developmental window of the associated trait in BrainSpan data, *CADM2* over-expression peaks in the orbitofrontal cortex at 9-14 years (Figure2c, Supplementary Figure 13), a region important for decision-making in social and reward-based contexts[94].

In addition, we observed heighted expression in the dorsolateral prefrontal cortex in early childhood (Supplementary Figure 13), outside the developmental window of the associated signal. However, the novel lead variant rs148774891 could not yet be related to a detectable eQTL (Supplementary Table 8).

**rs538645.** Positional mapping of the locus on chromosome 11 (rs538645, Figure 2d), associated with early childhood PSP, identified two genes (SET Pseudogene 16: *SETP16*, ENSG00000201535: *Y RNA*) with a CADD score of 14.02, indicating potential deleteriousness. Furthermore, eQTL mapping linked this locus to Archain 1 (*ARCN1,* $p_{eQTL}$=2.67x10$^{-7}$, rs7104519:LD-$r^2$=1, Supplementary Table 9), a high pLI score gene (pLI=1). *ARCN1* is highly expressed across all GTEx tissues and BrainSpan developmental stages (Supplementary Figure 10). Using data from the Allen Human Brain Atlas, we observed that *ARCN1* is predominantly expressed in occipital brain regions, particularly in the primary visual cortex (Supplementary Figure 12). The spatiotemporal expression heatmap for *ARCN1* in BrainSpan showed higher expression in the medulla and cerebellar cortex after birth (Figure 2f, Supplementary Figure 13). The lead SNP at this locus is in LD with SNPs previously reported for vitiligo[95] (rs638893:LD-$r^2$=0.63), depression[96–98] (rs75488969:LD-$r^2$=0.35, rs2187490:LD-$r^2$=0.33), height[99] (rs10892301:LD-$r^2$=0.36) and sex hormone-binding globulin levels[100] (rs78312641:LD-$r^2$=0.38)(Supplementary Figure 15).

**rs150685860.** Another low-frequency GWAS signal (rs150685860, EAF=2.09%), associated with parent-reported PSP at 2-3 years, resides within the XK-related 7 (*XKR7,* Figure 2g) gene on chromosome 20q11.21, a gene with a high pLI score (pLI=0.92) involved in apoptotic processes of the cell. The *XKR7* gene has been associated with body height[101], increased diastolic blood pressure[102] and decreased lean muscle mass[103]. The locus rs150685860 was unrelated to known GWAS signals in this gene (Supplementary Figure 15). In addition to positional mapping, SNPs within the same locus were mapped to *XKR7* via chromatin interaction mapping based on foetal cortex (Supplementary Table 9). Tissue expression analysis revealed that *XKR7* was expressed in cerebellum and cerebellar hemisphere tissues (Supplementary Figure 11). Gene-set analysis via MAGMA for parent-reported PSP GWAS at 2 to 3 years (where the locus was identified, but unrelated to the GWA locus itself) revealed one biological pathway in gene ontology (GO), the O-linked glycosylation of protein via threonine (GO:0018243, $N_{genes}$=10, beta=1.36, SE=0.29, p=$1.29 \times 10^{-6}$, beta/$STD_{geneSet}$=0.03; Supplementary Table 10).

Finally, across the six GWAS, there was no evidence of any gene-based signals passing the genome-wide gene-based significance threshold of $2.62 \times 10^{-6}$ (Supplementary Figure 5). Gene-set analysis in MAGMA did not reveal any association with tissue or age-specific gene expression (Supplementary Table 10), and mapped genes by positional, eQTL or chromatin interaction mapping were not enriched in any tissue or age-specific differentially expressed gene sets (Supplementary Table 11). Lastly, the reported GWA loci were neither related to adult traits within a phenome-wide association study in the UK biobank (Supplementary Figure 14) nor strongly related (LD-$r^2$>0.8) to previous GWAS signals (Supplementary Figure 15).

### SUPPLEMENTARY NOTE 5. GENETIC CFA OF SOCIAL BEHAVIOUR

#### Modelling approach

To identify the overarching genetic influences underlying social behaviour, we studied the latent structure of social behaviour using high-resolution GWAS, allowing for differences in social domain, reporter, and age ($N_{GWAS}$=66). To do so, we carried out a data-driven approach[104] utilising a combination of principal component analysis (PCA), exploratory factor analysis (EFA) and confirmatory factor analysis (CFA), where the CFA model was fitted with genomicSEM[105] software.

First, we investigated the number of genetic factors across the 66 high-resolution GWAS. To do so, we applied PCA of the LDSC-$r_g$ matrix, as estimated with the ldsc() function within the genomicSEM[105] R package (*R::genomicSEM*, v0.05). Eigenvalues of this PCA were used to estimate the number of shared genetic factors according to the Optimal Coordinate criterion (*R::nFactors*, v2.4.1)[106]. This criterion applies a joint Kaiser's rule (retain eigenvalues above 1) and Cattell's scree test (retain eigenvalues before the scree in the eigenvalue plot)[106]. Eigenvalue decomposition of the LDSC genetic correlation matrix identified four genetic factors (Figure 3b).

Second, provided that there is more than one genetic factor, we carried out an EFA in *lavaan* (*R::lavaan*, v0.6-10)[107] to describe the underlying genomic structure, fitted to the LDSC genetic covariance matrix. Factor solutions were estimated using Diagonally Weighted Least Squares (DWLS)[108] algorithm, where inverse weighting was carried out with the inverse of the diagonal of the SE matrix for the LDSC genetic covariance matrix. We tested models that allowed for correlation between factors (oblique geomin rotation) and models that did not allow for correlation between factors (orthogonal varimax rotation).

Third, we carried out a CFA, as informed by the factor structure of the EFA model. In particular, we retained EFA factor loadings explaining more than 0.25% of the heritability of each GWAS

(0.05≤|λ|). Then, we fitted a CFA with and without factor correlations within using the usermodel() function within genomicSEM[105]. Both models were further constrained using a cut-off of 0.05 on unstandardised factor loadings, consistent with the cut-off used in the EFA model. To identify the best-fitting factor structure, we assessed the model fit using three fit indices[109] from the genomicSEM R package[105], including the relative goodness-of-fit using the Akaike information criterion (AIC, the lower the better), the absolute goodness-of-fit using the Comparative Fit Index (CFI, optimal fit with values above 0.95), and the Standardised Root Mean Squared Residual (SRMR, optimal fit with values below 0.08).

#### Results

Our data-driven modelling approach identified a 4-factor model with correlated factors (Figure 3c-g, Supplementary Tables 13-15, Supplementary Figure 18) and a good model fit (CFI=0.98, SRMR = 0.05). The four identified factors explained genetic variation across (F1) parent-reported LPB and PSP, (F2) LPB, (F3) teacher-reported LPB and PSP, and (F4) PSP.

Correlations between factors (Figure 3c) were low to moderate (ranging from -0.33(SE=0.13) to 0.60(SE=0.15)), being highest between F2 (LPB) and F3 (teacher-reported LPB and PSP)($r_g$=0.60(SE=0.15)) and between F1 (parent-reported LPB and PSP) and F2 (LPB) ($r_g$=0.55(SE=0.19)).

### SUPPLEMENTARY NOTE 6. TARGET SAMPLES USED IN POLYGENIC SCORING ANALYSIS

#### Millennium Cohort Study (MCS)

The Millennium Cohort Study (MCS) is a nationally representative family-based study from the United Kingdom with data on childhood and adolescent social behaviour, which includes children born at the turn of the new century[110,111]. The study contains 18,552 families (18,827 children). Data has been collected at 9 months, as well as at 3, 5, 6, 11, 14 and 17 years old. Written consent was provided by the parents of the children participating in this study (https://cls.ucl.ac.uk/wp-content/uploads/2017/07/MCS-Ethical-review-and-consent-Shepherd-P-November-2012.pdf). We received ethical approval to access and analyse pre-collected de-identified genotype and phenotype data from this cohort from the Radboud University Ethics Committee Social Science (ECSW-2018-160).

Social behaviour measures of PSP and LPB in the MCS were assessed with the Strengths and Difficulties Questionnaire (SDQ)[44] using parent (at 3, 5, 7, 11, 14 and 17 years old), teacher (at 7 and 11 years old) and self-reports (at 17 years old)[112–117]. Descriptive data of the included phenotypes are available in Supplementary Table 16.

Genotyping was performed using the Infinium Global Screening Array-24 v.1.0. After individual and variant quality control (QC), we included in this study 6,541 unrelated individuals (50% males) of European ancestry with genetic and phenotype information available. Imputation was carried out using the Sanger imputation server (https://imputation.sanger.ac.uk), pre-phasing using EAGLE2, PBWT imputation pipeline, and HRC r1.1 as reference panel. Subsequently, best-guess genotypes were constructed, restricting the variants to high-quality imputed SNPs ($N_{SNPs}$=6,947,751, INFO>0.8; 95%-posterior genotyping probability>0.9; MAF>0.5%).

#### 1958 birth cohort (1958BC)

The 1958 birth cohort, also known as the National Child Development Study (NCDS), is a longitudinal study in the United Kingdom of 17,000 children born in a single week in 1958[118,119]. Oral or written consent was provided by the parents of the children participating in this study (https://cls.ucl.ac.uk/wp-content/uploads/2017/07/NCDS-Ethical-review-and-Consent-2014.pdf). We received ethical approval to access and analyse pre-collected de-identified genotype and phenotype data from this cohort from the Radboud University Ethics Committee Social Science (ECSW-2018-160).

A composite score for teacher-reported PSP at 16 years old was constructed by combining two teacher-reported items assessed at 16 years: “Not much liked by other children” and “Does things on own, rather solitary” (Cronbach’s alpha = 0.53). Items were scored using a 3-point Likert scale ("Does not apply"; "Applies somewhat"; "Certainly applies"). Both items were normalised and summed up to create the composite score. Descriptive data of the included phenotype are available at Supplementary Table 16.

Genotyping was performed using the Illumina 1.2M and Infinium HumanHap 550K v3 arrays. After individual and variant quality control (QC), we included 5,349 unrelated individuals (50% males) of European ancestry with genetic and phenotype information available. Genotype imputation was conducted using the Sanger imputation server (https://imputation.sanger.ac.uk), pre-phasing using EAGLE2, PBWT imputation pipeline, and HRC r1.1 as reference panel. Subsequently, best-guess genotypes were constructed, restricting the variants to high-quality imputed SNPS (NSNPs=7,302,748, INFO>0.8; 95%-posterior genotyping probability>0.9; MAF>0.5%).

#### Health and Retirement Study (HRS)

The Health and Retirement Study[120] is a longitudinal study including around 20,000 individuals over the age of 50 in the United States and including information on income, later-life health outcomes, and cognition, among others. Follow-up interviews are conducted every two years. Consent was provided by respondents before saliva collection. We received ethical approval to access and analyse pre-collected de-identified genotype and phenotype data from this cohort from the Radboud University Ethics Committee Social Science (ECSW-2018-160).

Measures from the Functional Limitations and Helpers (Respondent) and Leave-Behind (Respondent) questionnaires from the HRS Core 2010 were used to construct composite LPB and PSP scores. These scores aimed to match items of the peer problems and prosocial behaviour subscales from the SDQ[44]. Descriptive data of the included phenotypes are available at Supplementary Table 16.

To create a composite PSP score, we matched self-reported variables in the HRS to three of the five items of the peer problems SDQ subscale: “Rather solitary, tends to play alone” (11 variables: MLB020A-K; Cronbach’s alpha = 0.89), “Has at least one good friend” (8 variables: MLB018 and MLB016A-G; Cronbach’s alpha = 0.69), and “Generally liked by others” (1 variable: MLB019C).

- To generate a score for the item “Rather solitary, tends to play alone”, we used variables MLB020A-K, assessed with a 3-point Likert scale (“Often”; “Some of the time”; “Hardly ever or never”), and measuring how much of the time the person feels: lack of companionship (MLB020A), left out (MLB020B), isolated from others (MLB020C), ‘in tune’ with the people around them (MLB020D), alone (MLB020E), that there are people they can talk to (MLB020F), that there are people they can turn to (MLB020G), that there are people who really understand you (MLB020H), that there are people they feel close to (MLB020I), that they are part of a group of friends (MLB020J), that they have a lot in common with the people around them (MLB020K). Variables MLB020A, MLB020B, MLB020C and MLB020E were reverse-coded, and all variables were normalised and summed.

- To generate a score for the item “Has at least one good friend”, we included the variable MLB018 (“How many of your friends would you say you have a close relationship with?”) as well as variables MLB016A-G, assessed with a 4-point Likert scale (“A lot”, “Some”, “A little”, “Not at all”) to answer the following questions about friends: “How much do they really understand the way you feel about things?” (MLB016A), “How much can you rely on them if you have a serious problem?” (MLB016B), “How much can you open up to them if you need to talk about your worries?” (MLB016C), “How often do they make too many demands on you?” (MLB016D), “How much do they criticize you?” (MLB016E), “How much do they let you down when you are counting on them?” (MLB016F), “How much do they get on your nerves?” (MLB016G). The variable MLB018 was recoded into 4 (if 0 friends) or 1 (if one or more friends), and variables MLB016D, MLB016E, MLB016F and MLB016G were reverse-coded. The final item score was created by normalising all variables and summing these.
- To generate a score for the item “Generally liked by others” we used variable MLB019C (“How much you agree or disagree with the following statement: No one cares much what happens to you.”), which was assessed with a 6-point Likert scale (“strongly disagree”, “somewhat disagree”, “slightly disagree”, “slightly agree”, “somewhat agree”, “strongly agree”).

Subsequently, we normalised the 3 items and summed them to create a composite PSP score with a Cronbach’s alpha of 0.64.

To create a total LPB score, we matched self-reported variables in the HRS to one of the five items of the prosociality SDQ subscale: “Often volunteers to help others” (4 variables: MLB001C, MLB001D, MG086.recoded [4 variables: MG086, MG195-197], MG198.recoded [4 variables: MG198, MG199-201]). The variables MLB001C (“How often do you do volunteer work with children or young people?”) and MLB001D (“How often do you do any other volunteer or charity work?") were assessed using a 7-point Likert scale ("daily", "several times a week", "once a week", "several times a month", "at least once a month", "not in the last month", "never/not relevant"). The latter measures MG086.recoded ("How much time in the past 12 months have you spent doing volunteer work for religious, educational, health-related or other charitable organizations?", variables MG086, MG195-197) and MG198.recoded ("How much time in the past 12 months have you spent helping friends, neighbours, or relatives who did not live with you and did not pay you for the help?", variables MG198, MG199-201) are recoded variables created following the protocol previously described by Primes and Fieder[121] (see Table 4 in original publication) with 5 possible values ("0 hours", "1 to 50 hours", "51 to 100 hours", "101-200 hours", "more than 200 hours". The composite LPB score was created by normalising the four variables and summing them up (Cronbach's alpha=0.72).

Genotyping was performed using the Illumina HumanOmni2.5-4 v1.0 and Illumina HumanOmni2.5-8 v1.0 arrays. After individual and variant quality control (QC), we included in this study 4,926 unrelated individuals (43% males) of European ancestry with genetic and phenotype information available. Imputation was carried out using the Sanger imputation server (https://imputation.sanger.ac.uk), pre-phasing using EAGLE2, PBWT imputation pipeline, and HRC r1.1 as reference panel. Subsequently, best-guess genotypes were constructed, restricting the variants to high-quality imputed SNPS ($N_{SNPs}$=7,398,707, INFO>0.8; 95%-posterior genotyping probability>0.9; MAF>0.5%).

#### Drakenstein Child Health Study (DCHS)

The Drakenstein Child Health Study (DCHS) is a longitudinal birth cohort study from Cape Town (South Africa) aiming to investigate the impact of early life exposures on the development of health or disease[122]. The study contains 1,143 pregnant women, who were enrolled in public sector clinics from 2012 to 2015, and have followed mother-child pairs through childbirth, childhood and adolescence, with very high cohort retention. Written consent was provided by the parents of the study and is annually renewed. The study was approved by the Ethics Committee of the Faculty of Health Sciences, University of Cape Town, by Stellenbosch University and the Western Cape Provincial Research Committee.

Social behaviour was assessed with either the CBCL[50,51] (5 phenotypes: parent-reported PSP at 2, 4, 5, 7, 8 years) or the SDQ[44] (4 phenotypes: parent-reported LPB and PSP at 7 and 8 years). Descriptive data of the included phenotypes are available in Supplementary Table 16.

Genotyping was performed using the Illumina Infinium PsychArray and Global Screening Array-24 BeadChip platforms. After individual and variant quality control (QC), we included in this study 958 unrelated individuals (48.7% males) of admixed African ancestry with genetic and phenotype information available. Imputation was carried out using the Michigan Imputation Server. Subsequently, best-guess genotypes were constructed, restricting the variants to high-quality imputed SNPs ($N_{SNPs}$=4,852,842, INFO>0.8; 95%-posterior genotyping probability>0.9; MAF>0.05).

### SUPPLEMENTARY NOTE 7. POLYGENIC PREDICTION AND ACCURACY

#### Overview

Using polygenic scores (PGS) from social behaviour GWAS, we studied the association with social phenotypes in three independent European-ancestry samples (Millennium Cohort Study, MCS[110], and 1958 birth cohort, 1958BC[118], Health and Retirement Study, HRS[120]) and one independent African-ancestry sample (Drakenstein Child Health Study, DCHS[122]) to assess the (i) predictive ability, (ii) predictive accuracy, (iii) broad phenotypic transferability and (iv) cross-ancestry transferability of the social behaviour GWAS.

To correct for multiple testing, we estimated the number of independent measures using Matrix Spectral Decomposition[123] based on the phenotypic correlation matrix. This resulted in a threshold of $p<0.0037$ (0.05/14 independent measures) in the MCS, $p<0.025$ (0.05/2 independent measures) in the HRS, and $p<0.0064$ (0.05/8 independent measures) in the DCHS. In the 1958BC, no correction was required, given that only one measure was studied.

#### Confirming the predictive ability of the social behaviour GWAS with the MCS cohort

We validated the derived GWAS by matching the social behaviour GWAS (i.e. discovery GWAS) to 18 social phenotypes (ages 3 to 17 years, parent-/teacher-/self-reported) from the MCS (i.e. target sample), assessed with the Strengths and Difficulties Questionnaire (SDQ)[44] (Supplementary Table 16, Supplementary Note 6). Matching was carried out according to social domain, reporter, and age; for instance, the PGS of parent-reported LPB at 2 years predicting parent-reported LPB at 2 years in the MCS.

We restricted our analysis to the PGS of the 18 discovery GWAS matching the 18 MCS phenotypes. Discovery GWAS predicted their matching target phenotype with positive effective sizes throughout (incremental-$R^2$=0.18-1.13%, β=0.08(SE=0.019)-0.19(SE=0.024), Supplementary Figure 19). All associations passed the multiple-testing threshold (p<0.0037), except for parent-reported PSP at 3 years.

#### Evaluating the predictive accuracy of the social behaviour PGS within the MCS cohort

The predictive accuracy of social behaviour PGS was assessed using 66 PGS generated from high-resolution social behaviour GWAS and 18 MCS social behavioural outcomes.

We evaluated the predictive accuracy of PGS as the percentage of PGS associations from matching discovery-target pairs showing larger associations (incremental $R^2$) than non-matching pairs (Supplementary Table 18-Supplementary Table 19), for associations passing the multiple testing threshold (p<0.0037).

For domain-matched PGS compared to non-matched PGS, the signal was considered accurate if the incremental $R^2$ was higher for the matched PGS. Here, we observed that 81.6% of predictions (279/342) are accurately predicted for PSP outcomes and 45.6% of predictions (156/342) for LPB outcomes (Supplementary Table 18).

Similarly, for reporter-matched PGS compared to non-matched PGS, the signal was considered accurate if the incremental $R^2$ was higher for the matched PGS. For reporter-matched PGS, the predictive accuracy was highest for parent-report (65.53%) and teacher-report (66.07%) outcomes, followed by self-report outcomes (38.8%). For both domain- and reporter-matched PGS, the predictive accuracy increased even further (parent-report outcomes: 88.6%; teacher-report outcomes 71.4%; self-report outcomes: 50%), although the predictive accuracy for self-reported LPB using self-report PGS was poor, and the outcome was better captured by parent-report PGS (Supplementary Table 19).

Next, comparing the largest PGS signal (in incremental-$R^2$) with the fully matching PGS with respect to the target phenotype, we observed, for associations passing the multiple-testing threshold, 89% domain-matched (16/18), 78% reporter-matched (14/18) and 78% age-matched (within a 5-year age window; 14/18) signals capturing contextual information of the target phenotype, with similar magnitude (Figure 4, Supplementary Table 18).

#### Assessing the transferability of the social behaviour PGS within the 1958BC and HRS cohorts

In addition to the MCS cohort, we tested the transferability of social behaviour PGS to broadly defined social measures assessed with non-standardised instruments in early and late adulthood. To do so, we generated social behaviour PGS in two other population-based cohorts, including the UK 1958 British birth cohort (1958BC)[118] ($N_{ind}$=4,195, Supplementary Table 20) with measures of adolescent social behaviour (teacher-reported PSP at 16 years) and the US Health and Retirement Study (HRS)[120] ($N_{ind}$=4,788, Supplementary Table 21) with measures of late-adult social behaviour (HRS, self-reported PSP and LPB at 69 years).

In the 1958BC, social behaviour PGS predicted teacher-reported PSP at 16 years, with the largest predictive ability reaching 0.22% ($p=1.9 \times 10^{-3}$) with matching target phenotype and discovery GWAS, aligned according to social domain and reporter and age (Supplementary Table 20).

In the HRS, we predicted self-reported PSP and LPB at 69 years. The largest amount of variance explained by the social behaviour PGS was observed for adolescent PSP predicting self-reported PSP at 69 years (incremental $R^2$=0.22%, p=0.002, Supplementary Table 21). Note that for the HRS, phenotypic matching target and discovery GWAS was not possible, since the age range of the discovery GWAS did not include late adult social behaviour, but the largest amount of variance was explained by $PSP_{Adol}$ PGS, the largest powered PSP GWAS for later-life in our study.

#### Examining the cross-ancestry transferability of the social behaviour PGS with the DCHS cohort

We examined the extent to which genetic influences in social behaviour are transferable to non-European ancestries by predicting parent-reported social phenotypes in the childhood African-ancestry Drakenstein Child Health Study (DCHS) cohort[122], assessed with the SDQ[44] or CBCL[50] instruments (Supplementary Table 22).

Matching social domains and development stages, social behaviour GWAS PGS predict parent-reported LPB at ages 7 ($LPB_{MidCh}$ GWAS, incremental-$R^2$=0.95%, p=0.007) and 8 ($LPB_{MidCh}$ GWAS, $\beta$=0.26(SE=0.091), p=0.0046, incremental-$R^2$=0.89%) years.

### SUPPLEMENTARY NOTE 8. GENETIC CFA ACROSS SOCIAL BEHAVIOUR AND NEURODEVELOPMENTAL AND NEUROPSYCHIATRIC CONDITIONS

We studied the multivariate genomic dimensions across five NNCs and medium-resolution social behaviour GWAS. Specifically, we studied the meta-regression-derived medium-resolution social behaviour GWAS allowing for differences across social domain and developmental stages (6 GWAS: $LPB_{EarlyCh}$, $PSP_{EarlyCh}$, $LPB_{MidCh}$, $PSP_{MidCh}$, $LPB_{Adol}$, $PSP_{Adol}$). Social behaviour GWAS in early adulthood were excluded as underpowered (Z-scores < 3, Supplementary table 12). In addition, we selected five clinically-defined NNCs, as studied for genetic correlation analysis (Supplementary Table 23). We set the population prevalence for ADHD, autism, MDD to 0.05[124], 0.01[125], 0.16[126], respectively, for schizophrenia to ~0.01[127], and for BD to 0.03[128].

To identify the best-fitting model, we applied a data-driven modelling approach, as described above (Supplementary Note 5)[104]. We constructed genetic CFA models using the genomicSEM R package (*R::genomicSEM*, v0.05)[105]. We studied CFA models with both correlated and uncorrelated factors, as two or more different models can sometimes explain the genetic inter-relationship equally well (Supplementary Table 31-Supplementary Table 34). Both models were further constrained using a cut-off of 0.05 for unstandardised factor loadings, consistent with the cut-off used in the EFA model.

We focused on the constrained model only when the constrained model showed an improved model fit compared to the original (unconstrained) model (Supplementary Table 32). The final model (Figure 7b) was selected based on the best model fit indices (CFI=0.986, SRMR=0.069). All fit indices for all the models are presented in Supplementary Table 32.

The first factor explained shared variation across schizoaffective domains ($F_{SA}$), with the largest factor loading observed for BD ($\lambda$=0.44(SE=0.03)). Similar to the pattern observed for the bivariate genetic correlations (Figure 7a), we identified a factor driven by ADHD ($F_{ADHD}$, λ=0.37(SE=0.08)). This factor shared genetic variation with MDD and PSP GWAS across all developmental stages. The third factor was related to autism ($F_{Autism}$, λ=0.34(SE=0.06)), sharing variation with the LPB GWAS across all developmental stages and, weakly, with schizophrenia. $F_{ADHD}$ and $F_{Autism}$ were correlated with each other (r=0.59(SE=0.13)). The final factor ($F_{SB}$) explained shared variation across social behaviour GWAS only, particularly capturing the negative association between early PSP (λ=-0.11(SE=0.04)) and adolescent LPB (λ=0.17(SE=0.03)).

We further characterised the factors by mapping EA onto the structure of the best-fitting genetic CFA model, both as a sensitivity analysis and to help disentangle the complex relationships between mental health and social behaviour (Supplementary Table 34). EA can act as a proxy for both cognitive traits and socio-economic status. This model had a good fit (CFI=0.973, SRMR=0.069), albeit not as strong as the best-fitting genetic CFA model. In the EA-mapped model (Supplementary Figure 28, Supplementary Table 34), EA is captured by the schizoaffective factor ($F_{SA}$), the ADHD factor ($F_{ADHD}$), and the autism factor ($F_{Autism}$) but not the social behaviour factor ($F_{SB}$; λ=0.02(SE=0.03)). Furthermore, mapping EA increased the correlation between $F_{Autism}$ and $F_{ADHD}$ (r=0.66(SE=0.07)). Notably, once EA was mapped, ADHD (λ=-0.16(SE=0.04)) was also explained by $F_{Autism}$, and, thus, inversely linked to EA (λ=0.3(SE=0.07)). In contrast, autism (λ=0.34(SE=0.04)) was positively associated with EA via the same factor ($F_{Autism}$).

### SUPPLEMENTARY NOTE 9. ANALYSIS PLAN FOR THE GENOME-WIDE META-REGRESSION GWAS OF SOCIAL BEHAVIOUR IN EAGLE

#### Overview

Prior to analysis, each cohort received the Standard Operating Procedure (SOP) for the "meta-GWAS of low prosociality and problematic peer relationships". This SOP was embedded within the behaviour and cognition working group of the EArly Genetics and Life Course Epidemiology (EAGLE) consortium[129] (https://www.eagle-consortium.org/working-groups/behaviour-and-cognition/social-behaviour/). The SOP contained information on (1) background and state-of-the-art, (2) phenotype information, (3) meta-analysis strategy, (4) genotype preparation, (5) phenotype preparation, (6) association analysis, and (7) results formatting.

#### Genotype preparation

We recommended specific filtering steps for the pre-imputation quality control (QC) of the sample and genotyped SNPs. Individuals should be excluded provided that they have a mismatch in reported and genetic sex, they have a missingness above 3%, relatedness above IBD>0.125 (for population-based cohorts only), or non-European genetic ancestry. Single variants should be excluded if they have a missingness above 5%, deviate from Hardy-Weinberg (HWE p-value $<5x10^{-7}$) or are rare (MAF < 1%). After the sample and SNP QC, each cohort imputed their genetic data against the Haplotype Reference Consortium version 1.1 (HRCr1.1) panel[130] or the 1000 Genomes Project Phase 3[131] as reference panel using either Sanger[130,132] (https://imputation.sanger.ac.uk/) or Michigan[133] (https://imputationserver.sph.umich.edu) imputation server (Supplementary Table 37). No post-imputation QC was required.

#### Phenotype preparation

To ensure a standardised procedure across cohorts, we provided a template script which prepared the input genetic and phenotypic data for the association analysis, as well as creating an overview report of the phenotypic measures (phenotype histograms, age histograms, phenotype descriptives, phenotypic correlations). During this step, phenotypic scores will be Z-standardised based on the grand mean, which means that measures assessed with the same questionnaire (e.g. across sexes, ages and/or raters) will be Z-standardised together.

#### Association analysis

Once the genetic and phenotypic data were prepared, the cohorts conducted the association analysis. Association analyses were carried out with SNPTEST[134] or ProbABLE[135] software in cohorts of unrelated individuals and with GEMMA[136] software in samples of related individuals. Example code for SNPTEST and GEMMA was provided within the SOP. In the Longitudinal Study of Australian Children (LSAC) cohort, due to the availability of best-guess genotypes with INFO>0.3 only, association analyses were performed using PLINK 1.9[137] software (Supplementary Table 37).

For each phenotype, three GWAS were run, one for combined sexes (Equation 3), and for QC purposes (Methods) two sex-stratified GWAS (females only, males only, Equation 4).

$$Y_Z = \beta_0 + \beta_{age} * age + \beta_{age^2} * age^2 + \beta_{\mathrm{PCs}} * (PC_1, PC_2, \dots) + \beta_{sex} * sex + \boldsymbol{\beta_{SNP}} * \boldsymbol{SNP}$$

**Equation 3.** Linear model for GWAS containing both males and females

$$Y_Z = \beta_0 + \beta_{age} * age + \beta_{age^2} * age^2 + \beta_{\mathrm{PCs}} * (PC_1, PC_2, \dots) + \boldsymbol{\beta_{SNP}} * \boldsymbol{SNP}$$

**Equation 4.** Linear model for sex-stratified GWAS.

#### Results formatting

We provided the guidelines that each cohort had to follow before sharing their GWAS summary statistics. SNP positions had to be in build 37, data should be aligned to the forward (+) strand, the missing value indicator should be “NA”, exact column names and their format (numeric, character...). Specifically, each cohort shared a tab-delimited file with 15 columns: SNP identifier, chromosome, position, strand, effect allele, non-effect allele, HWE p-value, effect allele frequency, sample size, effect size, standard error of effect size, uncorrected two-sided p-value, imputed (0 if genotyped, 1 if imputed), imputation accuracy measure, and info score. File names were based on the following naming scheme: COHORT_DOMAIN_REPORTER_AGE_SEX_DATE.

### SUPPLEMENTARY NOTE 10. COMPARISON OF META-REGRESSION GWAS WITH CONVENTIONAL META-GWAS APPROACHES

#### Conventional meta-analysis approach

As part of sensitivity analyses, we compared the genome-wide random-effects meta-regression with a conventional meta-analysis approach, using multivariate N-GWAMA software[138], where observations are studied under the assumption of no contextual heterogeneity and sample overlap is accounted for using cross-trait linkage disequilibrium (LD) score intercepts. To do so, we employed a three-step approach (see Supplementary Figure 33 for a visual representation).

##### Step 1: Seed-based fixed-effects meta-analysis in METAL

Cohort-level summary statistics for LPB and PSP (Supplementary Table 2) were separately grouped into four developmental windows according to the age they were assessed: early childhood (2-5 years), mid childhood (6-11 years), adolescence (12-17 years), and young adulthood (18-29 years) and included repeated observations across age and reporters. Given that we obtained summary statistics across repeated observations from multiple cohorts, each developmental window has sample overlap. Therefore, we further divided each developmental bin into multiple seeds (where a seed comprises only summary statistics from independent observations, i.e. no sample overlap). In this context, we ensured that sample sizes for individual cohort-based summary statistics and the total number of included measures were represented equally across seeds. The number of seeds and the N range for each developmental window are shown in Supplementary Figure 33.

Fixed-effect meta-analyses were performed across the cohort-level summary statistics within each seed using METAL software[139] (v2011-03-25; METAL). METAL combines effect size estimates while weighting by the inverse of the corresponding standard error. Analyses were performed without correction for genomic control, given that the following GWAS steps rely on LDSC regression[140] and correlation[141] that adjust for population stratification.

#### Step 2: Developmental window-based meta-analysis in multivariate N-GWAMA

Subsequently, for each social domain, within each developmental window (early childhood, mid childhood, adolescence and young adulthood), seed-based summary statistics derived with METAL were meta-analysed using N-weighted GWAMA software[142] (v.1_2_6; N-GWAMA). Therefore, within this step, we generated 8 GWAS: $LPB_{EarlyCh}$, $LPB_{MidCh}$, $LPB_{Adol}$, $LPB_{Adult}$, $PSP_{EarlyCh}$, $PSP_{MidCh}$, $PSP_{Adol}$, $PSP_{Adult}$.

Assuming a unitary effect of the SNP, the multivariate Z-score for SNP j was calculated following Equation 5. Here, the Z-score of the $j^{th}$ SNP from each GWAS is weighted by sample size and LDSC-$h^2_{SNP}$ (Equation 6). If LDSC-$h^2_{SNP}$ estimates of seed-based GWAS were negative, but 95%CI overlapped with zero, the $h^2_{SNP}$ was fixed to $10^{-6}$. Subsequently, the sum of the weighted Z-scores was then scaled according to the weighted Z-score variances of $V_{ji} = 1$ under the null hypothesis of no effect and the weighted sample overlap between GWAS i and k as given by the LDSC-derived cross-trait intercept (CTI)[143]. If the sample overlap between seed-based GWAS could not be estimated using LDSC-$r_g$ we set the CTI to $10^{-6}$. Note that, to accurately account for sample overlap, the true number of independent observations across repeated measures would be needed, which is, however, difficult to obtain.

$$Z_{multi\ j} = \frac{\sum_{i=1}^{P}\left(w_{ji} * z_{ji}\right)}{\sqrt{\sum_{i=1}^{P}\left(w_{ji} * V_{ji}\right) + \sum_{i=1}^{P}\sum_{k=1}^{P}\left(\sqrt{w_{ji} * w_{jk}} * CTI_{i,k}\right)\ for\ i \neq k}}$$

**Equation 5.** Computation of the Z score within multivariate N-GWAMA. P indicates the number of GWAS included in the meta-analysis for SNP j.

$$w_{ji} = \sqrt{N_{ji} h^{2}_{SNP,i}}$$

**Equation 6.** Sample size weighting. The Z-score of SNP j from GWAS i was weighted, accounting for the GWAS sample size for SNP j and the LDSC $h^2_{SNP}$ for GWAS i.

Step 3: Social domain-based meta-analysis in multivariate N-GWAMA

Lastly, within each social domain, we meta-analysed the N-GWAMA-generated developmental window-based GWAS (step 2 above, 4 GWAS per social domain) using N-weighted GWAMA software[142] (v.1_2_6; N-GWAMA). Therefore, we generated two social domain-based GWAS: $LPB_{All}$ and $PSP_{All}$. As in step 2, we checked for negative $h^2_{SNP}$ estimates and for missing CTI, but no adjustment was required since both parameters were estimated by LDSC for all 8 developmental-window GWAS.

##### Results

We conducted a genome-wide conventional meta-GWAS using fixed-effect METAL[144] and multivariate N-GWAMA[138], an alternative approach to create $LPB_{All}$ and $PSP_{All}$ GWAS (Supplementary Figure 33). Subsequently, we computed LDSC $h^2_{SNP}$ and $r_g$ as well as the LDSC $r_g$ between the results of the meta-regression GWAS and the conventional meta-GWAS approach. Finally, we examined the pattern of overlap with population-based traits and NNCs (Supplementary Table 23) and compared the results across methods.

Notably, magnitude and strength of $h^2_{SNP}$ estimates decreased when adopting a conventional approach ($LPB_{All}$-$h^2_{SNP}$=1.2(SE=0.24), Z-score=5.2; $PSP_{All}$-$h^2_{SNP}$=1.2(SE=0.22), Z-score=5.3) compared to a meta-regression approach ($LPB_{All}$-$h^2_{SNP}$=3.6(SE=0.47)%, Z-score=7.6; $PSP_{All}$-$h^2_{SNP}$=3.1(SE=0.56)%, Z-score=5.5) (Supplementary Figure 29). Genetic correlations across low-resolution social domains derived with either method were strong, both for $LPB_{All}$ ($r_g$=0.75(SE=0.043)) and $PSP_{All}$ ($r_g$=0.89(SE=0.043))(Supplementary Figure 29), and genetic correlation patterns with population-based traits and NNCs were highly consistent (Supplementary Figure 30, Supplementary Table 36). It is noteworthy that LDSC intercepts, which capture the mean inflation due to confounding bias[145], were below one for all meta-regression GWAS, suggesting an (overly) conservative approach when accounting for sample overlap (Supplementary Table 12).

### SUPPLEMENTARY NOTE 11. COHORT ACKNOWLEDGEMENTS

#### 1958 birth cohort

We are grateful to the Centre for Longitudinal Studies (CLS), UCL Social Research Institute, for the use of these data and to the UK Data Service for making them available. However, neither CLS nor the UK Data Service bear any responsibility for the analysis or interpretation of these data.

#### ABCD-NL

We thank all participating hospitals, obstetric clinics, general practitioners and primary schools for their assistance in implementing the ABCD study (Amsterdam Born Children and their Development). We also gratefully acknowledge all the women and children who participated in this study for their cooperation. The ABCD study (Amsterdam Born Children and their Development) has been supported by grants from the Netherlands Organisation for Health Research and Development (ZonMW), the Netherlands Heart Foundation, and Sarphati Amsterdam. Genotyping was funded by the BBMRI-NL grant CP2013-50. T.G.M. Vrijkotte was supported by ZonMW (TOP 40–00812–98–11010).

#### ABCD-US

Data used in the preparation of this article were obtained from the Adolescent Brain Cognitive DevelopmentSM (ABCD) Study (https://abcdstudy.org), held in the NIMH Data Archive (NDA). The ABCD Study® is supported by the National Institutes of Health and additional federal partners under award numbers U01DA041048, U01DA050989, U01DA051016, U01DA041022, U01DA051018, U01DA051037, U01DA050987, U01DA041174, U01DA041106, U01DA041117, U01DA041028, U01DA041134, U01DA050988, U01DA051039, U01DA041156, U01DA041025, U01DA041120, U01DA051038, U01DA041148, U01DA041093, U01DA041089, U24DA041123,

U24DA041147. A full list of supporters is available at https://abcdstudy.org/federal-partners.html. A listing of participating sites and a complete listing of the study investigators can be found at https://abcdstudy.org/consortium_members/. ABCD consortium investigators designed and implemented the study and/or provided data, but did not necessarily participate in the analysis or writing of this report. This manuscript reflects the views of the authors and may not reflect the opinions or views of the NIH or ABCD consortium investigators. The ABCD data repository grows and changes over time. The ABCD data used in this report came from ABCD Curated Annual Release 4.0 (10.15154/1523041; genetics data).

#### ALSPAC

We are extremely grateful to all the families who took part in this study, the midwives for their help in recruiting them, and the whole ALSPAC team, which includes interviewers, computer and laboratory technicians, clerical workers, research scientists, volunteers, managers, receptionists and nurses.

The UK Medical Research Council and Wellcome (Grant ref: 217065/Z/19/Z) and the University of Bristol provide core support for ALSPAC. This publication is the work of the authors, and L.D.H., F.S., D.A. and B.S.P. will serve as guarantors for the contents of this paper. A comprehensive list of grant funding is available on the ALSPAC website (http://www.bristol.ac.uk/alspac/external/documents/grant-acknowledgements.pdf).

#### BREATHE

We acknowledge and thank all the children and their families for participating in the BREATHE project and for their altruism. The research leading to these results has received funding from the European Research Council under the ERC Grant Agreement number 268479 – the BREATHE project.

#### CATSS

CATSS is part of the Swedish Twin Registry (STR), which is managed by Karolinska Institutet and receives funding through the Swedish Research Council under grant no. 2017-00641. The computations/data handling were enabled by resources provided by the Swedish National Infrastructure for Computing (SNIC) at Uppmax, partially funded by the Swedish Research Council through grant agreement no. 2018-05973.

#### COPSAC

All funding received by COPSAC is listed on www.copsac.com. The Lundbeck Foundation (Grant no. R16-A1694), The Ministry of Health (Grant no 903516), Danish Council for Strategic Research (Grant no. 0603-00280B) and The Capital Region Research Foundation have provided core support to the COPSAC research centre. We acknowledge the huge work of the late Professor Hans Bisgaard, who was the founder of COPSAC and was head of the clinical research centre for more than 25 years.

#### DCHS

We thank the mothers and their children for participating in the study, and the study staff, the clinical and administrative staff of the Western Cape Government Health Department at Paarl Hospital and at the clinics for their support of the study.

#### ELVS

The Early Language in Victoria Study (ELVS) was supported by the Australian National Health & Medical Research Council (NHMRC) (project grant no. 237106, 9436958 and 1041947). The authors thank all the participating parents and children in ELVS for their long-term commitment to the study. We acknowledge the contribution of the Victorian Maternal and Child Health nurses who supported the recruitment of this cohort.

#### FinnTwin12

Phenotype and genotype data collection in the FinnTwin12 cohort has been supported by the Wellcome Trust Sanger Institute, the Broad Institute, ENGAGE – European Network for Genetic and Genomic Epidemiology, FP7-HEALTH-F4-2007, grant agreement no. 201413, National Institute of Alcohol Abuse and Alcoholism (grants AA-12502, AA-00145, and AA-09203 to R.J. Rose; AA15416 and K02AA018755 to D. M. Dick; R01AA015416 to Jessica Salvatore) and the Academy of Finland (grants 100499, 205585, 118555, 141054, 264146, 308248 to J.K., and the Centre of Excellence in Complex Disease Genetics (grants 312073, 336823, and 352792 to J.K.).

#### GenR

We gratefully acknowledge the contribution of children and parents, general practitioners, hospitals, midwives, and pharmacies in Rotterdam. The Generation R Study is conducted by the Erasmus Medical Centre in close collaboration with the School of Law and Faculty of Social Sciences at Erasmus University Rotterdam, the Municipal Health Service Rotterdam area, the Rotterdam Homecare Foundation, and the Stichting Trombosedienst & Artsenlaboratorium Rijnmond (STAR-MDC), Rotterdam. The generation and management of GWAS genotype data for the Generation R Study were done at the Genetic Laboratory of the Department of Internal Medicine, Erasmus MC, the Netherlands. We thank Pascal Arp, Mila Jhamai, Marijn Verkerk, Manoushka Ganesh, Lizbeth Herrera and Marjolein Peters for their help in creating, managing and QC of the GWAS database. The general design of the Generation R Study is made possible by financial support from Erasmus MC Rotterdam, Erasmus University Rotterdam, the Netherlands Organisation for Health Research and Development and the Ministry of Health, Welfare and Sport. C. Cecil is supported by the HorizonEurope Research and Innovation Programme (FAMILY, grant agreement no. 101057529, also to A. Neumann; HappyMums, grant agreement no. 101057390) and the European Research Council (TEMPO; grant agreement no. 101039672, also to A. Neumann). A. Neumann and H. Tiemeier are supported by a grant from the Dutch Ministry of Education, Culture, and Science and the Netherlands Organisation for Scientific Research (NWO grant no. 024.001.003, Consortium on Individual Development). The work of H. Tiemeier is further supported by the European Union's Horizon 2020 research and innovation program (Contract grant no. 633595, DynaHealth) and an NWO-VICI grant (NWO-ZonMW: 016.VICI.170.200). M. Luo was supported by the scholarship from the China Scholarship Council (201706990036). This work of M. Luo was further supported by the Ter Meulen Grant of the Royal Netherlands Academy of Arts and Sciences (KNAW).

#### GINILISA

The authors thank all families for their participation in the studies and the LISA and GINIplus study teams for their excellent work.

#### GLAKU

We thank all the GLAKU children and their parents for their enthusiastic participation. We also thank all the research nurses, research assistants, and laboratory personnel involved in the GLAKU study. The study has been supported by the Academy of Finland, the University of Helsinki, the Hope and Optimism Initiative, the Finnish Foundation for Pediatric Research, the Sigrid Juselius Foundation, the Jalmari and Rauha Ahokas Foundation, the Signe and Ane Gyllenberg Foundation, the Yrjo Jahnsson Foundation, the Juho Vainio Foundation, the Emil Aaltonen Foundation, and the Ministry of Education and Culture in Finland. The 352 samples were genotyped at the Genotyping and Sequencing Core Facility of the Estonian Genome Centre, University of Tartu. J. Lahti has received research support from the Strategic Research Council (SRC) established within the Academy of Finland (decision no. 352700).

#### HRS

The authors would like to thank all the participants, the Health and Retirement Study team from the University of Michigan and the Database of Genotypes and Phenotypes (dbGaP) for providing the data (dbGaP Study Accession no. phs000428.v2.p2). The Health and Retirement Study genetic data is sponsored by the National Institute on Ageing (grant no. U01AG009740, RC2AG036495, and RC4AG039029) and was conducted by the University of Michigan. Data use was approved by the National Centre for Biotechnology Information Genotypes and Phenotypes Database (NCBI dbGaP) Data Access Request system at the National Institutes of Health (Project ID 26700).

##### INMA_GSA

The INMA project was funded by grants from the Instituto de Salud Carlos III (Red INMA G03/176 and CB06/02/0041). The INMA-Gipuzkoa subcohort was funded by the Spanish Biomedical Research Center in Epidemiology and Public Health (CIBERESP) and by grants from the Instituto de Salud Carlos III (PI06/0867, PI09/00090, PI13/02187, PI18/01142, PI21/01491) co-funded by the ERDF, “A way to make Europe”, the Basque Department of Health (Projects GV-SAN2005111093, GV-SAN2009111069, GV-SAN2013111089, GV-SAN2015111065 and GV-SAN2018111086, GV-SAN2019111085), by the Provincial Government of Gipuzkoa (Projects DFG06/002, DFG08/001 and DFG15/221), and by annual agreements with the municipalities of the study area (Zumarraga, Urretxu, Legazpi, Azkoitia, Azpeitia, and Beasain).

##### INMA_Omni1

The authors would like to particularly thank all the participants of the INMA project for their generous collaboration. The INMA project was funded by grants from the Instituto de Salud Carlos III (Red INMA G03/176 and CB06/02/0041). The INMA-Sabadell cohort received funding from the Instituto de Salud Carlos III (FIS-FEDER: PI041436 and PI081151), the Generalitat de Catalunya-CIRIT (1999SGR 00241), and the EU sixth framework project (NEWGENERIS FP6-2003-Food-3-A-016320). The INMA-Valencia cohort received funding from the EU (FP7-ENV-2011 cod 282957 and HEALTH.2010.2.4.5-1), and from the Instituto de Salud Carlos III (FIS-FEDER:

03/1615, 04/1509, 04/1112, 04/1931, 05/1079, 05/1052, 06/1213, 07/0314, 09/02647, 11/0178, 11/01007, 11/02591, 11/02038, 13/1944, 13/2032, 14/00891, 14/01687).

#### INSchool

We are grateful to all the families and schools who kindly participated in the study. This work was funded by the Instituto de Salud Carlos III (PI20/00041, PI22/00464, PI23/00026 and PI23/00404), and co-financed by the European Regional Development Fund (ERDF), the Agència de Gestió d’Ajuts Universitaris i de Recerca (AGAUR, 2017SGR1461 and 2021SGR-00840), the Health Research and Innovation Strategy Plan (PERIS SLT006/17/285 and PERIS SLT006/17/287), the Fundació Marató de TV3 (202228-30 and 202228-31), the Fundació Bancària ”La Caixa”, and the Diputacions de Barcelona i Lleida. S.A. acknowledges her Miguel Servet contract (CP22/00026) awarded by the Instituto de Salud Carlos III and co-funded by the European Union Fund (Fondo Social Europeo Plus, FSE+).

#### LSAC

We thank the Longitudinal Study of Australian Children (LSAC) and Child Health CheckPoint study participants. Data used in this study were collected in partnership with the Department of Social Services (DSS), the Australian Institute of Family Studies (AIFS), the Australian Bureau of Statistics (ABS), and the Murdoch Children’s Research Institute (MCRI). The findings and views reported in this paper are those of the authors and should not be attributed to DSS, AIFS, ABS, or MCRI. The Child Health CheckPoint project was supported by the National Health and Medical Research Council (NHMRC) of Australia (Project Grants 1041352, 1109355, 1128516; Program grant 633003), The Royal Children’s Hospital Foundation (2014-241), the Murdoch Children’s Research Institute (MCRI), The University of Melbourne, the National Heart Foundation of Australia (100660), Financial Markets Foundation for Children (2014-055, 2016-310), the Victoria Deaf Education Institute, the University of Auckland Faculty Development Research Fund

(3712987), Cure Kids (7004), New Zealand Ministry of Business, Innovation and Employment (PROP-53087-INTCS-UOA), and the National Centre for Longitudinal Studies.

#### MCS

We are grateful to the Centre for Longitudinal Studies (CLS), UCL Social Research Institute, for the use of the Millennium Cohort Study data and to the UK Data Service for making them available. However, neither CLS nor the UK Data Service bear any responsibility for the analysis or interpretation of these data.

#### MoBa

This study includes data from the Norwegian Mother, Father and Child Cohort Study (MoBa) conducted by the Norwegian Institute of Public Health. We are grateful to all the participating families in Norway who take part in this ongoing cohort study. We thank the Norwegian Institute of Public Health (NIPH) for generating high-quality genomic data. MoBa is supported by the Norwegian Ministry of Health and Care Services and the Ministry of Education and Research. This research is part of the HARVEST collaboration, supported by the Research Council of Norway (#229624). We also thank the NORMENT Centre for providing genotype data, funded by the Research Council of Norway (#223273), South East Norway Health Authorities and Stiftelsen Kristian Gerhard Jebsen. We further thank the Center for Diabetes Research, the University of Bergen for providing genotype data and performing quality control and imputation of the data funded by the ERC AdG project SELECTionPREDISPOSED, Stiftelsen Kristian Gerhard Jebsen, Trond Mohn Foundation, the Research Council of Norway, the Novo Nordisk Foundation, the University of Bergen, and the Western Norway Health Authorities. AH was supported by funding from the HorizonEurope Research and Innovation Programme (FAMILY, grant agreement no. 101057529; Marie Skłodowska-Curie grant agreement [ESSGN 101073237] ), the Research Council of Norway (RCN) (#274611, #336085) and the South-Eastern Norway Regional Health

Authority (HSØ) (#2020022 & #2018059). E.C.C. was supported by funding from the RCN (#274611) and HSØ (#2021045) and is a member of the MRC Integrative Epidemiology Unit at the University of Bristol which is supported by the Medical Research Council and the University of Bristol (MC_UU_00032/1). T.R.K. was supported by funding from the RCN (#274611). E.Y. was supported by funding from the RCN (#288083 and #336078) and the European Union (Grant agreement No. 101045526 and No. 818425). The funders had no role in study design, data collection and analysis, decision to publish, or manuscript preparation. This publication is the work of the authors, who serve as the guarantors for the contents of this paper. Views and opinions expressed are those of the authors only and do not necessarily reflect those of the European Union or the European Research Council Executive Agency. Neither the European Union nor the granting authority can be held responsible for them. This work was performed on the TSD (Tjeneste for Sensitive Data) facilities, owned by the University of Oslo, operated and developed by the TSD service group at the University of Oslo, IT Department (USIT). The computations were performed on resources provided by Sigma2 - the National Infrastructure for High-Performance Computing and Data Storage in Norway.

**MUSP**

This study was funded by grants received from the National Health and Medical Research Council (NHMRC) and Australian Research Council (ARC). Thanks also to Shelby Marrington (Project Manager) and Greg Shuttlewood (Data Manager), who have supervised the day-to-day management of the study. We also extend our thanks to the mothers and children who have continued to participate in the study.

**NFBC1986**

We wish to thank all cohort members, researchers and NFBC project center personnel who participated in the NFBC data collections. The NFBC1986 cohort was supported by EU QLG1-

CT-2000-01643 (EUROBLCS) Grant no. E51560, NorFA Grant no. 731, 20056, 30167, USA / NIH 2000 G DF682 Grant no. 50945. NFBC1986 33-35y follow-up study received financial support from the University of Oulu (Strategic funding from donations) and Oulu University Hospital (K65760). Sylvain Sebert has received funding from the European Union's Horizon 2020 research and innovation program under grant agreement no. 733206 (LifeCycle), grant agreement no. 848158 (EarlyCause), grant agreement no. 873749 (LongITools) and Academy of Finland project 312123. Priyanka Choudhary has received funding from the University of Oulu and the Academy of Finland Profi6 (decision number AF 336449).

##### NTR

NTR warmly thanks all participating twin families. We acknowledge funding from the Netherlands Organization for Scientific Research (NWO): The Netherlands Twin Register infrastructure was supported by grant NWO 480-15-001/674: Netherlands Twin Registry Repository: researching the interplay between genome and environment, and by multiple grants from the Netherlands Organization for Scientific Research (NWO) that enabled data collections. We gratefully acknowledge funding by CID (Consortium on Individual Development) through the Gravitation Program of the ‘Dutch Ministry of Education, Culture, and Science’ and the ‘Dutch Research Council’ (NWO; grant number 024-001-003). Genotyping was made possible by grants from NWO/SPI 56-464-14192, Genetic Association Information Network (GAIN) of the Foundation for the National Institutes of Health, Biobanking and Biomolecular Research Infrastructure (BBMRI–NL, 184.033.111) and the BBMRI-NL funded BIOS Consortium (NWO184.021.007); and Aggression in Children: Unravelling gene-environment interplay to inform Treatment and InterventiON strategies project (ACTION; European Union Seventh Framework Program (FP7/2007-2013) under grant agreement no 602768); Rutgers University Cell and DNA Repository (NIMH U24 MH 068457-06), the Avera Institute, Sioux Falls (USA) and the National Institutes of Health (NIH R01 HD042157-01A1, MH081802, Grand Opportunity grants 1RC2

MH089951 and 1RC2 MH089995) and European Research Council (ERC-230374). D.I.B. acknowledges the Royal Netherlands Academy of Science Professor Award (PAH/6635).

##### The Raine Study

The authors gratefully acknowledge the NHMRC for their long-term funding to the study over the last 30 years and also the following institutes for providing funding for Core Management of the Raine Study: The University of Western Australia (UWA), Curtin University, Women and Infants Research Foundation, Telethon Kids Institute, Edith Cowan University, Murdoch University, The University of Notre Dame Australia and The Raine Medical Research Foundation. This work was supported by resources provided by the Pawsey Supercomputing Centre with funding from the Australian Government and the Government of Western Australia. The Raine Study was supported by the National Health and Medical Research Council of Australia [Grant Numbers 572613, 403981], the Canadian Institutes of Health Research [Grant Number MOP-82893].

##### SYS

The Saguenay Youth Study was supported by the Canadian Institutes of Health Research. We would like to thank the participants of the Saguenay Youth Study cohort and their families for their contributions.

##### TEDS

We gratefully acknowledge the ongoing contribution of the participants in the Twins Early Development Study (TEDS) and their families. TEDS is supported by a programme Grant to R.P. from the UK Medical Research Council (Grant no. MR/V012878/1, previously no. MR/M021475/1). K.R. is supported by a Sir Henry Wellcome Postdoctoral Fellowship.

##### TRAILS

Participating centres of TRAILS include the University Medical Centre and University of Groningen, the University of Utrecht, and the Parnassia Psychiatric Institute, all in the Netherlands. TRAILS has been financially supported by various grants from the Netherlands Organization for Scientific Research NWO (Medical Research Council program grant GB-MW 940-38-011; ZonMW Brainpower grant 100-001-004; ZonMw Risk Behavior and Dependence grant 60-60600-97-118; ZonMw Culture and Health grant 261-98-710; Social Sciences Council medium-sized investment grants GB-MaGW 480-01-006 and GB-MaGW 480-07-001; Social Sciences Council project grants GB-MaGW 452-04-314 and GB-MaGW 452-06-004; ZonMw Longitudinal Cohort Research on Early Detection and Treatment in Mental Health Care grant 636340002; NWO large-sized investment grant 175.010.2003.005; NWO Longitudinal Survey and Panel Funding 481-08-013 and 481-11-001; NWO Vici 016.130.002, 453-16-007/2735, and Vi.C.191.021; NWO Gravitation 024.001.003), the Dutch Ministry of Justice (WODC), the European Science Foundation (EuroSTRESS project FP-006), the European Research Council (ERC-2017-STG-757364 and ERC-CoG-2015-681466), Biobanking and Biomolecular Resources Research Infrastructure BBMRI-NL (CP 32), the Gratama foundation, the Jan Dekker foundation, the participating universities, and Accare Centre for Child and Adolescent Psychiatry. Statistical analyses were carried out on the Genetic Cluster Computer (http://www.geneticcluster.org), which is financially supported by the Netherlands Scientific Organisation (NWO 480-05-003) along with a supplement from the Dutch Brain Foundation. We are grateful to everyone who participated in this research or worked on this project to make it possible.
